# Assessing the impact of autologous neutralizing antibodies on viral rebound in postnatally SHIV-infected ART-treated infant rhesus macaques

**DOI:** 10.1101/2023.07.22.550159

**Authors:** Ellie Mainou, Stella J Berendam, Veronica Obregon-Perko, Emilie A Uffman, Caroline T Phan, George M Shaw, Katherine J Bar, Mithra R Kumar, Emily J Fray, Janet M Siliciano, Robert F Siliciano, Guido Silvestri, Sallie R Permar, Genevieve G Fouda, Janice McCarthy, Ann Chahroudi, Jessica M Conway, Cliburn Chan

**Author notes:** These authors have contributed equally (Jessica M Conway, Cliburn Chan).

## Abstract

While the benefits of early antiretroviral therapy (ART) initiation in perinatally infected infants are well documented, early ART initiation is not always possible in postnatal pediatric HIV infections, which account for the majority of pediatric HIV cases worldwide. The timing of onset of ART initiation is likely to affect the size of the latent viral reservoir established, as well as the development of adaptive immune responses, such as the generation of neutralizing antibody responses against the virus. How these parameters impact the ability of infants to control viremia and the time to viral rebound after ART interruption is unclear. To gain insight into the dynamics, we utilized mathematical models to investigate the effect of time of ART initiation via latent reservoir size and autologous virus neutralizing antibody responses in delaying viral rebound when treatment is interrupted. We used an infant nonhuman primate Simian/Human Immunodeficiency Virus (SHIV) infection model that mimics breast milk HIV transmission in human infants. Infant Rhesus macaques (RMs) were orally challenged with SHIV.C.CH505 375H dCT and either given ART at 4-7 days post-infection (early ART condition), at 2 weeks post-infection (intermediate ART condition), or at 8 weeks post-infection (late ART condition). These infants were then monitored for up to 60 months post-infection with serial viral load and immune measurements. We develop a stochastic mathematical model to investigate the joint effect of latent reservoir size, the autologous neutralizing antibody potency, and CD4+ T cell levels on the time to viral rebound and control of post-rebound viral loads. We find that the latent reservoir size is an important determinant in explaining time to viral rebound by affecting the growth rate of the virus. The presence of neutralizing antibodies also can delay rebound, but we find this effect for high potency antibody responses only.

## 1 Introduction

In 2020, an estimated 1.7 million children were living with HIV-1 infection worldwide [1]. Infants who acquire HIV must start ART as soon as possible after diagnosis and remain on lifelong ART to prevent HIV-associated disease [2]. The benefits of initiating ART soon after infection are well-documented [3–5]: early ART reduces mortality and improves clinical outcomes in infants living with HIV-1 [3, 5]. Studies have demonstrated that early ART initiation suppresses HIV viral replication and can result in preservation of CD4+ T cell counts both in infants and adults [3,4,6,7]. However, ART is not a cure, and treatment interruption leads to rebound of viremia to levels typical of chronic infection, with remarkable heterogeneity in rebound times [8]. Yet early ART may delay viral rebound after treatment interruption also in infants. For instance, the “Mississippi baby” was treated 30 hours after birth and discontinued at 18 months of age, and no detectable viremia was observed for 28 months after treatment discontinuation [9, 10]. This case study sparked hopes for functional cure of pediatric HIV, i.e. sustained suppression of viremia without ART, through very early administration of ART. Additional reports of potential pediatric HIV-1 remission, through early ART followed [11, 12], but were not attributed to a specific mechanism (cellular or humoral immunity). Despite such promising cases, a generalizable approach for a functional cure in infants and children has not been achieved [5, 13, 14].

In addition to infants, several studies demonstrated the effects of early ART in adults living with HIV. In a pooled analysis of study participants in six AIDS Clinical Trial Groups (ACTG), Li et al. reported widely varying rebound times after treatment interruption, ranging from a few days to months, with a significant number of participants (15 out of 235) maintaining viral loads below the detectable limit for up to 3 or more months after ART interruption [8]. The most comprehensive description of post-treatment control (PTC) was provided by the VISCONTI cohort, a group of 14 people living with HIV and treated early who were able to control viremia for up to 10 years after treatment interruption [15]. However, unlike elite controllers, who have HLA alleles favorable to HIV control, participants in the VISCONTI study did not show overrepresentation in those alleles [8, 15]. In fact, the VISCONTI cohort displayed less efficient HIV-1-specific CD8+ T cells [15]. The development of humoral immune responses was not examined in this cohort [15]. However, *in vitro* studies point to the effect of humoral responses and the functions of antibodies. For example, *in vitro* data suggests that an anti-CD2 monoclonal antibody can induce natural killer cell-driven antibody-dependent cell-mediated cytotoxicity (ADCC). Specifically targeting CD45RA-CD4+ memory T cells, killing can be induced and this can reduce HIV DNA levels in patient samples *ex vivo* [16, 17]. In addition, novel antibody-based therapies against HIV are advancing into the clinic [18]. Therefore, it is crucial to further study immune mechanisms such as the mechanisms of antibody development in postnatally HIV-infected children, and their capacity to control infection. This understanding could potentially lead to more effective and generalized approaches for functional cure of pediatric HIV and provide a better quality of life for these children.

The major hurdle to HIV cure is the establishment of the latent reservoir (LR), which consists of latently HIV-infected cells, i.e. cells with integrated HIV provirus. These latently infected cells are primarily resting memory CD4+ T cells [19]. It is a long-lived population of cells that is established early in infection, and escapes both ART and immune responses. Latently infected cells can “activate” by transitioning to a productive phenotype, which is thought to cause viral rebound if therapy is interrupted [20–26]. Early ART initiation is associated with lower latent reservoir sizes both in adults and infants [27, 28]. However, characterization of the latent reservoir has been largely focused on adults. In infant nonhuman primate models, naive CD4 T cells containing intact provirus— a small fraction of an adult’s latent reservoir [29] — significantly contribute to the size of the viral reservoirs in both blood and lymphoid tissues. This is noteworthy, as memory CD4+ T cells are typically considered the primary source of latent HIV and simian immunodeficiency virus (SIV) infection in adult humans and rhesus macaques [30]. By characterizing the profile of the latent viral reservoir, we can expand our knowledge of viral persistence and gain a clearer understanding of the targets for interventions in the pediatric population. [30].

There have been a number of studies that use mathematical models to investigate control and time to viral rebound after treatment interruption [31–37], but such studies are limited to adults. All these models make the well-accepted assumption that latent cell activation drives HIV-1 viral rebound. Pinkevych et al. used data from treatment interruption trials to derive the first estimates of recrudescence rates [33, 34]. Hill et al. considered inter-individual heterogeneity when modeling within-host viral rebound dynamics and used a continuous time branching process to derive estimates of viral rebound time distributions. Hill et al. assumed that viral growth after viral recrudescence is, on average, exponential, but the validity of this assumption remains unclear [31, 32]. Conway et al. used a stochastic model that incorporates study participant information on HIV reservoir size and pre-ATI drug regimen to refine previous estimates of viral recrudescence after ART cessation [36]; by incorporating this information, they were able to make individual predictions instead of focusing on population averages. These models have been very important in examining the effect of LR size on rebound times [33, 34, 36, 37] and the reduction of the latent reservoir needed to be derived by targeted therapeutics to achieve cure [31, 32]. However, these models neglect the potential effectiveness of the adaptive immune responses to control viremia, once latent cell activation occurs in the absence of ART. Considering the unique biological, immunological, and physiological differences between children and adults with HIV, it is crucial to understand the impact of these factors on viral rebound dynamics in pediatric populations in order to develop age-appropriate treatments and interventions.

In this present study, we focus on viral rebound in an infant nonhuman primate Simian/Human Immunodeficiency Virus (SHIV) infection model that mimics breast milk HIV transmission in human infants. Infant rhesus macaques (RMs) were orally challenged with SHIV.C.CH505 375H dCT 4 weeks after birth and ART was initiated late at 8 weeks post-infection (wpi), intermediately at 2 wpi and early at 4-7 days post-infection (dpi). In addition to regular viral load measurements, longitudinal neutralizing antibody responses as well as the latent reservoir size was assessed after ART interruption [30, 38, 39]. We investigate how the timing of treatment initiation affects the establishment of the latent reservoir and the development of neutralizing antibodies, and how these, in turn, affect viral rebound times. We extend the continuous time branching process model developed by Conway et al. [36] to account for the effect of neutralizing antibodies and latent reservoir sizes. The model output is a cumulative probability distribution that provides the probability of an individual’s viral rebound time by time t. We show that the major determinant for rebound time is the size of the latent reservoir, whereas neutralizing antibodies can delay rebound only when high neutralization potency is observed.

## 2 Methods

### 2.1 The data

#### 2.1.1 The experiments

Ten infant *Rhesus macaques* (RMs) were orally challenged with SHIV.C.CH505 four weeks after birth. Antiretroviral treatment (ART) was initiated within 8 weeks post-infection (wpi) and subjects were followed for viral rebound, defined as sustained detectable viremia. The experiment was replicated twice with different treatment initiation times: ten animals started treatment at 2 wpi (intermediate treatment group) and another 10 started 4-7 days post-infection (dpi) (early treatment group). The experiments are described in detail in [30, 38, 39].

We calibrate our model using the following experimental measurements: 1) viral load measurements, 2) potency of antibodies neutralizing autologous virus measured by a TZM-bl assay [40] and 3) the latent reservoir size assessed with Intact Proviral DNA assay (IPDA) [41].

#### 2.1.2 Estimation of CD4+ T cell population

While IPDA is measured as intact genomes per 10^6^ CD4+ T cells, our branching process model requires the latent reservoir size for the entire monkey, i.e. we need CD4+ T cell measurements for our subjects. Because CD4+ T cells were only measured for the late group, we estimated the latent reservoir size for the intermediate and early treatment groups by averaging the measure over all possible values of CD4 T+ cells. This requires knowledge of the distribution of CD4 T+ cells in infant macaques. We fit lognormal, normal, and Weibull distributions to the late treatment group’s CD4+ T cell measurements (Fig. 8A) and using Akaike Information Criterion (AIC) we determined that the lognormal distribution, with *logN*(19.33, 0.53) best explains the CD4+ T cell counts across the test subjects (Table 4).

#### 2.1.3 Estimation of antibody neutralization efficacy of the undiluted sample

Antibody neutralization against the challenge virus is measured in a serial dilution of plasma samples, starting at 1:20. To inform our model of antibody efficacy, we need to estimate the antibody percent neutralization of the undiluted sample. To estimate that, we fit a 4-parameter logistic model to the neutralization percent data at different dilutions. The model takes the form of

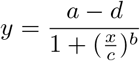

where *x* is the dilution, *y* is the percent neutralization and *a, b, c, d* are model parameters. From the fitted model we extrapolate the percent antibody neutralization of the undiluted plasma (Figure 8B for an example fit, Figure 5 for all fitted curves in the late treatment group). We define a strong neutralizing response as one where the percent neutralization of the undiluted plasma is *≥* 80%.

### 2.2 The model

Our baseline model is described in [36, 37]. In short, we assume that latent cell activation drives viral rebound [36], but not all activations cause rebound. Instead, a latent cell activation is followed by rounds of viral replication that may cause viral population sizes to grow past the detection threshold or to die out (Figure 1). We define *q* as the probability that a latent cell activation fails to cause viral rebound, so 1 *− q* is the probability that a latent cell activation will lead to viral rebound. Finally, we assume that there is a time delay associated with latent cell activation dynamics, and viral growth, between successful latent activation and detectable viremia. The detection threshold is defined as 60 HIV RNA copies/ml. For simplicity, we assume a fixed, delta-distributed delay [36, 37].

**Figure 1:**
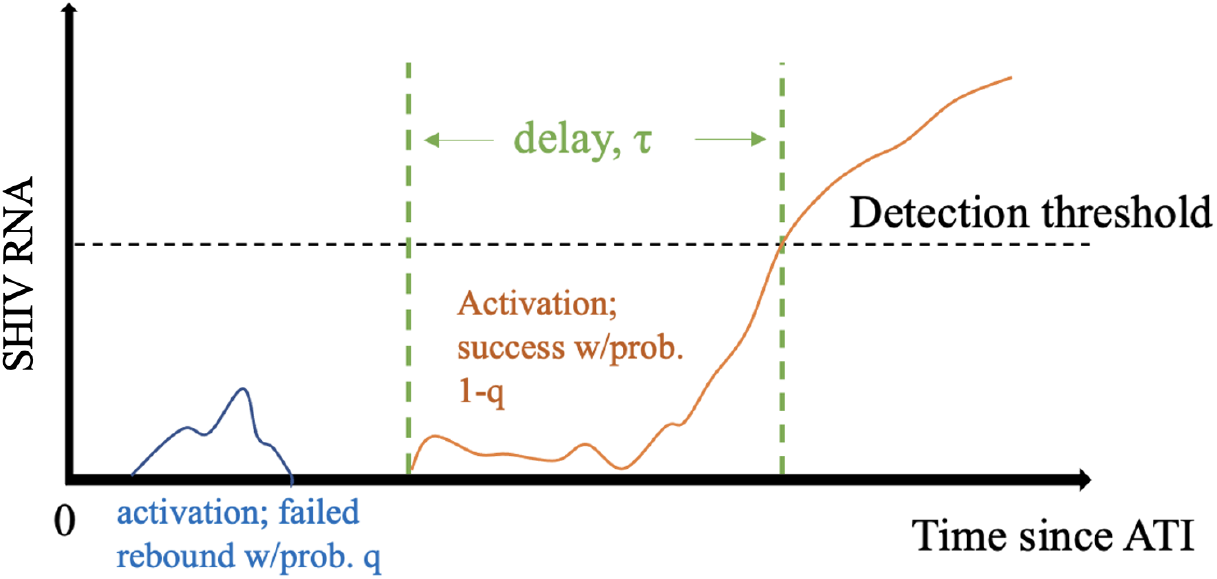
Model schematic. We assume that following ATI, latent cell activations are followed by chains of infection that may die out, i.e., go extinct, with probability q, or successfully re-establish high viral loads associated with chronic infection, with probability 1*−* q. In the latter case, we further assume a delay τ between activation and the time when plasma viral load crosses the detection threshold.

We take the latent reservoir at the time of ATI to be *L*_0_ and the *per capita* latent cell activation rate *a*. Hence, the recrudescence rate is *r* = (1 *− q*)*aL*_0_. We use a branching process to compute the probability of viral rebound at time *t*. The cumulative probability of successful activation at time t is 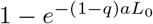. Assuming a fixed delay *τ*, the model cumulative probability of viral rebound by time *t* is

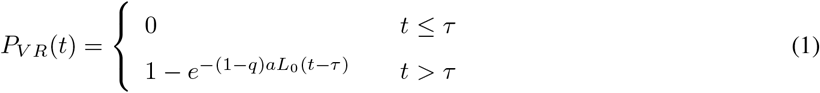

We extend our baseline model to account for the effect of neutralizing antibodies and the size of the latent reservoir as estimated from IPDA. In particular, we assume that antibodies may affect the probability of unsuccessful activation *q* and/or the delay *τ*. Additionally, we assume that the latent reservoir size may also affect the delay between ATI and rebound. We construct and test several simple functions to express *q* and *τ* as functions of neutralizing antibodies and latent reservoir size (Table 1). For example, we use the Hill function, 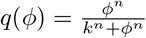 with *n* = 1, 2, 3, to express the effect of antibody neutralization *ϕ* on the probability of unsuccessful activation *q*, estimating k.

**Table 1:** List of expressions of the probability of unsuccessful activation q and the time delay, τ as functions of antibody responses, ϕ and latent reservoir size L_0_, which were tested in our model.

| $q(\phi, L_0)$ | $\tau(\phi, L_0)$ |
| --- | --- |
| Linear functions of $\phi$ | Linear functions of $\phi$ |
| 1. $q(\phi) = q_0\phi$ | 1. $\tau(\phi) = \tau_1 + (\tau_2 - \tau_1)\phi$ |
| 2. $q(\phi) = q_0 + q_1\phi$ | 2. $\tau(\phi) = \tau_1 + \tau_2\phi$ |
| Hill function of $\phi$ | Hill function of $\phi$ |
| 3. $q(\phi) = \frac{\phi^n}{k^n + \phi^n}, n = 1, 2, 3$ | 3. $\tau(\phi) = \tau_1 + \tau_2 \frac{\phi^n}{k^n + \phi^n}, n = 1, 2, 3$ |
| | Nonlinear function of $\phi$ |
| | 4. $\tau(\phi) = \tau_1 + (\tau_2 - \tau_1)\phi^2$ |
| | Linear function of $L_0$ |
| | 5. $\tau(L_0) = \tau_1 - \tau_2 L_0$ |
| | Nonlinear function of $L_0$ |
| | 6. $\tau(L_0) = \tau_1 + \frac{\tau_2}{L_0}$ |
| | 7. $\tau(L_0) = \frac{\tau_2}{L_0}$ |
| | Hill function of $L_0$ |
| | 8. $\tau(L_0) = \tau_1 + \tau_2(1 - \frac{L_0^n}{k^n + L_0^n}), n = 1, 2, 3$ |
| | Linear function of $\phi$ and $L_0$ |
| | 9. $\tau(\phi, L_0) = \tau_1 + \tau_2\phi - \tau_3 L_0$ |
| | Nonlinear functions of $\phi$ and $L_0$ |
| | 10. $\tau(\phi, L_0) = \tau_1 + \tau_2 \frac{\phi}{L_0}$ |
| | 11. $\tau(\phi, L_0) = \tau_1 + \tau_2\phi + \tau_3 \frac{1}{L_0} + \tau_4 \frac{\phi}{L_0}$ |
| | 12. $\tau(\phi, L_0) = \tau_1 + \tau_2\phi + \tau_3 \frac{1}{L_0} + \tau_4 \phi L_0$ |

To estimate model parameters, we seek to maximize the likelihood function given by

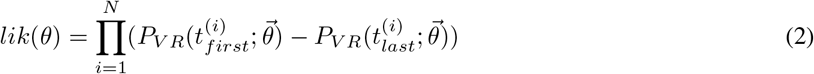

where *P*_*V R*_(*t*) is the cumulative probability of viral rebound by time *t* (eq. 3), multiplied over the *N* study subjects, 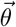 are the parameters and *t*_*last*_ and *t*_*first*_ are the last undetectable and first detectable viral load measurements for the *i*th subject. Note that we do not estimate viral rebound times from the data, using times when viral loads were last undetectable/first dectable directly instead. This allows us to avoid uncertainties due to the extrapolation of rebound times from our dataset. For all models we estimate parameters using maximum likelihood methods, specifically the Davidon-Fletcher-Powell optimization algorithm implemented b the Bhat package in R [42]. We compare model fits using the Akaike Information Criterion (AIC) [43].

We focus on short-term viral rebound of up to 60 days post-ATI, because measurements of antibody neutralization are sparse and only available up to 31 dpi, and therefore the data set might not reflect longer-term dynamics. We calibrate the model using data on 1) viral load measurements of days 0 *−* 60 post-ATI, 2) IPDA measurements at day 0 post-ATI, 3) antibody neutralization efficacy at day 0 post-ATI and 4) CD4+ T cell measurements at day 0 post-infection. We take the latent reservoir size, *L*_0_ to be the *log*_10_ of the IPDA measurement. Our choice for *L*_0_ is motivated by a correlation between rebound time and the *log*_10_ value of the IPDA measurements (Pearson correlation of −0.41, p-value= 0.03).

## 3 Results

We are interested in determining whether different treatment initiation times affect the establishment of the latent reservoir and the development of potent antibody immune responses and how these, in turn, affect rebound times. To investigate this question, we begin by computing our cumulative probability of viral rebound, *P*_*V R*_(*t*) for each treatment group without incorporating any information on the latent reservoir size or the potency of antibodies. We refer to these models as null models. We then incorporate the latent reservoir size and the strength of neutralizing antibody responses against the challenge virus in *P*_*V R*_(*t*) according to the functions listed in Table 1. We compare these models to the null model for each treatment group, as well as all treatment groups combined, to detect a net effect. We finally discuss how our model can be used to examine the effects of potential therapies.

### 3.1 Null models

To facilitate the reader’s comparison with the best model in each treatment group, we provide the parameter estimations for the null model in Table 2. It should be noted that *α* and 1 *− q* appear as a product only in *P*_*V R*_(*t*) and therefore are estimated as one composite parameter. Null model fitted curves to the last undetectable/first detectable viremia for each treatment group are reported to Figure 2.

**Table 2:** Summary of null model parameter estimates (mean and 95% CI) and AIC for each treatment group.

| Group | Parameter Estimates |  | AIC |
| --- | --- | --- | --- |
| | $\alpha(1 - q)$ | $\tau$ | |
| Late | 0.05 (0.02, 0.1) | 4.99 (2.05, 10.52) | 60.99 |
| Intermediate | 0.11 (0.05, 0.21) | 6.07 (4.39, 8.19) | 45.05 |
| Early | 0.012 (0.005, 0.03) | 17 (16.3, 17.7) | 35.2 |
| All groups | 0.03 (0.02, 0.04) | 4.88 (2.52, 8.73) | 157.8 |

**Figure 2:**
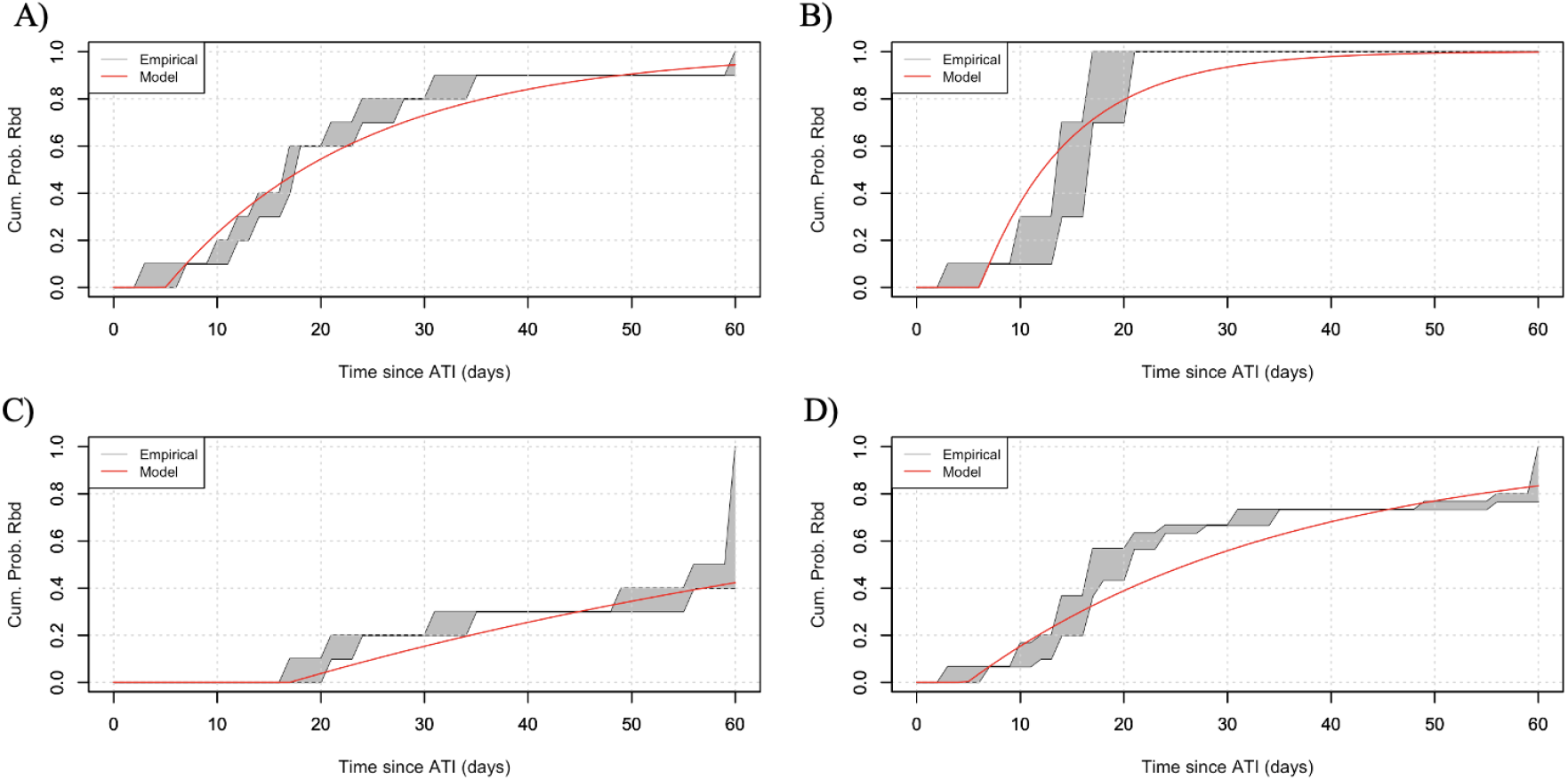
Null model predictions on time to viral rebound for subjects (grey shaded area) and the average time to viral rebound (thick, solid line), for a) the late treatment group, b) the intermediate treatment group, c) the early treatment group and d) all treatment groups combined. For the null model, we do not incorporate information on the latent reservoir size and the strength of neutralizing antibodies on the cumulative probability of rebound, *P*_*V R*_.

### 3.2 Models that best explain the data

#### LR size accelerates time to rebound and neutralizing antibodies delay it in the late treatment group

We fit *P*_*V R*_(*t*) (Eq. 3) to each treatment group separately, and then all treatment groups combined to discern a “net effect”. We find that the model that best explains data for the late treatment group is the one where the probability of unsuccessful activation *q* is affected by the antibody response, according to the function *q* = *q*_0_*ϕ*, and the time delay *τ* is affected by both neutralizing antibodies and the LR size, according to the function *τ* = *τ*_1_ + *τ*_2_*ϕ − τ*_3_*L*_0_ (Δ*AIC* = *−*5.9) (Figure 3A, Table 3). This finding suggests that viral rebound in the late treatment group is mediated by both the humoral immune response and the latent reservoir size. Our results suggest that antibodies play a dual role, altering both the outcome of a latent cell activation, and the speed with which plasma viremia crosses the detection threshold. Interestingly, a larger latent reservoir is associated with a shorter delay between activation and detectable viremia, suggesting that after the successful activation occurs, there are subsequent activations that contribute to overall viremia during this stochastic phase, consistent with model predictions for SIV in van Dorp et al. (2020) [44].

**Table 3:**
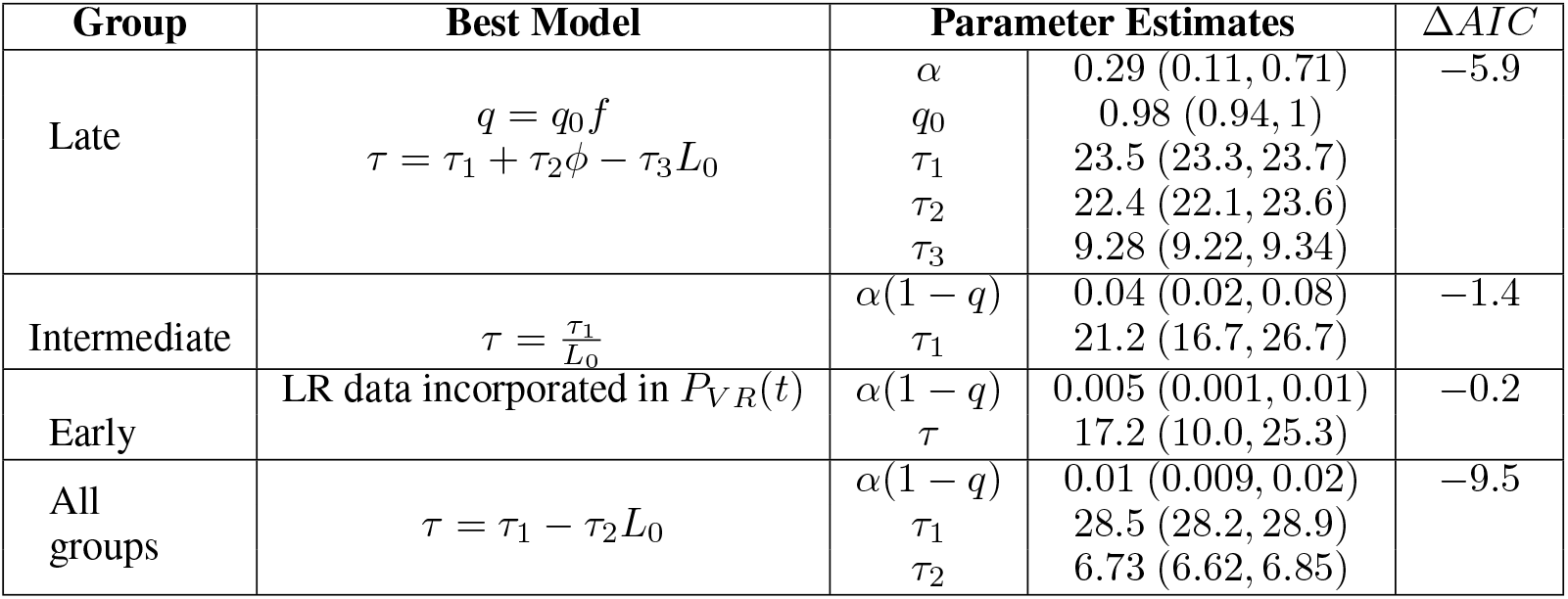
Summary of best models for each treatment group.

**Table 4:** Fitted Distributions to Late Treatment Group’s CD4+ T cell measurements.

| Distribution | Parameters | AIC |
| --- | --- | --- |
| Lognormal | mean=19.33, standard deviation= 0.53 | 406 |
| Normal | mean=$2.9 \times 10^8$, standard deviation=$1.7 \times 10^7$ | 412 |
| Weibull | shape=1.81, scale=$3.48 \times 10^8$ | 484 |

**Figure 3:**
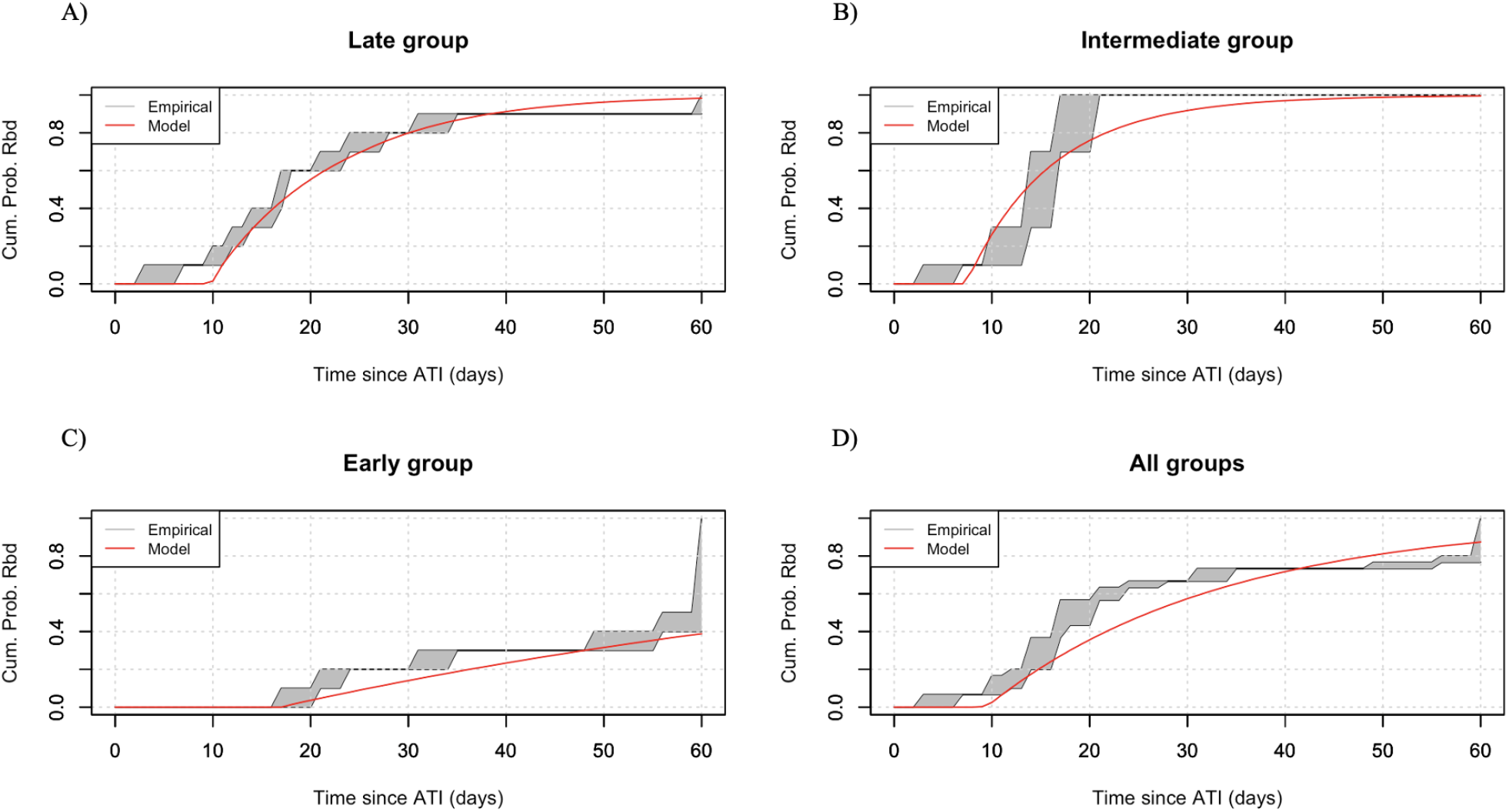
Best model predictions on time to viral rebound for subjects (grey shaded area) and the average time to viral rebound (thick, solid line), for a) the late treatment group, b) the intermediate treatment group, c) the early treatment group and d) all treatment groups combined.

#### LR size explains rebound time in the intermediate treatment group

For the intermediate treatment group, we find that it is the LR size that affects the delay, based on 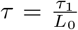. Incorporating an antibody response does not improve fits (Figure 3B, Table 3). However, we only claim that this model has equal statistical support relative to the null model (Δ*AIC* = *−*1.4, we claim statistical significance only for Δ*AIC ≥* 4 [45]). In the intermediate treatment group viral rebound can be explained by the latent reservoir size with at most negligible effects from neutralizing antibody responses. Our observation that a large reservoir size is associated with shorter delays in the late treatment group is also present with the intermediate group. We conclude that antibody responses are not strong enough to affect the dynamics in this intermediate time to ART initiation group.

#### LR size and neutralizing antibodies do not help explain rebound times in the early treatment

The model that best explains the data for the early treatment group is the one where the information on the latent reservoir size is incorporated in the *P*_*V R*_(*t*), but neither the latent reservoir, nor antibody responses affect the probability of unsuccessful activation or the time delay (Figure 3B, Table 3). However, again, this improvement over the null model is not statistically significant (Δ*AIC* = *−*0.2), nor is statistically worse. None of the subjects in the early treatment group shows reservoir sizes detectable by the IPDA assays. Incorporating information on reservoir detection threshold for each subject, improves model fits by AIC, compared to the null model. In addition, the prolonged rebound times observed in the early treatment group can be attributed to the small reservoir size. As with the intermediate treatment group, antibody responses are underdeveloped and do not seem to affect rebound times.

#### LR size is the most important determinant of rebound times across treatment groups

Finally, when we combine all treatment groups, we find that a larger LR size affects the time delay, *τ* = *τ*_1_ *− τ*_2_*L*_0_. Incorporating antibody responses does not improve model fits (Figure 3D, Table 3). This model provides a statistically significant improvement over the null (Δ*AIC* = *−*9.5), which suggests that overall, across all groups, rebound is driven by the latent reservoir size. Even though there are 5 out of 30 subjects that develop a strong antibody response, their effect is diluted by the majority of subjects who develop a weak or undetectable neutralizing response.

### 3.3 Effect of potential interventions

We then use our modeling framework to investigate the effect of potential interventions on viral rebound times. More specifically, we examine by how many days rebound is extended if we increase antibody neutralization or if we decrease latent reservoir size. To investigate this question, we use the best model for the late treatment group as well as the data of the late treatment group on antibody neutralization and latent reservoir size to derive the cumulative probability of rebound for each subject. We increase neutralization by 10% and 50% and we decrease the latent reservoir size by 1 and 2 logs. We re-derive the cumulative probability of rebound for each subject, separately for each case and calculate the median time to rebound for each subject. Again we focus on rebound for up to 60 days post-ATI. If we increase neutralization by 10%, the median time to viral rebound increases by approximately 7 days (Figure 4A, Supplemental Figure 5). If, however, neutralization efficacy is increased by 50%, the median rebound time for only 6 out of the 10 subjects falls beyond the 60-day time frame tested. For the remaining 4 subjects, the increase in median time to viral rebound is 31 days (Figure 4A, Supplemental Figure 5). Similarly, when latent reservoir decreases by 1 log, median time to rebound increases by an average of 12 days. When latent reservoir decreases by 2 logs, median time to rebound increases by an average of 29 days with a median time to rebound of more than 60 days for only two subjects (Figure 4B, Supplemental Figure 6).

**Figure 4:**
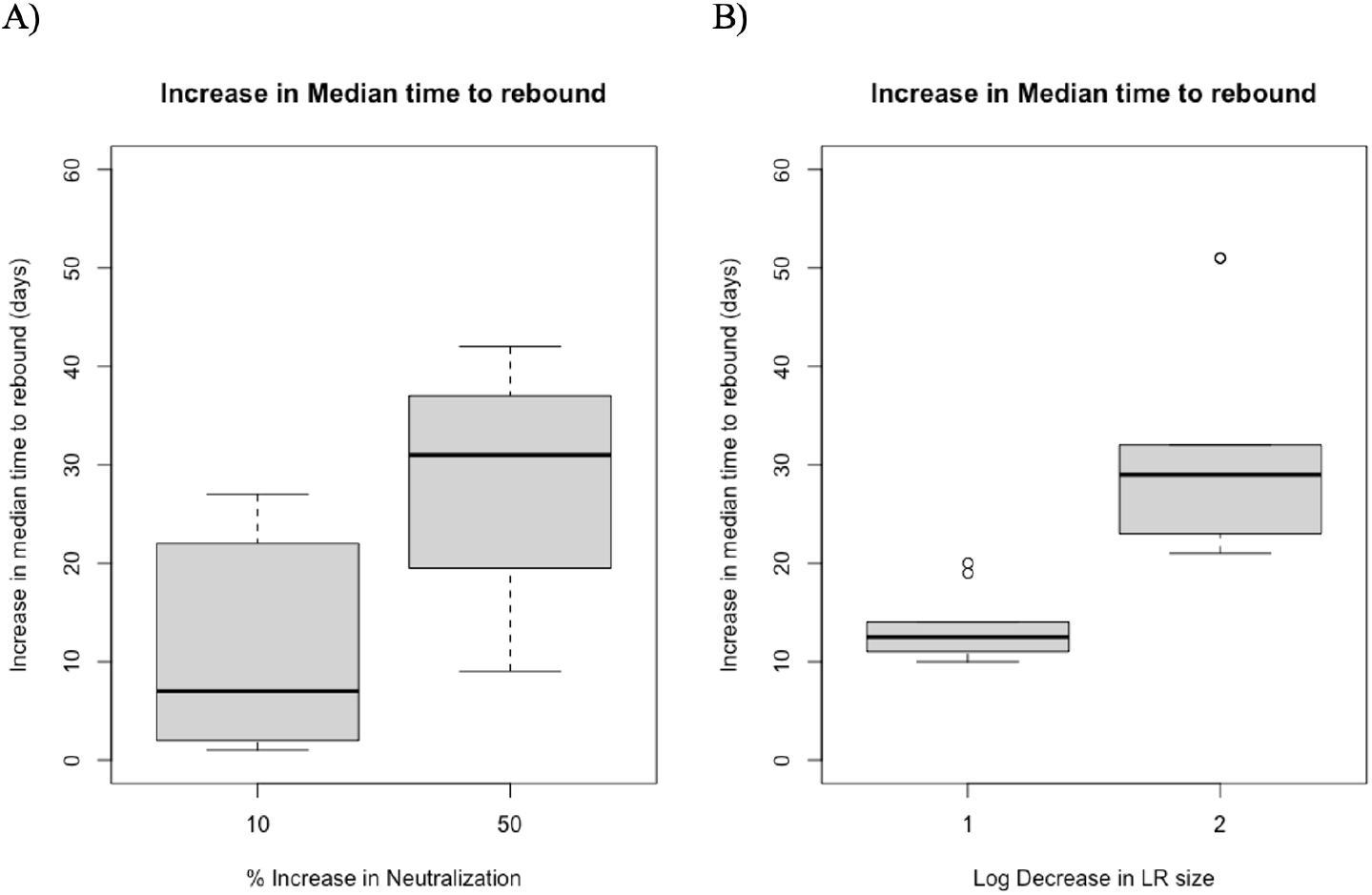
Increase in median time to viral rebound through an intervention that elevates the antibody neutralization strength by 10% and 50% (A) or decreases the latent reservoir size by 1 or 2 logs (B).

## 4 Discussion

In this study, a stochastic model incorporating subject-specific information on the potency of autologous virus neutralizing antibody responses and latent reservoir size was developed. The purpose of this model was to determine the contribution of these factors in explaining rebound times for SHIV-infected infant Rhesus macaques (RMs), a model of HIV breast milk transmission of postnatally infected human pediatric population. We find that the latent reservoir size accelerates time to viral rebound, whereas neutralizing antibodies can delay rebound. The effect of antibody responses on rebound times occurs for the late treatment group only, and not for the intermediate or early groups nor for the combination of all treatment groups. It should be noted that only subjects in the late treatment group developed detectable autologous virus neutralization and half of them developed strong neutralizing responses against autologous challenge virus. This suggests that antibodies against the autologous challenge virus affect viral dynamics post-ATI for high neutralization potencies only. Our results corroborate what is already known in the field in perinatally HIV-infected children and in HIV-infected adults [5, 8, 32, 46]. In In this study in a highly-relevant preclinical model that allows for precise knowledge of the challenge virus and timing of interventions, we focused specifically on investigating the viral rebound times following analytic treatment interruption (ATI). We focus on the first 60 days after ATI due to the sparse data on antibody responses and reservoir sizes. While viral rebound can occur within the first 60 days post-ATI, it is important to note that this does not necessarily signify a failure to control the infection. In some cases, control of the infection can be re-established later on. This is observed particularly in subjects of the early treatment group.

Our modeling results and experimental observations suggest a trade-off between the size of the latent reservoir and the strength of antibody responses. Half of the late-treated animals developed autologous virus neutralizing responses which were associated with prolonged viral control after ATI. Simultaneously, they have large reservoirs, which make it more likely for a successful activation to occur. On the contrary, early treated animals had no neutralizing antibodies to help with control post-ATI, but have small reservoirs, thus prolonging control. Subjects treated at the intermediate time point faced the drawbacks of both groups: treatment initiated too early to develop a neutralizing response, yet not early enough to prevent the seeding of a latent reservoir that could re-establish viremia post-treatment interruption, leading to a median rebound time for the intermediate treatment group that was similar to the late treatment group. The presence of antigen is necessary to stimulate a humoral immune response. Therefore, we believe that therapies that enhance antibody neutralization may be more effective in intermediate-treated individuals. This is because a stronger neutralizing response could potentially counteract the smaller number of latent cell activations that occur due to the smaller reservoir size. However, further research is needed to confirm this hypothesis and to fully understand the complex interplay between antibody responses and the size of the latent reservoir. To test, for example, the effect of agents aiming to increase neutralization or decimate the latent reservoir, this model can be used to predict by how long rebound can be prolonged and the variability in those estimates.

Our model predicts that larger latent reservoir sizes are associated with shorter delay times between successful activation and detectable viremia, which suggests that the latent reservoir has some impact on the net growth rate of viral loads during rebound. We conjecture that this effect may be associated with secondary activations contributing to viral load as in Van Dorp et al. (2020) [44]. This is also supported by experimental observations: for example, experiments where treatment interruptions were conducted on macaques that had been infected with a genetically barcoded SIV strain indicated that a significant number of cells could reactivate successfully from the latent reservoir [35].

These results are in accordance with what previous modeling work suggests about the determinants of viral rebound. Hill et al. [31]) used a stochastic mathematical model to calculate the percent reduction in the latent reservoir size that latency reversing agents need to achieve for ART-free control. They find that a 2000-fold decrease in LR size is needed to delay rebound for one year, suggesting that LR size is an important determinant for time to viral rebound. However, this is not observed universally. Sharaf et al. [47] found only a 7-fold difference in the latent reservoir size (total and intact proviral genomes) between post-treatment controllers and non-controllers, which suggests the size of the latent reservoir cannot be a major factor determining the outcome post ATI for this group of participants. Similarly, Conway et al. [46] explored the effect of the strength of the cellular immune response on viral rebound. Using a post-treatment HIV viral dynamics model and a bifurcation analysis they estimate the strength of the cytotoxic T lymphocyte killing needed to achieve post-treatment or elite control. A strong effector cell response is needed to achieve viral remission. Here, instead of a cellular response, we explore the effects of the antibody response and find that a strong neutralization potency against autologous challenge virus is needed to explain observed rebound times when treated late.

In making our model predictions, we took the latent reservoir to be proportional to the logarithm of the IPDA measurement. We chose this expression because it appears that the order of magnitude is more important than specific numbers. The latent reservoir is comprised primarily of memory cells [48], each one specific for a pathogen. There is evidence that the latent reservoir consists of clonal populations [49–51], which indicates that there may be heterogeneous subsets of latent cells. Here, we neglect the inherent heterogeneity of the latent reservoir in favor of average dynamics, but this assumption can be relaxed. Preliminary work tested non-constant latent cell activation rates [37], but these models performed poorly (results not shown). In addition to potential heterogeneity in latent cell activation, there is evidence for variability in the fraction of viruses in the latent reservoir to be neutralized by autologous IgG [52], indicating that certain viral lineages are more susceptible to autologous neutralzation. Measurement of the frequency of reservoir viruses capable of outgrowth in the presence of autologous IgG might refine predictions rebound times [52].

We also simplified our model by taking a constant latent reservoir size and neglecting long-term dynamics of cell death and proliferation. Prior to viral rebound, we expect that the reservoir size would decay at on-therapy rates [48]. However, the latent reservoir is typically large, and decay is slow. For HIV, it is estimated that the half-life is on average 44 months [24, 53]. In absence of other data, we assume that the reservoir decay rate for SHIV in infant macaques is equally slow, so that in the 60-day period investigated in this study, the effect of reservoir dynamics is minimal.

As pointed out by previous studies, the composition of the latent HIV reservoir in infants may differ from that of adults due to differences in immune system development and the timing of infection. The limited data available on the subject suggests that the majority of the latent reservoir in infants may be composed of naive cells, rather than fully differentiated memory cells as seen in adults. These findings have important implications for the development of strategies to target the latent reservoir in pediatric HIV infection [54, 55].

One potential limitation of this study is the use of an animal model of SHIV infection in infant macaques, rather than HIV infection in human infants. While nonhuman primate models have been used extensively in HIV research and have provided valuable insights into disease pathogenesis [30, 39, 56–60], the translation from macaques to humans and from SHIV to HIV may not be fully accurate. In the case of SHIV, many of the currently available SHIVs have significant limitations and cannot fully replicate the characteristics of primary SIV and HIV strains [61–64]. Even though the SHIV strain used in these experiments, SHIV.C.CH505, can mimic the early viral replication dynamics and pathogenesis of HIV infection in humans and thus can be used as an animal model for HIV pathogenesis [65], it does not necessarily mean that the results from this animal model can be translated to infant humans. Another potential limitation of this model when trying to translate these findings for HIV-infected human infants is the neglect of the effect of breast milk in viral control. In cases when infants are breastfed, the mother’s breast milk may provide protection. Antibodies specific to HIV were detected in the breast milk of lactating mothers with HIV infection [66, 67]. In addition, both colostrum and mature breast milk of HIV-infected women contains secretory IgA, secretory IgM and IgF against HIV antigens [68], as well as innate antiviral proteins that work against HIV [69].

While we acknowledge limitations and simplifying assumptions of our model, this study provides estimates of rebound times while incorporating subject-specific information on the latent reservoir size, as well as on the potency of the neutralizing antibody response against the autologous challenge virus in the context of postnatal HIV infection where ART treatment is usually delayed. Additionally, our results offer insight into the dynamics that shape viral rebound times and this study could be used to inform the design of clinical trials. Such clinical trial include broadly neutralizing antibodies for HIV therapy, as well as passive immunization trials tailored to the pediatric HIV population and their developing immune system.

## 5 Acknowledgements

J.M.C. acknowledges the support of the National Science Foundation (grant no. DMS-1714654) and National Institutes of Health (grant nos. R21-AI143443-01A1 and R01-OD011095). E.M. acknowledges the support of the National Science Foundation (grant no. DMS-1714654). G.M.S acknowledges the support of the National Institutes of Health (grant no. R01 AI160607). C.C. acknowledges the support of the National Institutes of Health (grant no. R25AI140495). S.R.P. acknowledges the support of the National Institutes of Health (grant no. P01-AI131276-05).

## 6 Supplementary Text and Figures

### 6.1 Unavailable or below the detection threshold measurements

Intermediate and early treatment groups had neutralization measurements below the detection threshold. Half of the subjects and all of the subjects in the intermediate and early treatment groups respectively had IPDA measurements below the detection threshold. IPDA is measured as intact genomes/ 10^6^ CD4+ T cells. Finally, there are no available CD4+ T cell measurements. In such cases, we take the average probability of viral rebound for values of up to the detection threshold for the type of measurement that is missing. For example, if antibody efficacy is below the detection threshold for a certain subject, the cumulative probability of viral rebound by time *t* becomes

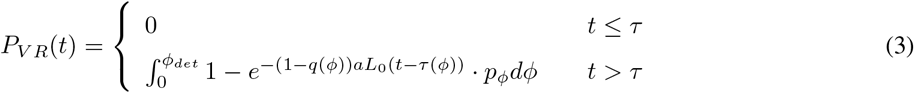

(Eq. 3 in the main text) where *ϕ*_*det*_ = 0.44 is the detection threshold of the TZM-bl assay [40] and *p*_*ϕ*_ is the probability density function of antibody neutralization. If we have two missing measurements for a given subject, for example antibody neutralization and IPDA, then the cumulative probability of viral rebound by time *t* becomes

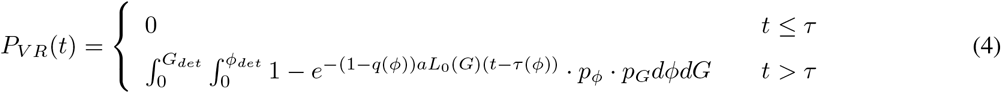

where *G* is the IPDA measurement/ 10^6^ CD4+ T cells, *G*_*det*_ is the detection threshold of the IPDA assay for that subject and *p*_*f*_ and *p*_*G*_ are the probability density functions of antibody neutralization and IPDA measurements respectively.

Considering the lack of data regarding the distribution of antibody neutralization efficacy and IPDA, we assume that these follow a uniform distribution. Therefore, 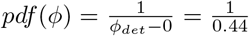[40], and 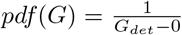, where *G*_*det*_ is the subject-specific IPDA detection threshold.

**Figure 5:**
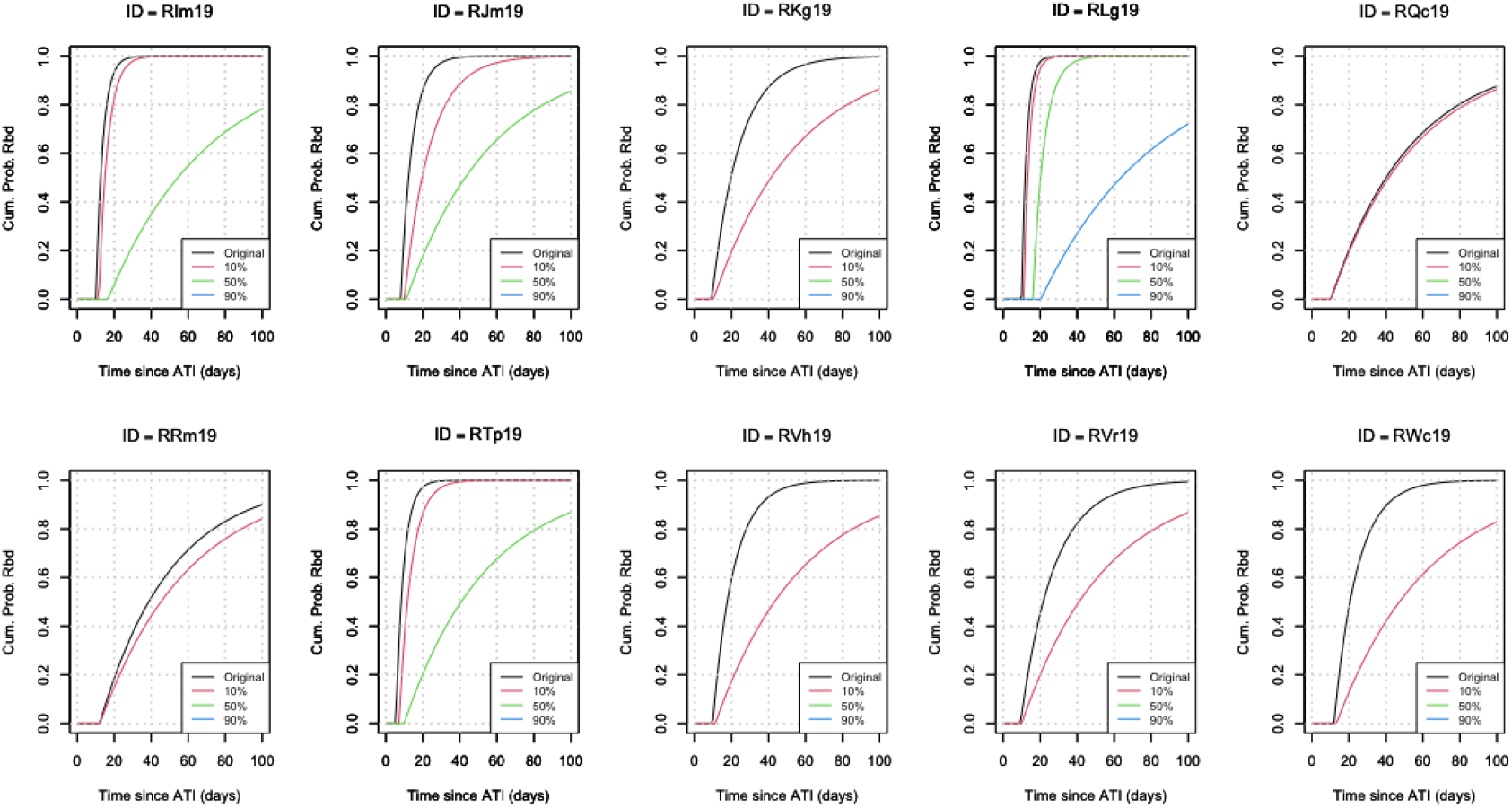
Cumulative Probability of Rebound for each monkey in the late treatment group using the original, data-derived percent neutralization value (black line), a percentage neutralization value increased by 10% (red line), by 50% (green line) and by 90% (blue line). If the increase results in *ϕ >* 1, we do not compute that curve.

**Figure 6:**
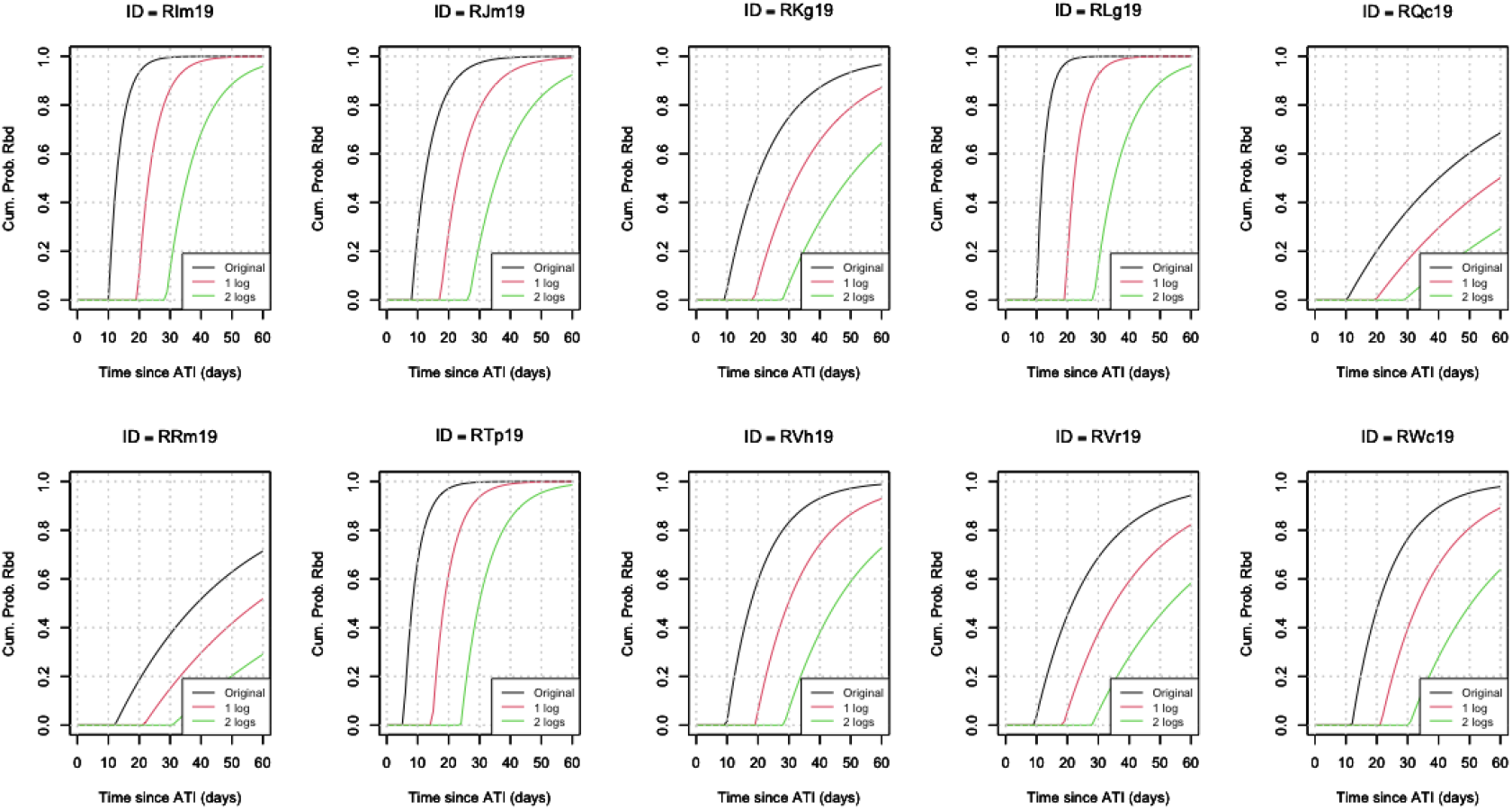
Cumulative Probability of Rebound for each monkey in the late treatment group using the original, data-derived latent reservoir size (black line), a decrease in reservoir size by 1 log (red line), and by 2 logs (green line).

### 6.2 Effect of potential interventions

**Figure 7:**
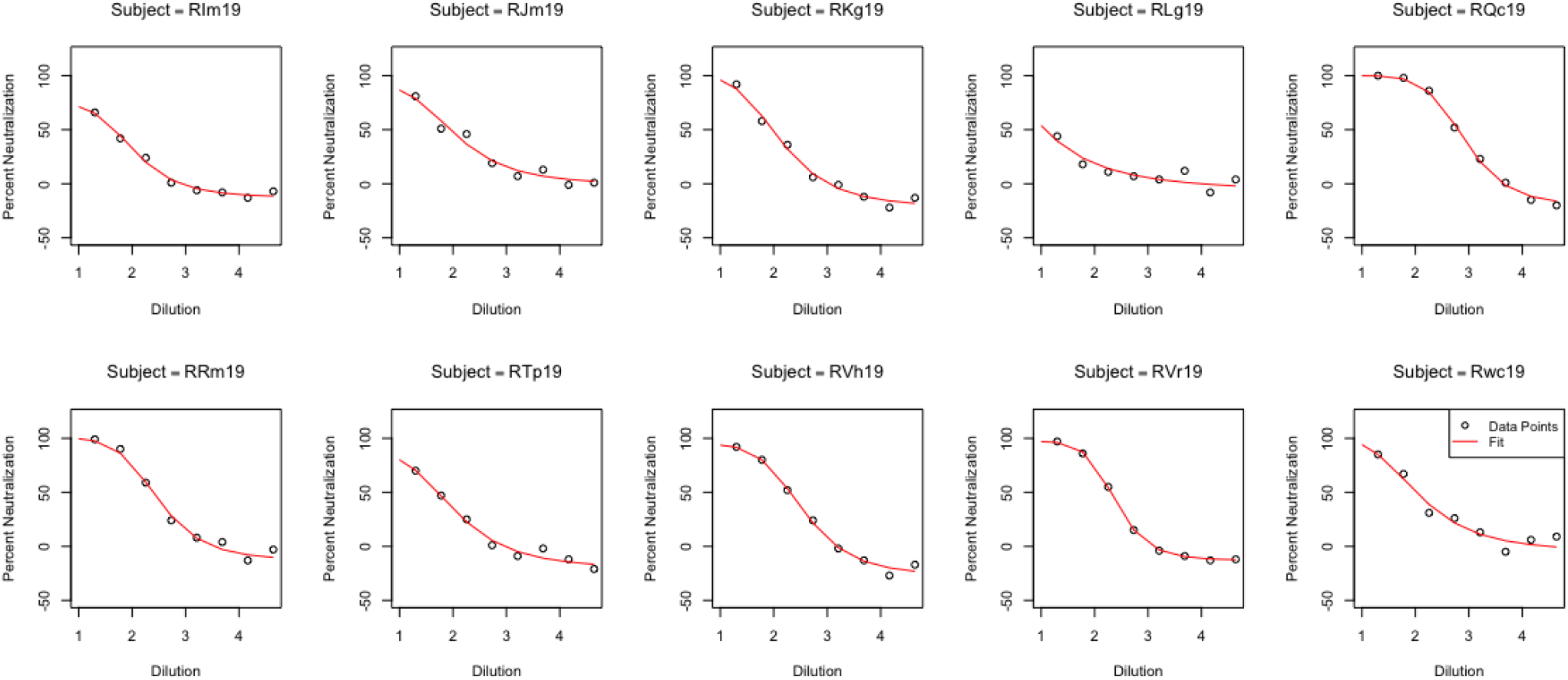
Fitted 4-parameter logistic model (red line) on the percentage neutralization by dilution (black point) for the late treatment group subjects.

**Figure 8:**
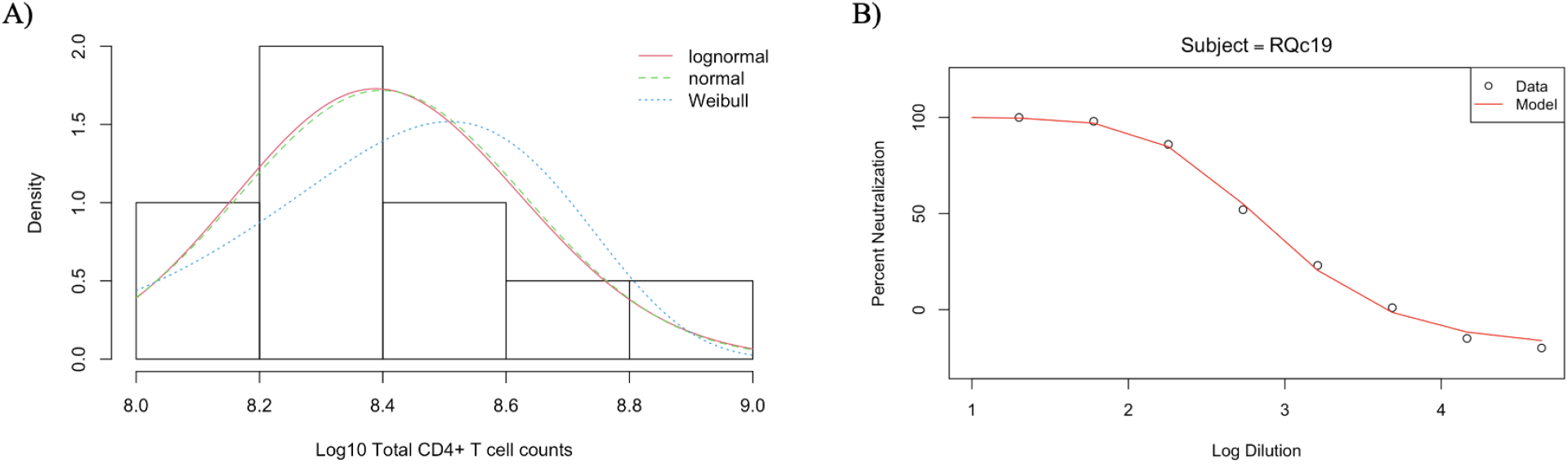
A) Fitted distributions to total CD4+ T cell counts measured in the late treatment group. The distribution that best explains CD4+ T cell data is *logN*(19.22, 0.53) (red line). B) Example of the 4-parameter logistic model fitted curve to the antibody neutralization measurements for the subject RQc19.

## References

[1] UNAIDS. Global HIV & AIDS Statistics — Fact sheet. Available at: http://www.unaids.org/en/resources/fact-sheet, 2021. Accessed July 11 2022.

[2] WHO. Consolidated guidelines on HIV prevention, testing, treatment, service delivery and monitoring: recommendations for a public health approach, 2021 update. 2021. Available at: apps.who.int/iris/handle/10665/342899.

[3] Luzuriaga K. Early combination antiretroviral therapy limits HIV-1 persistence in children. Annu Rev Med, 67:201–13, 2016. doi: 10.1146/annurev-med-091114-111159.

[4] Garcia-Broncano P, Maddali S, Einkauf KB, Jiang C, Gao C, Chevalier J, and et al. Early antiretroviral therapy in neonates with HIV-1 infection restricts viral reservoir size and induces a distinct innate immune profile. Sci Transl Med, 11:eaax7350, 2019. doi: 10.1126/scitranslmed.aax7350.

[5] Berendam SJ, Nelson AN, Yagnik B, Goswami R, Styles TM, Neja MA, Phan CT, Dankwa S, Byrd AU, Garrido C, Amara RR, Chahroudi A, Permar SR, and Fouda GG. Challenges and opportunities of therapies targeting early life immunity for pediatric hiv cure. Front. Immunol, 13:885272, 2022. doi: 10.3389/fimmu.2022.885272.

[6] Shiau S, Abrams EJ, Arpadi SM, and Kuhn L. Early antiretroviral therapy in HIV-infected infants: Can it lead to HIV remission? Lancet HIV, 5:e250–e8, 2018. doi: 10.1016/S2352-3018(18)30012-2.

[7] Cotton MF, Violari A, Otwombe K, Panchia R, Dobbels E, Rabie H, and et al. Early time-limited antiretroviral therapy versus deferred therapy in South African infants infected with HIV: Results from the children with HIV early antiretroviral (CHER) randomised trial. Lancet, 328:1555–63, 2013. doi: 10.1016/S0140-6736(13)61409-9.

[8] Li JZ, Etemad B, Ahmed H, Aga E, Bosch RJ, Mellors JW, Kuritzkes DR, Lederman MM, Para M, and Gandhi RT. The size of the expressed HIV reservoir predicts timing of viral rebound after treatment interruption. AIDS, 30, 2016. doi.org/10.1097/QAD.0000000000000953.

[9] Luzuriaga K, Gay H, Ziemniak C, Sanborn KB, Somasundaran M, Rainwater-Lovett K, Mellors JW, Rosenbloom D, and Persaud D. Viremic relapse after HIV-1 remission in a perinatally infected child. N Engl J Med, 372:786 –8, 2015. doi.org/10.1056/NEJMc1413931.

[10] Persaud D, Gay H, Ziemniak C, Chen YH, Piatak M, Jr, Chun TW, Strain M, Richman D, and Luzuriaga K. Absence of detectable HIV-1 viremia after treatment cessation in an infant. N Engl J Med, 369:1828 –35, 2013. doi.org/10.1056/NEJMoa1302976.

[11] Bitnun A, Samson L, Chun TW, Kakkar F, Brophy J, Murray D, Justement S, Soudeyns H, Ostrowski M, Mujib S, Harrigan PR, Kim J, Sandstrom P, and Read SE. Early initiation of combination antiretroviral therapy in HIV-1-infected newborns can achieve sustained virologic suppression with low frequency of CD4 T cells carrying HIV in peripheral blood. Clin Infect Dis, 59:1012–19, 2014. doi.org/10.1093/cid/ciu432.

[12] Frange P, Faye A, Avettand-Fenoel V, Bellaton E, Descamps D, Angin M, David A, Caillat-Zucman S, Peytavin G, Dollfus C, Le Chenadec J, Warszawski J, Rouzioux C, Saez-Cirion A, and ANRS EP47 VISCONTI Study Group ANRS EPF-CO10 Pediatric Cohort. HIV-1 virological remission lasting more than 12 years after interruption of early antiretroviral therapy in a perinatally infected teenager enrolled in the French ANRS EPF-CO10 paediatric cohort: a case report. Lancet, 3:e49–54, 2016. doi.org/10.1016/S2352-3018(15)00232-5.

[13] Giacomet V, Trabattoni D, Zanchetta N, Biasin M, Gismondo M, Clerici M, and Zuccotti. No cure of HIV infection in a child despite early treatment and apparent viral clearance. Lancet, 384:1320, 2014. doi.org/10.1016/S0140-6736(14)61405-7.

[14] Luzuriaga K and Mofenson LM. Challenges in the elimination of pediatric HIV-1 infection. N Engl J Med, 374:761–770, 2016. doi.org/10.1056/NEJMra1505256.

[15] Sáez-Cirión A, Bacchus C, Hocqueloux L, Avettand-Fenoel V, Girault I, and et al. Post-treatment HIv-1 controllers with a long-term virological remission after the interruption of early initiated antiretroviral therapy ANRS VISCONTI study. PLoS Pathog., 9:e1003211, 2013. doi: 10.1371/journal.ppat.1003211.

[16] Prado JG and Frater J. Editorial: Immune surveillance of the HIV reservoir: Mechanisms, therapeutic targeting and new avenues for HIV cure. Front. Immunol, 11:doi: 10.3389/fimmu.2020.00070, 2020.

[17] Tomalka AG, Resto-Garay I, Campbell KS, and Popkin DL. In vitro evidence that combination therapy with CD16-bearing NK-92 cells and FDA-approved alefacept can selectively target the latent hiv reservoir in CD4+ CD2hi memory t cells. Front. Immunol, 9, 2018. doi: 10.3389/fimmu.2018.02552.

[18] Klein F, Mouquet H, Dosenovic P, Scheid JF, Scharf L, and Nussenzweig MC. Antibodies in HIV-1 vaccine development and therapy. Science, 341:p, 2013. doi.org/10.1126/science.1241144.

[19] Chun TW, Carruth L, Finzi D, Shen X, DiGiuseppe JA, Taylor H, Hermankova M, Chadwick K, Margolick J, Quinn TC, Kuo YH, Brookmeyer R, Zeiger MA, Barditch-Crovo P, and Siliciano RF. Quantification of latent tissue reservoirs and total body viral load in HIV-1 infection. Nature, 387:183–8, 1997. doi: 10.1038/387183a0.

[20] Finzi D, Hermankova M, Pierson T, Carruth LM, Buck C, Chaisson RE, Quinn TC, Chadwick K, Margolick J, Brookmeyer R, Gallant J, Markowitz M, Ho DD, Richman DD, and Siliciano RF. Identification of a reservoir for HIV-1 in patients on highly active antiretroviral therapy. Science, 278:1295–1300, 1997. doi.org/10.1126/science.278.5341.1295.

[21] Finzi D, Blankson J, Siliciano JD, Margolick JB, Chadwick K, Pierson T, Smith K, Lisziewicz J, Lori F, Flexner C, Quinn TC, Chaisson RE, Rosenberg E, Walker B, Gange S, Gallant J, and Siliciano RF. Latent infection of CD4 T cells provides a mechanism for lifelong persistence of HIV-1, even in patients on effective combination therapy. Nat Med, 5:512–517, 1999. doi.org/10.1038/8394.

[22] Whitney JB, Hill AL, Sanisetty S, Penaloza-MacMaster P, Liu J, Shetty M, Parenteau L, Cabral C, Shields J, Blackmore S, Smith JY, Brinkman AL, Peter LE, Mathew SI, Smith KM, Borducchi EN, Rosenbloom DI, Lewis MG, Hattersley J, Li B, Hesselgesser J, Geleziunas R, Robb ML, Kim JH, Michael NL, and Barouch DH. Rapid seeding of the viral reservoir prior to SIV viremia in rhesus monkeys. Nature, 512:74–7, 2014. doi.org/10.1038/nature13594.

[23] Ho YC, Shan L, Hosmane NN, Wang J, Laskey SB, Rosenbloom DI, Lai J, Blankson JN, Siliciano JD, and Siliciano RF. Replication-competent noninduced proviruses in the latent reservoir increase barrier to HIV-1 cure. Cell, 155:540–51, 2013. doi.org/10.1016/j.cell.2013.09.020.

[24] Siliciano JD, Kajdas J, Finzi D, Quinn TC, Chadwick K, Margolick JB, Kovacs C, Gange SJ, and Siliciano RF. Long-term follow-up studies confirm the stability of the latent reservoir for HIV-1 in resting CD4+ T cells. Nat Med, 9:727–8, 2003. doi.org/10.1038/nm880.

[25] Joos B, Fischer M, Kuster H, Pillai SK, Wong JK, Böni J, Hirschel B, Weber R, Trkola A, and Günthard HF; Swiss HIV Cohort Study. HIV rebounds from latently infected cells, rather than from continuing low-level replication. Proc Natl Acad Sci USA, 5:16725–30, 2008. doi.org/10.1073/pnas.0804192105.

[26] Shiau S and Kuhn L. Antiretroviral treatment in HIV-infected infants and young children: Novel issues raised by the mississippi baby. Expert Rev Anti Infect Ther, 12:307–18, 2014. doi: 10.1586/14787210.2014.888311.

[27] Ananworanich J, Schuetz A, Vandergeeten C, Sereti I, de Souza M, Rerknimitr R, Dewar R, Marovich M, van Griensven F, Sekaly R, Pinyakorn S, Phanuphak N, Trichavaroj R, Rutvisuttinunt W, Chomchey N, Paris R, Peel S, Valcour V, Maldarelli F, Chomont N, Michael N, Phanuphak P, and Kim JH; RV254/SEARCH 010 Study Group. Impact of multi-targeted antiretroviral treatment on gut T cell depletion and HIV reservoir seeding during acute hiv infection. PLoS ONE, 7:e33948, 2012. doi: 10.1371/journal.pone.0033948.

[28] Hocqueloux L, Avettand-Fènoël V, Jacquot S, Prazuck T, Legac E, Mélard A, Niang M, Mille C, Le Moal G, Viard JP, and Rouzioux C; AC32 (Coordinated Action on HIV Reservoirs) of the Agence Nationale de Recherches sur le Sida et les Hépatites Virales (ANRS). Long-term antiretroviral therapy initiated during primary HIV-1 infection is key to achieving both low HIV reservoirs and normal T cell counts. J Antimicrob Chemother, 68:1169–78, 2013. doi: 10.1093/jac/dks533.

[29] Siliciano JD and Siliciano RF. In vivo dynamics of the latent reservoir for HIV-1: New insights and implications for cure. Annu Rev Pathol, 24:271–94, 2022. doi: 10.1146/annurev-pathol-050520-112001.

[30] Obregon-Perko V, Bricker KM, Mensah G, Uddin F, Kumar MR, Fray EJ, Siliciano RF, Schoof N, Horner A, Mavigner M, Liang S, Vanderford T, Sass J, Chan C, Berendam SJ, Bar KJ, Shaw GM, Silvestri G, Fouda GG, Permar SR, and Chahroudi A. Simian-human immunodeficiency virus SHIV.C.CH505 persistence in ART-suppressed infant macaques is characterized by elevated SHIV RNA in the gut and a high abundance of intact SHIV DNA in naive CD4+ T cells. J Virol, 95:e01669–20, 2020. doi: 10.1128/JVI.01669-20.

[31] Hill AL, Rosenbloom DI, Fu F, Nowak MA, and Siliciano RF. Predicting the outcomes of treatment to eradicate the latent reservoir for HIV-1. Proc Natl Acad Sci USA, 111:13475–80, 2014. doi.org/10.1073/pnas.1406663111.

[32] Hill AL, Rosenbloom DI, Goldstein E, Hanhauser E, Kuritzkes DR, Siliciano RF, and Henrich TJ. Real-time predictions of reservoir size and rebound time during antiretroviral therapy interruption trials for HIV. PLoS Pathog, 12:e1005535, 2016. doi.org/10.1371/journal.ppat.1005535.

[33] Pinkevych M, Cromer D, Tolstrup M, Grimm AJ, Cooper DA, Lewin SR, Søgaard OS, Rasmussen TA, Kent SJ, Kelleher AD, and Davenport MP. HIV reactivation from latency after treatment interruption occurs on average every 5-8 days – implications for HIV remission. PLoS Pathog., 11:e1005000, 2015. doi.org/10.1371/journal.ppat.1005000.

[34] Pinkevych M, Kent SJ, Tolstrup M, Lewin SR, Cooper DA, Søgaard OS, Rasmussen TA, Kelleher AD, Cromer D, and Davenport MP. Modeling of experimental data supports HIV reactivation from latency after treatment interruption on average once every 5-8 days. PLoS Pathog, 12:e1005740, 2016. doi.org/10.1371/journal.ppat.1005740.

[35] Fennessey CM, Pinkevych M, Immonen TT, Reynaldi A, Venturi V, Nadella P, Reid C, Newman L, Lipkey L, Oswald K, Bosche WJ, Trivett MT, Ohlen C, Ott DE, Estes JD, Del Prete GQ, Lifson JD, Davenport MP, and Keele BF. Genetically-barcoded SIV facilitates enumeration of rebound variants and estimation of reactivation rates in nonhuman primates following interruption of suppressive antiretroviral therapy. PLOS Pathogens, 13:e1006359, 2017. doi.org/10.1371/journal.ppat.1006359.

[36] Conway JM, Perelson AS, and Li JZ. Predictions of time to HIV viral rebound following ART suspension that incorporate personal biomarkers. PLoS Comput. Biol., 15:e1007229, 2019. doi:10.1371/journal.pcbi.1007229.

[37] Conway JM, Meily P, Li JZ, and Perelson AS. Unified model of short- and long-term HIV viral rebound for clinical trial planning. J. R. Soc. Interface, 18:20201015, 2021. doi.org/10.1098/rsif.2020.1015.

[38] Mavigner M, Habib J, Deleage C, Rosen E, Mattingly C, Bricker K, Kashuba A, Amblard F, Schinazi RF, Lawson B, Vanderford TH, Jean S, Cohen J, McGary C, Paiardini M, Wood MP, Sodora DL, Silvestri G, Estes J, and Chahroudi A. Simian immunodeficiency virus persistence in cellular and anatomic reservoirs in antiretroviral therapy-suppressed infant Rhesus macaques. J Virol, 92:e00562–18, 2018. doi.org/10.1128/JVI.00562-18.

[39] Obregon-Perko V, Bricker KM, Mensah G, Uddin F, Rotolo L, Vanover D, Desai Y, Santangelo PJ, Jean Sand Wood JS, Connor-Stroud FC, Ehnert S, Berendam SJ, Liang S, Vanderford TH, Bar KJ, Shaw GM, Silvestri G, Kumar A, Fouda GG, Permar SR, and Chahroudi A. Dynamics and origin of rebound viremia in SHIV-infected infant macaques following interruption of long-term ART. JCI Insight, 6:e152526, 2021. doi: 10.1172/jci.insight.152526.

[40] Sarzotti-Kelsoe M, Bailer RT, Turk E, Lin CL, Bilska M, Greene KM, Gao H, Todd CA, Ozaki DA, Seaman MS, Mascola JR, and Montefiori DC. Optimization and validation of the TZM-bl assay for standardized assessments of neutralizing antibodies against HIV-1. J Immunol Methods, 409:131–46, 2014. doi: 10.1016/j.jim.2013.11.022.

[41] Bruner KM, Wang Z, Simonetti FR, Bender AM, Kwon KJ, Sengupta S, Fray EJ, Beg SA, Antar AAR, Jenike KM, Bertagnolli LN, Capoferri AA, Kufera JT, Timmons A, Nobles C, Gregg J, Wada N, Ho YC, Zhang H, Margolick JB, Blankson JN, Deeks SG, Bushman FD, Siliciano JD, Laird GM, and Siliciano RF. A quantitative approach for measuring the reservoir of latent HIV-1 proviruses. Nature, 566:120–5, 2019. doi: 10.1038/s41586-019-0898-8.

[42] Luebeck G and Meza R. Bhat: general likelihood exploration. version 0.9-10. 2015, 2015. CRAN.Rproject.org/package=Bhat.

[43] Akaike H. A new look at the statistical model identification. IEEE Transactions on Automatic Control, 19:716–723, 1974. doi:10.1109/TAC.1974.1100705.

[44] van Dorp CH, Conway JM, Barouch DH, Whitney JB, and Perelson AS. Models of SIV rebound after treatment interruption that involve multiple reactivation events. PLoS Comput Biol, 16:e1008241, 2020. doi.org/10.1371/journal.

[45] Burnham KP and Anderson DR. Model selection and multimodel inference. New York: Springer, 2004.

[46] Conway JM and Perelson AS. Post-treatment control of HIV infection. Proc Natl Acad Sci USA, 112:5467–72, 2015. doi: 10.1073/pnas.1419162112.

[47] Sharaf R, Lee GQ, Sun X, Etemad B, Aboukhater LM, Hu Z, Brumme ZL, Aga E, Bosch RJ, Wen Y, Namazi G, Gao C, Acosta EP, and et al. HIV-1 proviral landscapes distinguish posttreatment controllers from noncontrollers. J Clin Invest, 128:4074–4085, 2018. doi: 10.1172/JCI120549.

[48] Chomont N, El-Far M, Ancuta P, Trautmann L, Procopio FA, Yassine-Diab B, and et al. HIV reservoir size and persistence are driven by T cell survival and homeostatic proliferation. Nat Med, 15:893–901, 2009. doi.org/10.1038/nm.1972.

[49] Kearney MF, Wiegand A, Shao W, Coffin JM, Mellors JW, Lederman M, Ganghi RT, Keele BF, and Li JZ. Origin of rebound plasma HIV includes cells with identical proviruses that are transcriptionally active before stopping of antiretroviral therapy. JVirol, 90:1369–76, 2016. doi.org/10.1128/JVI.02139-15.

[50] Simonetti FR, Sobolewski MD, Fyne E, Shao W, Spindler J, Hattori J, Anderson EM, Watters SA, Hill S, Wu X, Wells D, Su L, Luke BT, Halvas EK, Besson G, Penrose KJ, Yang Z, Kwan RW, Van Waes C, Uldrick T, Citrin DE, Kovacs J, Polis MA, Rehm CA, Gorelick R, Piatak M, Keele BF, Kearney MF, Coffin JM, Hughes SH, Mellors JW, and Maldarelli F. Clonally expanded CD4+ T cells can produce infectious HIV-1 in vivo. Proc Natl Acad Sci USA, 113:1883–8, 2016. doi: 10.1073/pnas.1522675113.

[51] Maldarelli F, Wu X, Su L, Simonetti FR, Shao W, Hill S, Splindler L, Mellors JW, Kearney MF, Coffin JM, and Hughes SH. Specific HIV integration sites are linked to clonal expansion and persistence of infected cells. Science, 345:179–83, 2014. doi.or g/10.1126/science.1254194.

[52] Bertagnolli LN, Varriale J, Sweet S, Brockhurst J, Simonetti FR, White J, and et al. Autologous IgG antibodies block outgrowth of a substantial but variable fraction of viruses in the latent reservoir for HIV-1. Proc Natl Acad Sci USA, 117:32066–32077, 2022.

[53] Crooks AM, Bateson R, Cope AB, Dahl NP, Griggs MK, Kuruc JD, Gay CL, Eron JJ, Margolis DM, Bosch RJ, and Archin NM. Precise quantitation of the latent HIV-1 reservoir: Implications for eradication strategies. J Infect Dis, 212:1361–1365, 2015. doi.org/10.1093/infdis/jiv218.

[54] Zerbato JM, McMahon DK, Sobolewski MD, Mellors JW, and Sluis-Cremer N. Naive CD4+ T cells harbor a large inducible reservoir of latent, replication-competent Human Immunodeficiency Virus Type 1. Clin Infect Dis, 69:1919–25, 2019. doi: 10.1093/cid/ciz108.

[55] Persaud D, Pierson T, Ruff C, Finzi D, Chadwick KR, Margolick JB, Ruff A, Hutton N, Ray S, and Siliciano RF. A stable latent reservoir for HIV-1 in resting CD4(+) T lymphocytes in infected children. J Clin Invest, 105:995–1003, 2000. doi: 10.1172/JCI9006.

[56] Oda T, Kim KS, Fujita Y, Ito Y, and Iwami S Miura T. Quantifying antiviral effects against simian/human immunodeficiency virus induced by host immune response. J Theor Biol, 509, 2021. doi: 10.1016/j.jtbi.2020.110493.

[57] McBrien JB, Kumar NA, and Silvestri G. Mechanisms of CD8+ T cell-mediated suppression of HIV/SIV replication. Eur J Immunol, 48:898–914, 2018. doi: 10.1002/eji.201747172.

[58] Bosinger SE, Sodora DL, and Silvestri G. Generalized immune activation and innate immune responses in simian immunodeficiency virus infection. Curr Opin HIV AIDS, 6, 2011. doi: 10.1097/COH.0b013e3283499cf6.

[59] Schmitz JE, Kuroda MJ, Santra S, Sasseville VG, Simon MA, Lifton MA, Racz P, Tenner-Racz K, Dalesandro M, Scallon BJ, Ghrayeb J, Forman MA, Montefiori DC, Rieber EP, Letvin NL, and Reimann KA. Control of viremia in simian immunodeficiency virus infection by CD8+ lymphocytes. Science, 283:857–60, 1999. doi: 10.1126/science.283.5403.857.

[60] Ciupe SM, Miller CJ, and Forde JE. A bistable switch in virus dynamics can explain the differences in disease outcome following SIV infections in Rhesus macaques. Front. Microbiol, 9:1216, 2018. doi: 10.3389/fmicb.2018.01216.

[61] Sharma A, Boyd DF, and Overbaugh J. Development of SHIVs with circulating, transmitted hiv-1 variants. J Med Primatol, 44:296 –300, 2015. doi.org/10.1111/jmp.12179.

[62] Boyd DF, Peterson D, Haggarty BS, Jordan APO, Hogan MJ, Goo L, Hoxie JA, and Overbaugh J. Mutations in HIV-1 envelope that enhance entry with the macaque CD4 receptor alter antibody recognition by disrupting quaternary interactions within the trimer. J Virol, 89:894 –907, 2015. doi.org/10.1128/JVI.02680-14.

[63] Sagar M. HIV-1 transmission biology: selection and characteristics of infecting viruses. J Infect Dis, 202:S289 –S296, 2010. 10.1086/655656.

[64] Hemelaar J, Gouws E, Ghys PD, and Osmanov S. Global trends in molecular epidemiology of HIV-1 during 2000-2007. AIDS, 25:679 –689, 2011. doi.org/10.1097/QAD.0b013e328342ff93.

[65] Bar KJ, Coronado E, Hensley-McBain T, O’Connor MA, Osborn JM, Miller C, Gott TM, Wangari S, Iwayama N, Ahrens CY, Smedley J, Moats C, Lynch RM, Haddad EK, Haigwood NL, Fuller DH, Shaw GM, Klatt NR, and Manuzak JA. Simian-human immunodeficiency virus SHIV.CH505 infection of rhesus macaques results in persistent viral replication and induces intestinal immunopathology. J Virol, 93:e00372–19, 2019. doi.org/10.1128/JVI.00372-19.

[66] Fouda GG, Yates NL, Pollara J, Shen X, Overman GR, Mahlokozera T, Wilks AB, Kang HH, Salazar-Gonzalez JF, Salazar MG, Kalilani L, Meshnick SR, Hahn BH, Shaw GM, Lovingood RV, Denny TN, Haynes B, Letvin NL, Ferrari G, Montefiori DC, Tomaras GD, Permar SR, and the Center for HIV/AIDS Vaccine Immunology. HIV-specific functional antibody responses in breast milk mirror those in plasma and are primarily mediated by IgG antibodies. J Virol, 85, 2011. doi.org/10.1128/JVI.05174-11.

[67] Bélec L, Bouquety JC, Georges AJ, Siopathis MR, and Martin PM. Antibodies to HIV-1 in the breast milk of healthy, seropositive women. 1990, 85:1022–26, Pediatrics. doi.org/10.1542/peds.85.6.1022.

[68] Becquart P, Hocini H, Garin B, Kazatchkine MD Sépou A, and Bélec L. Compartmentalization of the igg immune response to hiv-1 in breast milk. AIDS, 13:1323–31, 1999. doi: 10.1097/00002030-199907300-00008.

[69] Henrick BM, Yao X-D, Nasser L, Roozrogousheh A, and Rosenthal KL. Breastfeeding behaviors and the innate immune system of human milk: Working together to protect infants against inflammation, HIV-1, and other infections. Front. Immunol, 8:doi: 10.3389/fimmu.2017.01631, 2017.

