## Supplementary Tables for "Assessing the impact of autologous neutralizing antibodies on viral rebound in postnatally SHIV-infected ART-treated infant rhesus macaques"

| Subject | Day | a | b | c | d | Undiluted Ne |
| --- | --- | --- | --- | --- | --- | --- |
| RIm19 | 0 | 0 | 7.73520802 | 1.96312352 | 67.1872215 | 66.8250383 |
| RIm19 | 14 | 0 | 10 | 2.13986687 | 20.0679982 | 20.0580345 |
| RIm19 | 31 | 0 | 10 | 1.16497104 | 100 | 82.1561424 |
| RJm19 | 0 | 0 | 4.20781398 | 2.05992017 | 90.0241339 | 85.9177565 |
| RJm19 | 14 | 0 | 10 | 1.49076797 | 43.9928791 | 43.1960891 |
| RJm19 | 31 | 0 | 3.4551969 | 1.52414391 | 69.6936084 | 56.517377 |
| RKg19 | 4 | 0 | 7.07600165 | 1.97872617 | 94.2568185 | 93.5092878 |
| RKg19 | 14 | 0 | 7.43118055 | 2.02315109 | 89.1819021 | 88.7100335 |
| RKg19 | 32 | 0 | 5.73846066 | 1.7656907 | 87.5030402 | 84.2760741 |
| RLg19 | 7 | 1.78081566 | 3.75372183 | 1.19819331 | 100 | 66.9448129 |
| RLg19 | 14 | 0 | 10 | 1.90776637 | 26.2106838 | 26.1697061 |
| RLg19 | 32 | 9.6732023 | 7.55766203 | 1.90838978 | 32.208487 | 32.039286 |
| RQc19 | 4 | 0 | 10 | 2.76001395 | 99.1117571 | 99.1078934 |
| RQc19 | 14 | 5.6233796 | 10 | 2.83514863 | 99.0794685 | 99.0766835 |
| RQc19 | 32 | 0 | 10 | 2.84106858 | 99.8231809 | 99.8202675 |
| RRm19 | 0 | 0 | 8.35574124 | 2.36745441 | 98.9882068 | 98.9144403 |
| RRm19 | 14 | 0 | 9.33712348 | 2.12865503 | 93.160498 | 93.0800903 |
| RRm19 | 31 | 0 | 9.69435644 | 2.37334482 | 96.9797422 | 96.9574732 |
| RTp19 | 0 | 0 | 8.26692109 | 1.99540759 | 70.3550378 | 70.1230191 |
| RTp19 | 10 | 0 | 6.43395782 | 2.12549225 | 33.2575121 | 32.9994962 |
| RTp19 | 31 | 0 | 10 | 1.91548307 | 32.1883806 | 32.1400454 |
| RVh19 | 4 | 0 | 9.78860346 | 2.35311303 | 88.8597567 | 88.8393045 |
| RVh19 | 14 | 0 | 10 | 2.28328053 | 87.2874055 | 87.2647462 |
| RVh19 | 32 | 0 | 8.55591977 | 3.11558743 | 97.7030118 | 97.6971613 |
| RVr19 | 0 | 0 | 10 | 2.30448772 | 95.4883798 | 95.46578 |
| RVr19 | 14 | 0 | 9.70135508 | 1.92428054 | 84.2393955 | 84.0925134 |
| RVr19 | 31 | 0 | 8.0947986 | 1.90086002 | 86.8706466 | 86.3937359 |
| Rwc19 | 0 | 3.09614279 | 5.42257387 | 2.05392441 | 92.6227583 | 90.8515822 |
| Rwc19 | 14 | 0 | 10 | 1.18080521 | 100 | 84.0501724 |
| Rwc19 | 31 | 0 | 7.45305257 | 1.16212092 | 99.9999518 | 75.3950631 |

| Model | Model Description | Parameter | Parameter Values | 95% CI |
| --- | --- | --- | --- | --- |
| 1 | Null | $\alpha(1-q)$<br>$\tau$ | 0.05<br>4.9 | (0.02, 0.1)<br>(2.05, 10.52) |
| 2 | $q=\min(q_0\phi, 1)$ | $\alpha$<br>$q_0$<br>$\tau$ | 0.51<br>0.95<br>5.7 | (0.14, 1.5)<br>(0.85, 1.06)<br>(3.49, 8.98) |
| 3 | $\tau=\tau_1+(\tau_2-\tau_1)\phi$ | $\alpha(1-q)$<br>$\tau_1$<br>$\tau_2$ | 0.05<br>3.37E-06<br>6.54 | (0.02, 0.1)<br>(3.36701e-6, 3.36704e-6)<br>(3.14, 12.38) |
| 4 | $\tau=\tau_1+(\tau_2-\tau_1)\phi^z$ | $\alpha(1-q)$<br>$\tau_1$<br>$\tau_2$ | 0.05<br>4.37E-06<br>6.07 | (0.004, 0.57)<br>(4.36820e-6, 4.36829e-6)<br>(0, 39.9) |
| 5 | $q=\min(q_0\phi, 1)$<br>$\tau=\tau_1+(\tau_2-\tau_1)\phi$ | $\alpha$<br>$q_0$<br>$\tau_1$<br>$\tau_2$ | 0.71<br>0.98<br>19.24<br>2.09 | (0.26, 1.67)<br>(0.93, 1.03)<br>(18.8, 19.4)<br>(1.77, 2.47) |
| 6 | $q=\min(q_0\phi, 1)$<br>$\tau=\tau_1+(\tau_2-\tau_1)\phi^z$ | $\alpha$<br>$q_0$<br>$\tau_1$<br>$\tau_2$ | 0.72<br>0.98<br>13.65<br>1.07 | (0.26, 1.66)<br>(0.93, 1.03)<br>(13.29, 14)<br>(0.25, 4.29) |
| 7 | LR size | $\alpha(1-q)$<br>$\tau$ | 0.01<br>4.9 | (0.007, 0.02)<br>(1.96, 11) |
| 8 | LR size<br>$q=\min(q_0\phi, 1)$ | $\alpha$<br>$q_0$<br>$\tau$ | 0.16<br>0.97<br>5.68 | (0.05, 0.43)<br>(0.89, 1.05)<br>(3.43, 9.03) |
| 9 | LR size<br>$\tau=\tau_1+(\tau_2-\tau_1)\phi$ | $\alpha(1-q)$<br>$\tau_1$<br>$\tau_2$ | 0.014<br>4.50E-06<br>6.5 | (0.007, 0.029)<br>(4.5013e-6, 4.501424e-6)<br>(3.07, 12.48) |
| 10 | LR size<br>$\tau=\tau_1+(\tau_2-\tau_1)\phi^z$ | $\alpha(1-q)$<br>$\tau_1$<br>$\tau_2$ | 0.015<br>4.50E-06<br>8.62 | (0.007, 0.031)<br>(4.5013e-6, 4.501415e-6)<br>(4.79, 14.29) |
| 11 | LR size<br>$q=\min(q_0\phi, 1)$<br>$\tau=\tau_1+(\tau_2-\tau_1)\phi$ | $\alpha$<br>$q_0$<br>$\tau_1$<br>$\tau_2$ | 0.2<br>0.98<br>19.12<br>2.19 | (0.08, 0.51)<br>(0.94, 1.02)<br>(18.5, 19.72)<br>(1.85, 2.58) |
| 12 | LR size<br>$q=\min(q_0\phi, 1)$<br>$\tau=\tau_1+(\tau_2-\tau_1)\phi^z$ | $\alpha$<br>$q_0$<br>$\tau_1$<br>$\tau_2$ | 0.2<br>0.98<br>13.62<br>1.15 | (0.08, 0.5)<br>(0.94, 1.02)<br>(13.29, 14.01)<br>(0.75, 1.76) |
| 13 | $\tau=\tau_1+\tau_2\phi$ | $\alpha(1-q)$<br>$\tau_1$<br>$\tau_2$ | 0.05<br>2.25E-06<br>6.54 | (0.02, 0.1)<br>(2.2506e-6, 2.25071e-6)<br>(3.14, 12.38) |
| 14 | $q=\min(q_0\phi, 1)$ | $\alpha$ | 0.51 | (0.14, 1.5) |
| | $\tau=\tau_1+\tau_2\phi$ | $q_0$ | 0.95 | (0.85, 1.06) |
| | | $\tau_1$ | 5.71 | (3.41, 8.74) |
| | | $\tau_2$ | 4.50E-06 | (4.5013e-6, 4.50147e-6) |
| 15 | LR size<br>$\tau=\tau_1+\tau_2\phi$ | $\alpha(1-q)$ | 0.014 | (0.007, 0.02) |
| | | $\tau_1$ | 4.50E-06 | (4.5013e-6, 4.50141e-6) |
| | | $\tau_2$ | 6.5 | (3.07, 12.48) |
| 16 | LR size<br>$q=\min(q_0\phi, 1)$<br>$\tau=\tau_1+\tau_2\phi$ | $\alpha$ | 0.16 | (0.05, 0.48) |
| | | $q_0$ | 0.97 | (0.89, 1.05) |
| | | $\tau_1$ | 5.68 | (3.43, 9.03) |
| | | $\tau_2$ | 4.50E-06 | (4.5013e-6, 4.50141e-6) |
| 17 | $\tau=\tau_1+(\tau_2+\tau_1)\phi$ | $\alpha(1-q)$ | 0.05 | (0.02, 0.1) |
| | | $\tau_1$ | 2.25E-06 | (2.2506e-6, 2.250709e-6) |
| | | $\tau_2$ | 6.54 | (3.14, 12.38) |
| 18 | $q=\min(q_0\phi, 1)$<br>$\tau=\tau_1+(\tau_2+\tau_1)\phi$ | $\alpha$ | 0.046 | (0.12, 1.43) |
| | | $q_0$ | 0.95 | (0.82, 1.07) |
| | | $\tau_1$ | 3.12 | (1.83, 5.06) |
| | | $\tau_2$ | 4.50E-06 | (4.5013e-6, 4.50142e-6) |
| 19 | LR size<br>$\tau=\tau_1+(\tau_2+\tau_1)\phi$ | $\alpha(1-q)$ | 0.015 | (0.007, 0.3) |
| | | $\tau_1$ | 4.50E-06 | (4.5013e-6, 4.501415e-6) |
| | | $\tau_2$ | 6.5 | (3.07, 12.49) |
| 20 | LR size<br>$q=\min(q_0\phi, 1)$<br>$\tau=\tau_1+(\tau_2+\tau_1)\phi$ | $\alpha$ | 0.15 | (0.048, 0.46) |
| | | $q_0$ | 0.96 | (0.84, 1.05) |
| | | $\tau_1$ | 3.12 | (1.85, 5.15) |
| | | $\tau_2$ | 4.50E-06 | (4.5012e-6, 4.5142e-06) |
| 21 | $q=\phi^n/(k^n+\phi^n)$, $n=1$ | $\alpha(1-q)$ | 4.99 | (NaN, NaN) |
| | | $k$ | 0.0098 | (NaN, NaN) |
| | | $\tau$ | 5.15 | (2.49, 9.9) |
| 22 | $q=\phi^n/(k^n+\phi^n)$, $n=2$ | $\alpha(1-q)$ | 4.99 | (0, 5) |
| | | $k$ | 0.0094 | (0.065, 0.13) |
| | | $\tau$ | 5.34 | (2.74, 9.75) |
| 23 | $q=\phi^n/(k^n+\phi^n)$, $n=3$ | $\alpha(1-q)$ | 1.4 | (0, 5) |
| | | $k$ | 0.3 | (0.00094, 83.23) |
| | | $\tau$ | 5.52 | (3.05, 9.46) |
| 24 | $q=q_0+q_1\phi$ | $\alpha$ | 0.67 | (0.2, 1.85) |
| | | $q_0$ | 0.24 | (NaN, NaN) |
| | | $q_1$ | 0.72 | (NaN, NaN) |
| | | $\tau$ | 5.71 | (3.49, 8.97) |
| 25 | LR size<br>$q=\phi^n/(k^n+\phi^n)$, $n=1$ | $\alpha$ | 6.12 | (0, 10) |
| | | $k$ | 0.002 | (4.16e-9, 1.99) |
| | | $\tau$ | 5.06 | (2.21, 10.53) |
| 26 | LR size<br>$q=\phi^n/(k^n+\phi^n)$, $n=2$ | $\alpha$ | 7.27 | (0, 10) |
| | | $k$ | 0.04 | (0.00065, 1.13) |
| | | $\tau$ | 5.24 | (2.55, 10.01) |
| 27 | LR size<br>$q=\phi^n/(k^n+\phi^n)$, $n=3$ | $\alpha$ | 5.02 | (0, 10) |
| | | $k$ | 0.12 | (7.06e-5, 1.98) |
| | | $\tau$ | 5.44 | (2.95, 9.49) |
| 28 | LR size<br>$q=q_0+q_1\phi$ | $\alpha$ | 0.31 | (00.1, 0.91) |
| | | $q_0$ | 0.49 | (NaN, NaN) |
| | | $q_1$ | 0.49 | (NaN, NaN) |
| | | $\tau$ | 5.68 | (3.43, 9.03) |
| 29 | $\tau=\tau_1+\tau_2\phi^n/(k^n+\phi^n)$, $n=1$ | $\alpha(1-q)$ | 0.05 | (0.026, 0.1) |
| | | $\tau_1$ | 1.96 | (NaN, NaN) |
| | | $\tau_2$ | 6.99 | (NaN, NaN) |
| | | $k$ | 1.03 | (NaN, NaN) |
| 30 | $\tau=\tau_1+\tau_2\phi^n/(k^n+\phi^n)$, $n=2$ | $\alpha(1-q)$ | 0.05 | (0.026, 0.1) |
| | | $\tau_1$ | 4.55 | (0.59, 14.7) |
| | | $\tau_2$ | 3.41 | (NaN, NaN) |
| | | $k$ | 1.66 | (0.04, 40.5) |
| 31 | $\tau=\tau_1+\tau_2\phi^n/(k^n+\phi^n)$, $n=3$ | $\alpha(1-q)$ | 0.053 | (0.027, 0.1) |
| | | $\tau_1$ | 4.66 | (1.3, 11.39) |
| | | $\tau_2$ | 3.25 | (NaN, NaN) |
| | | $k$ | 1.29 | (0.085, 16.58) |
| 32 | $q=\min(q_0\phi, 1)$<br>$\tau=\tau_1+\tau_2\phi^n/(k^n+\phi^n)$, $n=1$ | $\alpha$ | 0.51 | (0.14, 1.5) |
| | | $q_0$ | 0.95 | (0.85, 1.06) |
| | | $\tau_1$ | 5.44 | (NaN, NaN) |
| | | $\tau_2$ | 0.26 | (NaN, NaN) |
| | | $k$ | 1.20E-05 | (0, NaN) |
| 33 | $q=\min(q_0\phi, 1)$<br>$\tau=\tau_1+\tau_2\phi^n/(k^n+\phi^n)$, $n=2$ | $\alpha$ | 0.51 | (0.14, 1.5) |
| | | $q_0$ | 0.95 | (0.85, 1.06) |
| | | $\tau_1$ | 5.7 | (3.36, 8.81) |
| | | $\tau_2$ | 39.9 | (NaN, NaN) |
| | | $k$ | 57.02 | (0, 100) |
| 33 | $q=\min(q_0\phi, 1)$<br>$\tau=\tau_1+\tau_2\phi^n/(k^n+\phi^n)$, $n=3$ | $\alpha$ | 0.51 | (0.14, 1.5) |
| | | $q_0$ | 0.95 | (0.85, 1.06) |
| | | $\tau_1$ | 5.7 | (3.36, 8.81) |
| | | $\tau_2$ | 39.9 | (NaN, NaN) |
| | | $k$ | 96.7 | (0, NaN) |
| 34 | LR size<br>$\tau=\tau_1+\tau_2\phi^n/(k^n+\phi^n)$, $n=1$ | $\alpha(1-q)$ | 0.014 | (0.007, 0.02) |
| | | $\tau_1$ | 4.06 | (0.15, 30.7) |
| | | $\tau_2$ | 1.89 | (NaN, NaN) |
| | | $k$ | 0.98 | (0, 40) |
| 35 | LR size<br>$\tau=\tau_1+\tau_2\phi^n/(k^n+\phi^n)$, $n=2$ | $\alpha(1-q)$ | 0.014 | (0.007, 0.02) |
| | | $\tau_1$ | 4.91 | (1.96, 11) |
| | | $\tau_2$ | 1.24 | (NaN, NaN) |
| | | $k$ | 39.9 | (NaN, NaN) |
| 36 | LR size<br>$\tau=\tau_1+\tau_2\phi^n/(k^n+\phi^n)$, $n=3$ | $\alpha(1-q)$ | 0.015 | (0.008, 0.27) |
| | | $\tau_1$ | 4.42 | (NaN, NaN) |
| | | $\tau_2$ | 39.9 | (NaN, NaN) |
|  |  | k | 2.56 | (NaN, NaN) |
| 37 | LR size<br>$q=\min(q_0\phi, 1)$<br>$\tau=\tau_1+\tau_2\phi^n/(k^n+\phi^n)$, $n=1$ | $\alpha$ | 0.16 | (0.05, 0.48) |
| | | $q_0$ | 0.97 | (0.89, 1.05) |
| | | $\tau_1$ | 2.5 | (NaN, NaN) |
| | | $\tau_2$ | 3.18 | (NaN, NaN) |
|  |  | k | 1.12E-05 | (0, 100) |
| 38 | LR size<br>$q=\min(q_0\phi, 1)$<br>$\tau=\tau_1+\tau_2\phi^n/(k^n+\phi^n)$, $n=2$ | $\alpha$ | 0.16 | (0.05, 0.48) |
| | | $q_0$ | 0.97 | (0.89, 1.05) |
| | | $\tau_1$ | 5.6 | (3.46, 8.95) |
| | | $\tau_2$ | 23 | (NaN, NaN) |
|  |  | k | 92.85 | (0, 100) |
| 39 | LR size<br>$q=\min(q_0\phi, 1)$<br>$\tau=\tau_1+\tau_2\phi^n/(k^n+\phi^n)$, $n=3$ | $\alpha$ | 0.16 | (0.05, 0.48) |
| | | $q_0$ | 0.97 | (0.89, 1.05) |
| | | $\tau_1$ | 2.45 | (NaN, NaN) |
| | | $\tau_2$ | 3.23 | (NaN, NaN) |
|  |  | k | 0.003 | (0, 100) |
| 40 | LR size<br>$\tau=\tau_1-\tau_2LR$ | $\alpha(1-q)$ | 0.014 | (0.007, 0.02) |
| | | $\tau_1$ | 23.77 | (23.37, 24.17) |
| | | $\tau_2$ | 4.99 | (4.84, 5.14) |
| 41 | LR size<br>$\tau=\tau_1+\tau_2/LR$ | $\alpha(1-q)$ | 0.014 | (0.007, 0.029) |
| | | $\tau_1$ | 4.50E-06 | (4.50E-06, 4.50E-06) |
| | | $\tau_2$ | 19.95 | (7.3, 32.6) |
| 42 | LR size<br>$\tau=\tau_1/LR$ | $\alpha(1-q)$ | 0.014 | (0.007, 0.029) |
| | | $\tau_1$ | 19.95 | (8.39, 36.24) |
| 43 | LR size<br>$q=\min(q_0\phi, 1)$<br>$\tau=\tau_1+\tau_2LR$ | $\alpha$ | 0.22 | (0.09, 0.56) |
| | | $q_0$ | 0.98 | (0.94, 1.02) |
| | | $\tau_1$ | 22.42 | (22.1, 22.7) |
| | | $\tau_2$ | 4.5 | (4.43, 4.57) |
| 44 | LR size<br>$q=\min(q_0\phi, 1)$<br>$\tau=\tau_1/LR$ | $\alpha$ | 0.21 | (0.075, 0.58) |
| | | $q_0$ | 0.98 | (0.92, 1.03) |
| | | $\tau_1$ | 23.63 | (15.98, 30.31) |
| 45 | LR size<br>$\tau=\tau_1+\tau_2(1-LR^n/(k^n+LR^n))$, $n=1$ | $\alpha(1-q)$ | 0.014 | (0.007, 0.028) |
| | | $\tau_1$ | 4.5 | (0.67, 21.98) |
| | | $\tau_2$ | 11 | (NaN, NaN) |
|  |  | k | 0.23 | (NaN, NaN) |
| 46 | LR size<br>$\tau=\tau_1+\tau_2(1-LR^n/(k^n+LR^n)), n=2$ | $\alpha(1-q)$ | 0.014 | (0.007, 0.028) |
| | | $\tau_1$ | 4.49 | (0.17, 41.6) |
| | | $\tau_2$ | 59.9 | (NaN, NaN) |
|  |  | k | 0.44 | (0, 98.1) |
| 47 | LR size<br>$\tau=\tau_1+\tau_2(1-LR^n/(k^n+LR^n)), n=3$ | $\alpha(1-q)$ | 0.014 | (0.007, 0.028) |
| | | $\tau_1$ | 1.1 | (0.23, 4.84) |
| | | $\tau_2$ | 59.9 | (1, 60) |
|  |  | k | 1.53 | (1.42, 1.66) |
| 48 | LR size<br>$q=\min(q_0\phi, 1)$<br>$\tau=\tau_1+\tau_2(1-LR^n/(k^n+LR^n)), n=1$ | $\alpha$ | 0.14 | (0.025, 0.74) |
| | | $q_0$ | 0.95 | (0.78, 1.12) |
| | | $\tau_1$ | 2.77 | (0.63, 11.6) |
| | | $\tau_2$ | 19.78 | (0, 200) |
|  |  | k | 0.65 | (0.100) |
| 49 | LR size<br>$q=\min(q_0\phi, 1)$<br>$\tau=\tau_1+\tau_2(1-LR^n/(k^n+LR^n)), n=2$ | $\alpha$ | 0.24 | (0.079, 0.71) |
| | | $q_0$ | 0.98 | (0.93, 1.03) |
| | | $\tau_1$ | 1.85 | (1.59, 2.15) |
| | | $\tau_2$ | 199.9 | (0, 200) |
|  |  | k | 0.56 | (0.55, 0.57) |
| 50 | LR size<br>$q=\min(q_0\phi, 1)$<br>$\tau=\tau_1+\tau_2(1-LR^n/(k^n+LR^n)), n=3$ | $\alpha$ | 0.21 | (0.086, 0.54) |
| | | $q_0$ | 0.98 | (0.93, 1.03) |
| | | $\tau_1$ | 1.79 | (1.61, 1.98) |
| | | $\tau_2$ | 199.4 | (NaN, NaN) |
|  |  | k | 0.96 | (0.95, 0.98) |
| 51 | $q=\phi^n/(k^n+\phi^n), n=1$<br>$\tau=\tau_1+\tau_2\phi^n/(k^n+\phi^n), n=3$ | $\alpha$ | 4.99 | (NaN, NaN) |
| | | $\tau_1$ | 3.91 | (NaN, NaN) |
| | | $\tau_2$ | 1.25 | (NaN, NaN) |
|  |  | k | 0.009 | (0.004, 0.019) |
| 52 | $q=\phi^n/(k^n+\phi^n), n=1$<br>$\tau=\tau_1+\tau_2\phi^n/(k^n+\phi^n), n=2$ | $\alpha$ | 4.99 | (NaN, NaN) |
| | | $\tau_1$ | 3.98 | (NaN, NaN) |
| | | $\tau_2$ | 1.16 | (NaN, NaN) |
|  |  | k | 0.009 | (NaN, NaN) |
| 53 | $q=\phi^n/(k^n+\phi^n), n=1$<br>$\tau=\tau_1+\tau_2\phi^n/(k^n+\phi^n), n=3$ | $\alpha$ | 4.99 | (4.99, 4.996) |
| | | $\tau_1$ | 4.23 | (NaN, NaN) |
| | | $\tau_2$ | 0.91 | (0.01, 40.3) |
|  |  | k | 0.009 | (0.004, 0.019) |
| 54 | $q=\phi^n/(k^n+\phi^n), n=2$<br>$\tau=\tau_1+\tau_2\phi^n/(k^n+\phi^n), n=1$ | $\alpha$ | 4.99 | (NaN, NaN) |
| | | $\tau_1$ | 5.34 | (2.78, 10.11) |
| | | $\tau_2$ | 2.25E-05 | (0, NaN) |
|  |  | k | 0.093 | (0.066, 0.13) |
| 55 | $q=\phi^n/(k^n+\phi^n)$, n=2 | $\alpha$ | 4.99 | (NaN, NaN) |
| | $\tau=\tau_1+\tau_2\phi^n/(k^n+\phi^n)$, n=2 | $\tau_1$ | 5.34 | (2.78, 10.12) |
| | | $\tau_2$ | 2.25E-05 | 0, NaN) |
|  |  | k | 0.093 | (0.066, 0.13) |
| 56 | $q=\phi^n/(k^n+\phi^n)$, n=2 | $\alpha$ | 4.99 | (4.99, 4.99) |
| | $\tau=\tau_1+\tau_2\phi^n/(k^n+\phi^n)$, n=3 | $\tau_1$ | 0.6 | (NaN, NaN) |
| | | $\tau_2$ | 4.74 | (2.29, 9.67) |
|  |  | k | 0.093 | (0.066, 0.13) |
| 57 | $q=\phi^n/(k^n+\phi^n)$, n=3 | $\alpha$ | 1.42 | (0, 5) |
| | $\tau=\tau_1+\tau_2\phi^n/(k^n+\phi^n)$, n=1 | $\tau_1$ | 5.52 | (3.11, 9.69) |
| | | $\tau_2$ | 2.25E-05 | (2.25E-05, 2.25E-05) |
|  |  | k | 0.3 | (0.0008, 72.63) |
| 58 | $q=\phi^n/(k^n+\phi^n)$, n=3 | $\alpha$ | 1.45 | (0, 5) |
| | $\tau=\tau_1+\tau_2\phi^n/(k^n+\phi^n)$, n=2 | $\tau_1$ | 5.52 | (3.11, 9.69) |
| | | $\tau_2$ | 2.25E-05 | (2.25E-05, 2.25E-05) |
|  |  | k | 0.3 | (0.0007, 78.43) |
| 59 | $q=\phi^n/(k^n+\phi^n)$, n=3 | $\alpha$ | 1.4 | (0, 5) |
| | $\tau=\tau_1+\tau_2\phi^n/(k^n+\phi^n)$, n=3 | $\tau_1$ | 5.51 | (3.11, 9.69) |
| | | $\tau_2$ | 2.25E-05 | (2.25E-05, 2.25E-05) |
|  |  | k | 0.3 | (0.0009, 66.9) |
| 60 | LR size | $\alpha$ | 6.21 | (0, 10) |
| | $q=\phi^n/(k^n+\phi^n)$, n=1 | $\tau_1$ | 3.46 | (NaN, NaN) |
| | $\tau=\tau_1+\tau_2\phi^n/(k^n+\phi^n)$, n=1 | $\tau_2$ | 1.6 | (NaN, NaN) |
|  |  | k | 0.0021 | (0, 87.6) |
| 61 | LR size | $\alpha$ | 6.51 | (0, 10) |
| | $q=\phi^n/(k^n+\phi^n)$, n=1 | $\tau_1$ | 2.06 | (NaN, NaN) |
| | $\tau=\tau_1+\tau_2\phi^n/(k^n+\phi^n)$, n=2 | $\tau_2$ | 2.99 | (NaN, NaN) |
|  |  | k | 0.0021 | (0, 61.02) |
| 62 | LR size | $\alpha$ | 5.21 | (0, 10) |
| | $q=\phi^n/(k^n+\phi^n)$, n=1 | $\tau_1$ | 3.24 | (NaN, NaN) |
| | $\tau=\tau_1+\tau_2\phi^n/(k^n+\phi^n)$, n=3 | $\tau_2$ | 1.81 | (NaN, NaN) |
|  |  | k | 0.002 | (0, 99.98) |
| 63 | LR size | $\alpha$ | 9.03 | (0, 10) |
| | $q=\phi^n/(k^n+\phi^n)$, n=2 | $\tau_1$ | 4.8 | (NaN, NaN) |
| | $\tau=\tau_1+\tau_2\phi^n/(k^n+\phi^n)$, n=1 | $\tau_2$ | 0.46 | (NaN, NaN) |
|  |  | k | 0.03 | (0.01, 0.1) |
| 64 | LR size | $\alpha$ | 8.58 | (0, 10) |
| | $q=\phi^n/(k^n+\phi^n)$, $n=1$ | $\tau_1$ | 3.88 | (NaN, NaN) |
| | $\tau=\tau_1+\tau_2\phi^n/(k^n+\phi^n)$, $n=2$ | $\tau_2$ | 1.35 | (NaN, NaN) |
| | | $k$ | 0.03 | (0.005, 0.27) |
| 65 | LR size | $\alpha$ | 7.97 | (0, 10) |
| | $q=\phi^n/(k^n+\phi^n)$, $n=2$ | $\tau_1$ | 3.73 | (NaN, NaN) |
| | $\tau=\tau_1+\tau_2\phi^n/(k^n+\phi^n)$, $n=3$ | $\tau_2$ | 1.5 | (NaN, NaN) |
| | | $k$ | 0.03 | (0.002, 0.68) |
| 66 | LR size | $\alpha$ | 5.5 | (0, 10) |
| | $q=\phi^n/(k^n+\phi^n)$, $n=3$ | $\tau_1$ | 5.44 | (3.01, 9.74) |
| | $\tau=\tau_1+\tau_2\phi^n/(k^n+\phi^n)$, $n=1$ | $\tau_2$ | 0.00041 | (0.000407, 0.0004073) |
| | | $k$ | 0.12 | (0.0003, 29.37) |
| 67 | LR size | $\alpha$ | 9.81 | (0, 10) |
| | $q=\phi^n/(k^n+\phi^n)$, $n=3$ | $\tau_1$ | 5.44 | (3.01, 9.74) |
| | $\tau=\tau_1+\tau_2\phi^n/(k^n+\phi^n)$, $n=2$ | $\tau_2$ | 2.25E-05 | (2.25E-05, 2.25E-05) |
| | | $k$ | 0.1 | (0.07, 0.14) |
| 68 | LR size | $\alpha$ | 0.05 | (NaN, NaN) |
| | $q=\phi^n/(k^n+\phi^n)$, $n=3$ | $\tau_1$ | 5 | (3.01, 8.24) |
| | $\tau=\tau_1+\tau_2\phi^n/(k^n+\phi^n)$, $n=3$ | $\tau_2$ | 2.00E+00 | (NaN, NaN) |
| | | $k$ | 0.67 | (NaN, NaN) |
| 69 | LR size | $\alpha$ | 5.19 | (0, 10) |
| | $q=\phi^n/(k^n+\phi^n)$, $n=1$ | $\tau_1$ | 23.7 | (23.47, 23.9) |
| | $\tau=\tau_1-\tau_2\text{LR}$ | $\tau_2$ | 4.97 | (4.83, 5.11) |
| | | $k$ | 0.002 | (0, 99.45) |
| 70 | LR size | $\alpha$ | 5.77 | (0, 10) |
| | $q=\phi^n/(k^n+\phi^n)$, $n=2$ | $\tau_1$ | 23.63 | (23.37, 23.86) |
| | $\tau=\tau_1-\tau_2\text{LR}$ | $\tau_2$ | 4.93 | (4.79, 5.07) |
| | | $k$ | 0.04 | (0, 31.81) |
| 71 | LR size | $\alpha$ | 9.04 | (0, 10) |
| | $q=\phi^n/(k^n+\phi^n)$, $n=3$ | $\tau_1$ | 23.9 | (23.73, 24.19) |
| | $\tau=\tau_1-\tau_2\text{LR}$ | $\tau_2$ | 5.06 | (4.92, 5.19) |
| | | $k$ | 0.1 | (0.043, 0.27) |
| 72 | LR size | $\alpha$ | 9.99 | (NaN, NaN) |
| | $q=\phi^n/(k^n+\phi^n)$, $n=1$ | $\tau_1$ | 1.12E-05 | (0, NaN) |
| | $\tau=\tau_1+\tau_2/\text{LR}$ | $\tau_2$ | 20.68 | (10.28, 37.23) |
| | | $k$ | 0.0012 | (0.0006, 0.002) |
| 73 | LR size | $\alpha$ | 9.99 | (NaN, NaN) |
| | $q=\phi^n/(k^n+\phi^n)$, $n=2$ | $\tau_1$ | 1.12E-05 | (1.12E-05, 1.12E-05) |
| | $\tau=\tau_1+\tau_2\phi/\text{LR}$ | $\tau_2$ | 21.55 | (13.55, 32.5) |
| | | $k$ | 0.03 | (0.02, 0.04) |
| 74 | LR size | $\alpha$ | 0.01 | (0, 9.99) |
| | $q=\phi^n/(k^n+\phi^n)$, $n=3$ | $\tau_1$ | 5 | (2.93, 8.38) |
| | $\tau=\tau_1+\tau_2/LR$ | $\tau_2$ | 3 | (0.26, 26.2) |
| | | $k$ | 0.1 | (0.001, 5.69) |
| 75 | LR size | $\alpha$ | 6.32 | (0, 10) |
| | $q=\phi^n/(k^n+\phi^n)$, $n=1$ | $\tau_1$ | 20.68 | (9.82, 35.13) |
| | $\tau=\tau_1/LR$ | $k$ | 0.002 | (0, 76.26) |
| 76 | LR size | $\alpha$ | 9.43 | (0, 10) |
| | $q=\phi^n/(k^n+\phi^n)$, $n=2$ | $\tau_1$ | 21.5 | (11.69, 33.89) |
| | $\tau=\tau_1/LR$ | $k$ | 0.03 | (0.01, 0.08) |
| 77 | LR size | $\alpha$ | 0.1 | (0.0002, 8.10) |
| | $q=\phi^n/(k^n+\phi^n)$, $n=3$ | $\tau_1$ | 11.65 | (8.77, 15.2) |
| | $\tau=\tau_1/LR$ | $k$ | 0.1 | (0.008, 1.22) |
| 78 | LR size | $\alpha$ | 9.99 | (0, NaN) |
| | $q=\phi^n/(k^n+\phi^n)$, $n=1$ | $\tau_1$ | 3.37E-05 | (0, NaN) |
| | $\tau=\tau_1+\tau_2(1-LR^n/(k^n+LR^n))$, $n=1$ | $\tau_2$ | 299.9 | (0, NaN) |
| | | $k_q$ | 0.001 | (0.0006, 0.002) |
| | | $k_d$ | 0.07 | (0.035, 0.13) |
| 79 | LR size | $\alpha$ | 9.39 | (0, 10) |
| | $q=\phi^n/(k^n+\phi^n)$, $n=1$ | $\tau_1$ | 0.04 | (0.004, 0.44) |
| | $\tau=\tau_1+\tau_2(1-LR^n/(k^n+LR^n))$, $n=2$ | $\tau_2$ | 299.99 | (0, 300) |
| | | $k_q$ | 0.001 | (0.0002, 0.009) |
| | | $k_d$ | 0.51 | (0.5, 0.517) |
| 80 | LR size | $\alpha$ | 7.16 | (0, 10) |
| | $q=\phi^n/(k^n+\phi^n)$, $n=1$ | $\tau_1$ | 1.6 | (1.35, 1.89) |
| | $\tau=\tau_1+\tau_2(1-LR^n/(k^n+LR^n))$, $n=3$ | $\tau_2$ | 299.9 | (0, 300) |
| | | $k_q$ | 0.001 | (3.07, 10.73) |
| | | $k_d$ | 0.84 | (0.83, 0.85) |
| 81 | LR size | $\alpha$ | 0.01 | (0, 7.3) |
| | $q=\phi^n/(k^n+\phi^n)$, $n=2$ | $\tau_1$ | 3 | (0.53, 16.1) |
| | $\tau=\tau_1+\tau_2(1-LR^n/(k^n+LR^n))$, $n=1$ | $\tau_2$ | 3 | (NaN, NaN) |
| | | $k_q$ | 0.1 | (0.007, 1.31) |
| | | $k_d$ | 0.1 | (NaN, NaN) |
| 82 | LR size | $\alpha$ | 9.12 | (0, 10) |
| | $q=\phi^n/(k^n+\phi^n)$, $n=2$ | $\tau_1$ | 0.02 | (NaN, NaN) |
| | $\tau=\tau_1+\tau_2(1-LR^n/(k^n+LR^n))$, $n=2$ | $\tau_2$ | 299.9 | (0, 300) |
| | | $k_q$ | 0.03 | (0.01, 0.14) |
| | | $k_d$ | 0.51 | (0.5, 0.517) |
| 83 | LR size<br>$q=\phi^n/(k^n+\phi^n)$, $n=2$<br>$\tau=\tau_1+\tau_2(1-LR^n/(k^n+LR^n))$, $n=3$ | $\alpha$<br>$\tau_1$<br>$\tau_2$<br>$k_q$<br>$k_d$ | 0.01<br>3<br>3<br>0.1<br>0.1 | (0, 7.28)<br>(2.5, 3.5)<br>(NaN, NaN)<br>(0.007, 1.31)<br>(NaN, NaN) |
| 84 | LR size<br>$q=\phi^n/(k^n+\phi^n)$, $n=3$<br>$\tau=\tau_1+\tau_2(1-LR^n/(k^n+LR^n))$, $n=1$ | $\alpha$<br>$\tau_1$<br>$\tau_2$<br>$k_q$<br>$k_d$ | 9.99<br>3.37E-05<br>79.81<br>0.1<br>0.3 | (NaN, NaN)<br>(0, 300)<br>(0.008, 299.9)<br>(0.08, 0.13)<br>(0.0001, 84.05) |
| 85 | LR size<br>$q=\phi^n/(k^n+\phi^n)$, $n=3$<br>$\tau=\tau_1+\tau_2(1-LR^n/(k^n+LR^n))$, $n=2$ | $\alpha$<br>$\tau_1$<br>$\tau_2$<br>$k_q$<br>$k_d$ | 2.79<br>0.29<br>140.3<br>0.16<br>0.75 | (0, 10)<br>(0.026, 3.14)<br>(135.1, 145.5)<br>(0.003, 6.62)<br>(0.73, 0.76) |
| 86 | LR size<br>$q=\phi^n/(k^n+\phi^n)$, $n=3$<br>$\tau=\tau_1+\tau_2(1-LR^n/(k^n+LR^n))$, $n=3$ | $\alpha$<br>$\tau_1$<br>$\tau_2$<br>$k_q$<br>$k_d$ | 5.79<br>0.37<br>26.1<br>0.12<br>2.3 | (0, 10)<br>(0.24, 0.57)<br>(25.2, 27.1)<br>(0.003, 4.15)<br>(2.27, 2.33) |
| 87 | LR size<br>$\tau=\tau_1+\tau_2\phi/LR$ | $\alpha(1-q)$<br>$\tau_1$<br>$\tau_2$ | 0.01<br>1.12E-05<br>26.08 | (0.007, 0.03)<br>(1.12E-05, 1.12E-05)<br>(13.18, 45.05) |
| 88 | LR size<br>$\tau=\tau_1+\tau_2\phi-\tau_3LR$ | $\alpha(1-q)$<br>$\tau_1$<br>$\tau_2$<br>$\tau_3$ | 0.017<br>9.45<br>38.85<br>9.67 | (0.09, 0.03)<br>(9.25, 9.65)<br>(38.58, 39.13)<br>(9.6, 9.74) |
| 89 | LR size<br>$q=\min(q_0\phi, 1)$<br>$\tau=\tau_1+\tau_2\phi/LR$ | $\alpha$<br>$q_0$<br>$\tau_1$<br>$\tau_2$ | 0.16<br>0.96<br>1.12E-05<br>28.44 | (0.05, 0.5)<br>(0.88, 1.05)<br>(1.12E-05, 1.12E-05)<br>(18.41, 41.17) |
| 90 | LR size<br>$q=\min(q_0\phi, 1)$<br>$\tau=\tau_1+\tau_2\phi-\tau_3LR$ | $\alpha$<br>$q_0$<br>$\tau_1$<br>$\tau_2$<br>$\tau_3$ | 0.29<br>0.98<br>23.5<br>22.4<br>9.28 | (0.11, 0.71)<br>(0.94, 1.02)<br>(23.3, 23.7)<br>(22.1, 22.6)<br>(9.22, 9.34) |
| 91 | LR size<br>$q=\phi^n/(k^n+\phi^n)$, $n=1$<br>$\tau=\tau_1+\tau_2\phi/LR$ | $\alpha$ | 5.23 | (0, 10) |
| | | $k$ | 0.002 | (0, 2) |
| | | $\tau_1$ | 3.29 | (NaN, NaN) |
| | | $\tau_2$ | 8.75 | (NaN, NaN) |
| 92 | LR size<br>$q=\phi^n/(k^n+\phi^n)$, $n=2$<br>$\tau=\tau_1+\tau_2\phi/LR$ | $\alpha$ | 8.04 | (0, 10) |
| | | $k$ | 0.04 | (0.002, 0.52) |
| | | $\tau_1$ | 1.12E-05 | (1.12E-05, 1.12E-05) |
| | | $\tau_2$ | 27.02 | (15.2, 43.35) |
| 93 | LR size<br>$q=\phi^n/(k^n+\phi^n)$, $n=3$<br>$\tau=\tau_1+\tau_2\phi/LR$ | $\alpha$ | 4.79 | (0, 10) |
| | | $k$ | 0.13 | (0, 2) |
| | | $\tau_1$ | 1.12E-05 | (1.12E-05, 1.12E-05) |
| | | $\tau_2$ | 27.63 | (16.57, 42.32) |
| 94 | LR size<br>$q=\phi^n/(k^n+\phi^n)$, $n=1$<br>$\tau=\tau_1+\tau_2\phi-\tau_3LR$ | $\alpha$ | 8.77 | (0, 10) |
| | | $k$ | 0.002 | (0, 0.06) |
| | | $\tau_1$ | 27.06 | (26.7, 27.4) |
| | | $\tau_2$ | 29.54 | (29.1, 29.9) |
| | | $\tau_3$ | 12.19 | (12.08, 12.3) |
| 95 | LR size<br>$q=\phi^n/(k^n+\phi^n)$, $n=2$<br>$\tau=\tau_1+\tau_2\phi-\tau_3LR$ | $\alpha$ | 9.87 | (0, 10) |
| | | $k$ | 0.04 | (0, 0.06) |
| | | $\tau_1$ | 23.6 | (22.9, 24.4) |
| | | $\tau_2$ | 28.04 | (27.2, 28.8) |
| | | $\tau_3$ | 10.82 | (10.5, 11.06) |
| 96 | LR size<br>$q=\phi^n/(k^n+\phi^n)$, $n=3$<br>$\tau=\tau_1+\tau_2\phi-\tau_3LR$ | $\alpha$ | 8.76 | (0, 10) |
| | | $k$ | 0.11 | (0.03, 0.33) |
| | | $\tau_1$ | 25.03 | (24.63, 25.4) |
| | | $\tau_2$ | 24.7 | (24.3, 25.1) |
| | | $\tau_3$ | 10.32 | (10.21, 10.44) |
| 97 | LR size<br>$\tau=\tau_1+\tau_2\phi-\tau_3LR+\tau_4\phi/LR$ | $\alpha(1-q)$ | 0.017 | (0.009, 0.03) |
| | | $\tau_1$ | 10.38 | (NaN, NaN) |
| | | $\tau_2$ | 4.36 | (NaN, NaN) |
| | | $\tau_3$ | 4.51 | (NaN, NaN) |
| | | $\tau_4$ | 43.49 | (NaN, NaN) |
| 98 | LR size<br>$\tau=\tau_1+\tau_2\phi-\tau_3LR+\tau_4\phi/LR$ | $\alpha(1-q)$ | 0.015 | (0.007, 0.03) |
| | | $\tau_1$ | 3.37E-05 | (3.37E-05, 3.37E-05) |
| | | $\tau_2$ | 3.37E-05 | (3.37E-05, 3.37E-05) |
| | | $\tau_3$ | 3.37E-05 | (3.37E-05, 3.37E-05) |
| | | $\tau_4$ | 26.08 | (13.78, 47.53) |
| 99 | LR size<br>$q=\min(q_0\phi, 1)$<br>$\tau=\tau_1+\tau_2\phi-\tau_3LR+\tau_4\phi/LR$ | $\alpha$ | 0.22 | (0.08, 0.61) |
| | | $q_0$ | 0.97 | (0.92, 1.03) |
| | | $\tau_1$ | 34.53 | (34.1, 34.9) |
| | | $\tau_2$ | 37.1 | (35.8, 38.3) |
| | | $\tau_3$ | 16.8 | (16.7, 16.9) |
| | | $\tau_4$ | 10.06 | (6.63, 15.17) |
| 100 | LR size<br>$q=\phi^n/(k^n+\phi^n)$, $n=1$<br>$\tau=\tau_1+\tau_2\phi-\tau_3LR+\tau_4\phi/LR$ | $\alpha$ | 6.05 | (0, 10) |
| | | $k$ | 0.002 | (0, 2) |
| | | $\tau_1$ | 2.99 | (2.68, 3.35) |
| | | $\tau_2$ | 6.95 | (6.57, 7.35) |
| | | $\tau_3$ | 6.32 | (6.24, 6.41) |
| | | $\tau_4$ | 92.34 | (91.29, 93.4) |
| 101 | LR size<br>$q=\phi^n/(k^n+\phi^n)$, $n=2$<br>$\tau=\tau_1+\tau_2\phi-\tau_3LR+\tau_4\phi/LR$ | $\alpha$ | 0.01 | (0, 5.46) |
| | | $k$ | 0.1 | (0.019, 0.44) |
| | | $\tau_1$ | 4 | (3.44, 4.65) |
| | | $\tau_2$ | 3.1 | (2.5, 3.84) |
| | | $\tau_3$ | 1 | (0.85, 1.16) |
| | | $\tau_4$ | 1.9 | (0.83, 4.33) |
| 102 | LR size<br>$q=\phi^n/(k^n+\phi^n)$, $n=3$<br>$\tau=\tau_1+\tau_2\phi-\tau_3LR+\tau_4\phi/LR$ | $\alpha$ | 0.01 | (0, 9.98) |
| | | $k$ | 0.1 | (0.02, 0.41) |
| | | $\tau_1$ | 4.87 | (NaN, NaN) |
| | | $\tau_2$ | 2.15 | (NaN, NaN) |
| | | $\tau_3$ | 1 | (0, 300) |
| | | $\tau_4$ | 1.32 | (NaN, NaN) |
| 103 | LR size<br>$q=\min(q_0\phi, 1)$<br>$\tau=\tau_1+\tau_2\phi-\tau_3LR+\tau_4\phi/LR$ | $\alpha$ | 0.21 | (0.07, 0.58) |
| | | $q_0$ | 0.98 | (0.92, 1.03) |
| | | $\tau_1$ | 3.37E-05 | (3.37E-05, 3.37E-05) |
| | | $\tau_2$ | 3.37E-05 | (3.37E-05, 3.37E-05) |
| | | $\tau_3$ | 23.6 | (17.16, 32.2) |
| | | $\tau_4$ | 3.37E-05 | (3.37E-05, 3.37E-05) |
| 104 | LR size<br>$q=\phi^n/(k^n+\phi^n)$, $n=1$<br>$\tau=\tau_1+\tau_2\phi+\tau_3/LR+\tau_4\phi/LR$ | $\alpha$ | 9.99E+00 | (NaN, NaN) |
| | | $k$ | 0.0014 | (0.0007, 0.002) |
| | | $\tau_1$ | 3.37E-05 | (0, NaN) |
| | | $\tau_2$ | 3.37E-05 | (0, NaN) |
| | | $\tau_3$ | 3.37E-05 | (0, NaN) |
| | | $\tau_4$ | 26.5 | (14.56, 46.6) |
| 105 | LR size | $\alpha$ | 0.01 | (0, 5.72) |
| | $q = \phi^n / (k^n + \phi^n), n=2$<br>$\tau = \tau_1 + \tau_2 \phi + \tau_3 / LR + \tau_4 \phi / LR$ | k<br>$\tau_1$<br>$\tau_2$<br>$\tau_3$<br>$\tau_4$ | 0.1<br>3<br>1<br>1<br>0.5 | (0.019, 0.44)<br>(0.69, 12.6)<br>(0.02, 30.34)<br>(0.001, 204.5)<br>(0, 293.9) |
| 106 | LR size<br>$q = \phi^n / (k^n + \phi^n), n=3$<br>$\tau = \tau_1 + \tau_2 \phi + \tau_3 / LR + \tau_4 \phi / LR$ | $\alpha$<br>k<br>$\tau_1$<br>$\tau_2$<br>$\tau_3$<br>$\tau_4$ | 0.01<br>0.1<br>3<br>1<br>1<br>0.5 | (0, 10)<br>(0.02, 0.42)<br>(0.9, 9.8)<br>(0.29, 13.05)<br>(0.05, 120.5)<br>(0.002, 168.6) |
| 107 | LR size<br>$\tau = \tau_1 + \tau_2 \phi - \tau_3 LR + \tau_4 \phi * LR$ | $\alpha(1-q)$<br>$\tau_1$<br>$\tau_2$<br>$\tau_3$<br>$\tau_4$ | 0.018<br>5.08<br>40.35<br>8.51<br>-0.32 | (0.009, 0.3)<br>(4.72, 5.47)<br>(39.9, 40.7)<br>(8.42, 8.61)<br>(-0.44, -0.21) |
| 108 | LR size<br>$q = \min(q_0 \phi, 1)$<br>$\tau = \tau_1 + \tau_2 \phi - \tau_3 LR + \tau_4 \phi * LR$ | $\alpha$<br>$q_0$<br>$\tau_1$<br>$\tau_2$<br>$\tau_3$<br>$\tau_4$ | 0.29<br>0.98<br>17.09<br>33.9<br>7.41<br>-3.32 | (0.11, 0.71)<br>(0.94, 1.02)<br>(16.9, 17.24)<br>(33.74, 34.2)<br>(7.35, 7.46)<br>(-3.4, -3.24) |
| 109 | LR size<br>$q = \phi^n / (k^n + \phi^n), n=1$<br>$\tau = \tau_1 + \tau_2 \phi - \tau_3 LR + \tau_4 \phi * LR$ | $\alpha$<br>k<br>$\tau_1$<br>$\tau_2$<br>$\tau_3$<br>$\tau_4$ | 7.66<br>0.002<br>17.69<br>24.7<br>10.9<br>2.84 | (0, 10)<br>(0, 0.61)<br>(17.32, 18.06)<br>(24.31, 25.111)<br>(10.78, 11.01)<br>(2.71, 2.96) |
| 110 | LR size<br>$q = \phi^n / (k^n + \phi^n), n=2$<br>$\tau = \tau_1 + \tau_2 \phi - \tau_3 LR + \tau_4 \phi * LR$ | $\alpha$<br>k<br>$\tau_1$<br>$\tau_2$<br>$\tau_3$<br>$\tau_4$ | 0.01<br>0.1<br>3.73<br>1.97<br>1<br>0.5 | (0, 5.48)<br>(0.019, 0.44)<br>(0, 300)<br>(0, 299.9)<br>(0, 300)<br>(0.34, 0.65) |
| 111 | LR size<br>$q = \phi^n / (k^n + \phi^n), n=3$<br>$\tau = \tau_1 + \tau_2 \phi - \tau_3 LR + \tau_4 \phi * LR$ | $\alpha$<br>k<br>$\tau_1$ | 0.01<br>0.1<br>3.73 | (0, 10)<br>(0.02, 0.42)<br>(0, 300) |

|  |  |  |  |
| --- | --- | --- | --- |
| | $\tau_2$ | 1.97 | (0, 300) |
| | $\tau_3$ | 1 | (0, 300) |
| | $\tau_4$ | 0.5 | (0.44, 0.55) |

| AIC | $\Delta AIC$ |
| --- | --- |
| 60.99 | 0 |
| 58.91 | -2.08 |
| 62.5 | 1.51 |
| 62.32 | 1.33 |
| 59.18 | -1.81 |
| 59.5 | -1.49 |
| 61.74 | 0.75 |
| 58.67 | -2.32 |
| 63.2 | 2.21 |
| 62.36 | 1.37 |
| 59.44 | -1.55 |
| 59.78 | -1.21 |
| 62.5 | 1.51 |
| 60.91 | -0.08 |
| 63.1 | 2.11 |
| 60.68 | -0.31 |
| 62.5 | 1.51 |
| 61.13 | 0.14 |
| 63.2 | 2.21 |
| 60.84 | -0.15 |
| 61.6 | 0.61 |
| 60.63 | -0.36 |
| 60.2 | -0.79 |
| 60.91 | -0.08 |
| 62.17 | 1.18 |
| 60.97 | -0.02 |
| 60.26 | -0.73 |
| 60.67 | -0.32 |
| 64.85 | 3.86 |
| 64.89 | 3.9 |
| 64.84 | 3.85 |
| 62.91 | 1.92 |
| 62.91 | 1.92 |
| 62.91 | 1.92 |
| 65.7 | 4.71 |
| 65.74 | 4.75 |
| 65.36 | 4.37 |
| 62.67 | 1.68 |
| 62.67 | 1.68 |
| 62.67 | 1.68 |
| 62.51 | 1.52 |
| 63.33 | 2.34 |
| 61.33 | 0.34 |
| 56.86 | -4.13 |
| 56.88 | -4.11 |
| 65.7 | 4.71 |
| 65.63 | 4.64 |
| 64.8 | 3.81 |
| 62.23 | 1.24 |
| 59.66 | -1.33 |
| 59.42 | -1.57 |
| 63.6 | 2.61 |
| 63.6 | 2.61 |
| 63.6 | 2.61 |
| 62.63 | 1.64 |
| 62.63 | 1.64 |
| 62.63 | 1.64 |
| 62.2 | 1.21 |
| 62.2 | 1.21 |
| 62.2 | 1.21 |
| 64.1 | 3.11 |
| 64.1 | 3.11 |
| 64.17 | 3.18 |
| 62.96 | 1.97 |
| 62.96 | 1.97 |
| 62.96 | 1.97 |
| 62.25 | 1.26 |
| 62.25 | 1.26 |
| 63.61 | 2.62 |
| 62.5 | 1.51 |
| 60.81 | -0.18 |
| 59.26 | -1.73 |
| 63.59 | 2.6 |
| 62.11 | 1.12 |
| 170.17 | 109.18 |
| 61.6 | 0.61 |
| 60.11 | -0.88 |
| 124.54 | 63.55 |
| 65.61 | 4.62 |
| 64.9 | 3.91 |
| 64.91 | 3.92 |
| 131.85 | 70.86 |
| 63.14 | 2.15 |
| 131.81 | 70.82 |
| 63.08 | 2.09 |
| 61.69 | 0.7 |
| 61.54 | 0.55 |
| 62.78 | 1.79 |
| 60.05 | -0.94 |
| 59.83 | -1.16 |
| 55.06 | -5.93 |
| 63.86 | 2.87 |
| 62.06 | 1.07 |
| 61.4 | 0.41 |
| 60.14 | -0.85 |
| 58.77 | -2.22 |
| 57.26 | -3.73 |
| 64.2 | 3.21 |
| 66.78 | 5.79 |
| 58.98 | -2.01 |
| 62.57 | 1.58 |
| 133.8 | 72.81 |
| 171.81 | 110.82 |
| 62.88 | 1.89 |
| 67.23 | 6.24 |
| 134.48 | 73.49 |
| 173.27 | 112.28 |
| 62.12 | 1.13 |
| 56.93 | -4.06 |
| 62.62 | 1.63 |
| 133.81 | 72.82 |
| 171.81 | 110.82 |

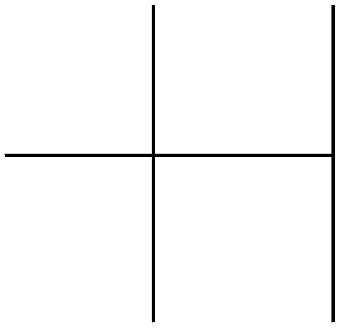

\_\_\_\_\_

\_\_\_\_\_

\_\_\_\_\_

\_\_\_\_\_

\_\_\_\_\_

\_\_\_\_\_

\_\_\_\_\_

\_\_\_\_\_

\_\_\_\_\_

\_\_\_\_\_

\_\_\_\_\_

\_\_\_\_\_

\_\_\_\_\_

\_\_\_\_\_

\_\_\_\_\_

\_\_\_\_\_

\_\_\_\_\_

\_\_\_\_\_

\_\_\_\_\_

\_\_\_\_\_

\_\_\_\_\_

\_\_\_\_\_

\_\_\_\_\_

\_\_\_\_\_

\_\_\_\_\_

\_\_\_\_\_

\_\_\_\_\_

\_\_\_\_\_

\_\_\_\_\_

\_\_\_\_\_

\_\_\_\_\_

\_\_\_\_\_

\_\_\_\_\_

\_\_\_\_\_

\_\_\_\_\_

\_\_\_\_\_

\_\_\_\_\_

\_\_\_\_\_

\_\_\_\_\_

\_\_\_\_\_

\_\_\_\_\_

\_\_\_\_\_

\_\_\_\_\_

\_\_\_\_\_

\_\_\_\_\_

\_\_\_\_\_

\_\_\_\_\_

\_\_\_\_\_

\_\_\_\_\_

\_\_\_\_\_

| Model | Model Description | Parameters | Parameter Value | 95% CI |
| --- | --- | --- | --- | --- |
| 1 | Null | $\alpha(1-q)$<br>$\tau$ | 0.11<br>6.07 | (0.059, 0.21)<br>(4.4, 8.2) |
| 2 | $q=\min(q_0\phi, 1)$ | $\alpha$<br>$q_0$<br>$\tau$ | 0.11<br>0.002<br>6.07 | (0.06, 0.23)<br>(1e-14, 2)<br>(4.41, 8.22) |
| 3 | $\tau=\tau_1+(\tau_2-\tau_1)\phi$ | $\alpha(1-q)$<br>$\tau_1$<br>$\tau_2$ | 0.11<br>6.07<br>5.38 | (0.06, 0.22)<br>(NaN, NaN)<br>(NaN, NaN) |
| 4 | $\tau=\tau_1+(\tau_2-\tau_1)\phi^2$ | $\alpha(1-q)$<br>$\tau_1$<br>$\tau_2$ | 0.11<br>5.22<br>18.08 | (0.06, 0.22)<br>(NaN, NaN)<br>(NaN, NaN) |
| 5 | $q=\min(q_0\phi, 1)$<br>$\tau=\tau_1+(\tau_2-\tau_1)\phi$ | $\alpha$<br>$q_0$<br>$\tau_1$<br>$\tau_2$ | 0.11<br>0.0005<br>4.97<br>9.89 | (0.06, 0.22)<br>(NaN, NaN)<br>(NaN, NaN)<br>(NaN, NaN) |
| 6 | $q=\min(q_0\phi, 1)$<br>$\tau=\tau_1+(\tau_2-\tau_1)\phi^2$ | $\alpha$<br>$q_0$<br>$\tau_1$<br>$\tau_2$ | 0.11<br>0.37<br>5.49<br>14.33 | (0.04, 0.4)<br>(1e-14, NaN)<br>(NaN, NaN)<br>(NaN, NaN) |
| 7 | LR size | $\alpha(1-q)$<br>$\tau$ | 0.04<br>6.08 | (0.02, 0.07)<br>(4.42, 8.22) |
| 8 | LR size<br>$q=\min(q_0\phi, 1)$ | $\alpha$<br>$q_0$<br>$\tau$ | 0.04<br>0.0009<br>608 | (0.0178, 0.08)<br>(1e-14, 2)<br>(4.42, 8.21) |
| 9 | LR size<br>$\tau=\tau_1+(\tau_2-\tau_1)\phi$ | $\alpha(1-q)$<br>$\tau_1$<br>$\tau_2$ | 0.03<br>5.43<br>4.45 | (0.01, 0.06)<br>(3.65, 7.85)<br>(NaN, NaN) |
| 10 | LR size<br>$\tau=\tau_1+(\tau_2-\tau_1)\phi^2$ | $\alpha(1-q)$<br>$\tau_1$<br>$\tau_2$ | 0.03<br>4.87<br>22.4 | (0.02, 0.07)<br>(NaN, NaN)<br>(NaN, NaN) |
| 11 | LR size<br>$q=\min(q_0\phi, 1)$<br>$\tau=\tau_1+(\tau_2-\tau_1)\phi$ | $\alpha$<br>$q_0$<br>$\tau_1$<br>$\tau_2$ | 0.03<br>2.25E-07<br>6.1<br>5.99 | (0.03, 0.055)<br>(NaN, NaN)<br>(0.027, 93.8)<br>(1.76E-8, 99.99) |
| 12 | LR size<br>$q=\min(q_0\phi, 1)$ | $\alpha$<br>$q_0$ | 0.03<br>0.01 | (0.01, 0.07)<br>(4.31E-9, 1.99) |
| | $\tau=\tau_1+(\tau_2-\tau_1)\phi^2$ | $\tau_1$ | 5.72 | (NaN, NaN) |
| | | $\tau_2$ | 11.18 | (NaN, NaN) |
| 13 | $\tau=\tau_1+\tau_2\phi$ | $\alpha(1-q)$ | 0.11 | (0.05, 0.21) |
| | | $\tau_1$ | 4.9 | (NaN, NaN) |
| | | $\tau_2$ | 5.22 | (NaN, NaN) |
| 14 | $q=\min(q_0\phi, 1)$ | $\alpha$ | 0.11 | (0.06, 0.22) |
| | $\tau=\tau_1+\tau_2\phi$ | $q_0$ | 0.0002 | (NaN, NaN) |
| | | $\tau_1$ | 3.77 | (NaN, NaN) |
| | | $\tau_2$ | 10.04 | (NaN, NaN) |
| 15 | LR size | $\alpha(1-q)$ | 0.037 | (0.02, 0.07) |
| | $\tau=\tau_1+\tau_2\phi$ | $\tau_1$ | 5.81E+00 | (NaN, NaN) |
| | | $\tau_2$ | 1.22 | (NaN, NaN) |
| 16 | LR size | $\alpha$ | 0.037 | (0.02, 0.08) |
| | $q=\min(q_0\phi, 1)$ | $q_0$ | 0.001 | (1e-14, 2) |
| | $\tau=\tau_1+\tau_2\phi$ | $\tau_1$ | 5.89 | (NaN, NaN) |
| | | $\tau_2$ | 8.20E-01 | (NaN, NaN) |
| 17 | $\tau=\tau_1+(\tau_2+\tau_1)\phi$ | $\alpha(1-q)$ | 0.11 | (0.05, 0.21) |
| | | $\tau_1$ | 3.04 | (NaN, NaN) |
| | | $\tau_2$ | 10.02 | (NaN, NaN) |
| 18 | $q=\min(q_0\phi, 1)$ | $\alpha$ | 0.11 | (0.034, 0.33) |
| | $\tau=\tau_1+(\tau_2+\tau_1)\phi$ | $q_0$ | 0.28 | (NaN, NaN) |
| | | $\tau_1$ | 4.13 | (NaN, NaN) |
| | | $\tau_2$ | 4.38E+00 | (NaN, NaN) |
| 19 | LR size | $\alpha(1-q)$ | 0.04 | (0.019, 0.07) |
| | $\tau=\tau_1+(\tau_2+\tau_1)\phi$ | $\tau_1$ | 3.39 | (NaN, NaN) |
| | | $\tau_2$ | 8.14 | (NaN, NaN) |
| 20 | LR size | $\alpha$ | 0.039 | (0.02, 0.06) |
| | $q=\min(q_0\phi, 1)$ | $q_0$ | 0.18 | (3.69e-7, 1.99) |
| | $\tau=\tau_1+(\tau_2+\tau_1)\phi$ | $\tau_1$ | 3.85 | (NaN, NaN) |
| | | $\tau_2$ | 5.96 | (NaN, NaN) |
| 21 | $q=\phi^n/(k^n+\phi^n), n=1$ | $\alpha(1-q)$ | 0.11 | (0.06, 0.21) |
| | | $k$ | 407.62 | (1.78e-9, 500) |
| | | $\tau$ | 6.07 | (4.41, 8.22) |
| 22 | $q=\phi^n/(k^n+\phi^n), n=2$ | $\alpha(1-q)$ | 0.11 | (0.05, 0.21) |
| | | $k$ | 498.88 | (NaN, NaN) |
| | | $\tau$ | 6.07 | (4.4, 8.24) |
| 23 | $q=\phi^n/(k^n+\phi^n), n=3$ | $\alpha(1-q)$ | 0.11 | (0.05, 0.22) |
|  |  | k | 2.92 | (0.36, 22.96) |
| | | $\tau$ | 6.07 | (4.41, 8.22) |
| 24 | $q=q_0+q_1\phi$ | $\alpha$ | 0.11 | (0.049, 0.26) |
| | | $q_0$ | 0.11 | (1.1e-9, 0.99) |
| | | $q_1$ | 0.0003 | (1e-14, NaN) |
| | | $\tau$ | 6.07 | (4.41, 8.22) |
| 25 | LR size<br>$q=\phi^n/(k^n+\phi^n)$, $n=1$ | $\alpha$ | 0.042 | (0.021, 0.08) |
|  |  | k | 1.99 | (5.89e-13, 2) |
| | | $\tau$ | 6.08 | (4.42, 8.21) |
| 26 | LR size<br>$q=\phi^n/(k^n+\phi^n)$, $n=2$ | $\alpha$ | 0.03 | (0.019, 0.07) |
|  |  | k | 1.99 | (1.99, 1.99) |
| | | $\tau$ | 6.08 | (4.42, 8.22) |
| 27 | LR size<br>$q=\phi^n/(k^n+\phi^n)$, $n=3$ | $\alpha$ | 0.03 | (0.019, 0.07) |
|  |  | k | 1.99 | (1.99, 1.99) |
| | | $\tau$ | 6.08 | (4.42, 8.22) |
| 28 | LR size<br>$q=q_0+q_1\phi$ | $\alpha$ | 0.06 | (0.03, 0.11) |
| | | $q_0$ | 0.41 | (0.34, 0.5) |
| | | $q_1$ | 0.0002 | (NaN, NaN) |
| | | $\tau$ | 6.08 | (4.42, 8.22) |
| 29 | $\tau=\tau_1+\tau_2\phi^n/(k^n+\phi^n)$, $n=1$ | $\alpha(1-q)$ | 0.11 | (0.05, 0.21) |
| | | $\tau_1$ | 4.22 | (NaN, NaN) |
| | | $\tau_2$ | 2.86 | (NaN, NaN) |
|  |  | k | 0.08 | (NaN, NaN) |
| 30 | $\tau=\tau_1+\tau_2\phi^n/(k^n+\phi^n)$, $n=2$ | $\alpha(1-q)$ | 0.11 | (0.05, 0.21) |
| | | $\tau_1$ | 4.22 | (NaN, NaN) |
| | | $\tau_2$ | 2.49 | (NaN, NaN) |
|  |  | k | 0.08 | (NaN, NaN) |
| 31 | $\tau=\tau_1+\tau_2\phi^n/(k^n+\phi^n)$, $n=3$ | $\alpha(1-q)$ | 0.11 | (0.05, 0.21) |
| | | $\tau_1$ | 3.16 | (NaN, NaN) |
| | | $\tau_2$ | 3.7 | (NaN, NaN) |
|  |  | k | 0.08 | (NaN, NaN) |
| 32 | $q=\min(q_0\phi, 1)$<br>$\tau=\tau_1+\tau_2\phi^n/(k^n+\phi^n)$, $n=1$ | $\alpha$ | 0.11 | (NaN, NaN) |
| | | $q_0$ | 0.0002 | (NaN, NaN) |
| | | $\tau_1$ | 3.72 | (NaN, NaN) |
| | | $\tau_2$ | 3.74 | (NaN, NaN) |
|  |  | k | 0.08 | (NaN, NaN) |
| 33 | $q=\min(q_0\phi, 1)$<br>$\tau=\tau_1+\tau_2\phi^n/(k^n+\phi^n)$, $n=2$ | $\alpha$ | 0.11 | (0.03, 0.38) |
| | | $q_0$ | 0.25 | (2.18e-12, 2) |
| | | $\tau_1$ | 3.86 | (NaN, NaN) |
| | | $\tau_2$ | 2.23 | (NaN, NaN) |
|  |  | k | 0.004 | (1.24e-6, 12.13) |
| 33 | $q=\min(q_0\phi, 1)$<br>$\tau=\tau_1+\tau_2\phi^n/(k^n+\phi^n), n=3$ | $\alpha$ | 0.11 | (0.03, 0.38) |
| | | $q_0$ | 0.25 | (2.18e-12, 2) |
| | | $\tau_1$ | 3.86 | (NaN, NaN) |
| | | $\tau_2$ | 2.23 | (NaN, NaN) |
|  |  | k | 0.004 | (1.24e-6, 12.13) |
| 34 | LR size<br>$\tau=\tau_1+\tau_2\phi^n/(k^n+\phi^n), n=1$ | $\alpha(1-q)$ | NaN | (NaN, NaN) |
| | | $\tau_1$ | NaN | (NaN, NaN) |
| | | $\tau_2$ | NaN | (NaN, NaN) |
|  |  | k | NaN | (NaN, NaN) |
| 35 | LR size<br>$\tau=\tau_1+\tau_2\phi^n/(k^n+\phi^n), n=2$ | $\alpha(1-q)$ | 0.045 | (0.04, 0.05) |
| | | $\tau_1$ | 1.73 | (1.43, 2.1) |
| | | $\tau_2$ | 299.99 | (NaN, NaN) |
|  |  | k | 0.4 | (0.39, 0.41) |
| 36 | LR size<br>$\tau=\tau_1+\tau_2\phi^n/(k^n+\phi^n), n=3$ | $\alpha(1-q)$ | NaN | (NaN, NaN) |
| | | $\tau_1$ | NaN | (NaN, NaN) |
| | | $\tau_2$ | NaN | (NaN, NaN) |
|  |  | k | NaN | (NaN, NaN) |
| 37 | LR size<br>$q=\min(q_0\phi, 1)$<br>$\tau=\tau_1+\tau_2\phi^n/(k^n+\phi^n), n=1$ | $\alpha$ | 0.02 | (3.53e-5, 1.99) |
| | | $q_0$ | 5.43 | (0.006, 283.52) |
| | | $\tau_1$ | 1.23 | (6.15e-11, 300) |
| | | $\tau_2$ | 0.16 | (1e-14, 100) |
|  |  | k | 4.00E-02 | (0.017, 0.08) |
| 38 | LR size<br>$q=\min(q_0\phi, 1)$<br>$\tau=\tau_1+\tau_2\phi^n/(k^n+\phi^n), n=2$ | $\alpha$ | 0.03 | (0.027, 0.051) |
| | | $q_0$ | 9.36E-05 | (NaN, NaN) |
| | | $\tau_1$ | 4.7 | (3.54, 6.55) |
| | | $\tau_2$ | 1.32 | (0.79, 2.81) |
|  |  | k | 0.0007 | (0.027, 0.05) |
| 39 | LR size<br>$q=\min(q_0\phi, 1)$<br>$\tau=\tau_1+\tau_2\phi^n/(k^n+\phi^n), n=3$ | $\alpha$ | 0.003 | (0.019, 0.07) |
| | | $q_0$ | 8.64E-05 | (NaN, NaN) |
| | | $\tau_1$ | 4.62 | (3.85, 5.55) |
| | | $\tau_2$ | 1.49 | (0.19, 11.27) |
|  |  | k | 0.01 | (0.0007, 0.17) |
| 40 | LR size<br>$\tau=\tau_1-\tau_2\text{LR}$ | $\alpha(1-q)$ | 0.046 | (0.02, 0.08) |
| | | $\tau_1$ | 30.4 | (29.97, 30.8) |
| | | $\tau_2$ | 7.27 | (7.1, 7.44) |
| 41 | LR size<br>$\tau=\tau_1+\tau_2/LR$ | $\alpha(1-q)$ | 0.04 | (0.02, 0.08) |
| | | $\tau_1$ | 3.37E-05 | (3.37e-5, 3.37e-5) |
| | | $\tau_2$ | 21.23 | (16.76, 26.77) |
| 42 | LR size<br>$\tau=\tau_1/LR$ | $\alpha(1-q)$ | 0.043 | (0.02, 0.08) |
| | | $\tau_1$ | 21.23 | (16.76, 26.77) |
| 43 | LR size<br>$q=\min(q_0\phi, 1)$<br>$\tau=\tau_1-\tau_2LR$ | $\alpha$ | 0.04 | (0.02, 0.08) |
| | | $q_0$ | 0.001 | (1e-14, 2) |
| | | $\tau_1$ | 30.74 | (30.04, 30.9) |
| | | $\tau_2$ | 7.3 | (7.11, 7.49) |
| 44 | LR size<br>$q=\min(q_0\phi, 1)$<br>$\tau=\tau_1/LR$ | $\alpha$ | 0.43 | (0.02, 0.09) |
| | | $q_0$ | 0.019 | (8.11e-9, 1.99) |
| | | $\tau_1$ | 21.22 | (16.75, 26.78) |
| 45 | LR size<br>$\tau=\tau_1+\tau_2(1-LR^n/(k^n+LR^n)), n=1$ | $\alpha(1-q)$ | 0.03 | (0.02, 0.06) |
| | | $\tau_1$ | 5.23 | (NaN, NaN) |
| | | $\tau_2$ | 1.14 | (NaN, NaN) |
|  |  | k | 0.05 | (NaN, NaN) |
| 46 | LR size<br>$\tau=\tau_1+\tau_2(1-LR^n/(k^n+LR^n)), n=2$ | $\alpha(1-q)$ | 0.03 | (0.01, 0.07) |
| | | $\tau_1$ | 4.92 | (NaN, NaN) |
| | | $\tau_2$ | 1.91 | (NaN, NaN) |
|  |  | k | 0.14 | (5.54e-10, 99.9) |
| 47 | LR size<br>$\tau=\tau_1+\tau_2(1-LR^n/(k^n+LR^n)), n=3$ | $\alpha(1-q)$ | 0.03 | (0.02, 0.07) |
| | | $\tau_1$ | 5.21 | (NaN, NaN) |
| | | $\tau_2$ | 0.9 | (NaN, NaN) |
|  |  | k | 0.01 | (NaN, NaN) |
| 48 | LR size<br>$q=\min(q_0\phi, 1)$<br>$\tau=\tau_1+\tau_2(1-LR^n/(k^n+LR^n)), n=1$ | $\alpha$ | NaN | (NaN, NaN) |
| | | $q_0$ | NaN | (NaN, NaN) |
| | | $\tau_1$ | NaN | (NaN, NaN) |
| | | $\tau_2$ | NaN | (NaN, NaN) |
|  |  | k | NaN | (NaN, NaN) |
| 49 | LR size<br>$q=\min(q_0\phi, 1)$<br>$\tau=\tau_1+\tau_2(1-LR^n/(k^n+LR^n)), n=2$ | $\alpha$ | 1.12E-05 | (NaN, NaN) |
| | | $q_0$ | 1.93 | (1.79, 2.08) |
| | | $\tau_1$ | 299.9 | (NaN, NaN) |
| | | $\tau_2$ | 0.39 | (0.39, 0.4) |
|  |  | k | 0.045 | (0.04, 0.048) |
| 50 | LR size<br>$q=\min(q_0\phi, 1)$<br>$\tau=\tau_1+\tau_2(1-LR^n/(k^n+LR^n)), n=3$ | $\alpha$ | NaN | (NaN, NaN) |
| | | $q_0$ | NaN | (NaN, NaN) |
| | | $\tau_1$ | NaN | (NaN, NaN) |
| | | $\tau_2$ | NaN | (NaN, NaN) |

|  |  | k | NaN | (NaN, NaN) |
| --- | --- | --- | --- | --- |
| 51 | $q=\phi^n/(k^n+\phi^n)$, n=1<br>$\tau=\tau_1+\tau_2\phi^n/(k^n+\phi^n)$, n=3 | $\alpha$<br>$\tau_1$<br>$\tau_2$<br>k | 0.11<br>5.83<br>44.09<br>39.35 | (0.05, 0.22)<br>(3.14, 10.68)<br>(3.21, 166.09)<br>(3.71, 157.67) |
| 52 | $q=\phi^n/(k^n+\phi^n)$, n=1<br>$\tau=\tau_1+\tau_2\phi^n/(k^n+\phi^n)$, n=2 | $\alpha$<br>$\tau_1$<br>$\tau_2$<br>k | 0.13<br>2.40E-05<br>199.9<br>1.54 | (0.06, 0.26)<br>(NaN, NaN)<br>(1e-14, NaN)<br>(1.35, 1.76) |
| 53 | $q=\phi^n/(k^n+\phi^n)$, n=1<br>$\tau=\tau_1+\tau_2\phi^n/(k^n+\phi^n)$, n=3 | $\alpha$<br>$\tau_1$<br>$\tau_2$<br>k | 0.11<br>6.07<br>199.96<br>197.9 | (0.05, 0.22)<br>(4.44, 8.28)<br>(NaN, NaN)<br>(197.7, 198.05) |
| 54 | $q=\phi^n/(k^n+\phi^n)$, n=2<br>$\tau=\tau_1+\tau_2\phi^n/(k^n+\phi^n)$, n=1 | $\alpha$<br>$\tau_1$<br>$\tau_2$<br>k | 0.11<br>5.76<br>1.02E+01<br>6.9 | (0.05, 0.22)<br>(2.64, 12.31)<br>(0.42, 115.34)<br>(0.24, 101.57) |
| 55 | $q=\phi^n/(k^n+\phi^n)$, n=2<br>$\tau=\tau_1+\tau_2\phi^n/(k^n+\phi^n)$, n=2 | $\alpha$<br>$\tau_1$<br>$\tau_2$<br>k | 0.11<br>6.07<br>4.77<br>7.57 | (0.05, 0.22)<br>(4.43, 8.28)<br>(NaN, NaN)<br>(NaN, NaN) |
| 56 | $q=\phi^n/(k^n+\phi^n)$, n=2<br>$\tau=\tau_1+\tau_2\phi^n/(k^n+\phi^n)$, n=3 | $\alpha$<br>$\tau_1$<br>$\tau_2$<br>k | 0.11<br>6.07<br>4.57<br>180.05 | (0.05, 0.22)<br>(4.44, 8.28)<br>(1e-14, NaN)<br>(0.003, 199.9) |
| 57 | $q=\phi^n/(k^n+\phi^n)$, n=3<br>$\tau=\tau_1+\tau_2\phi^n/(k^n+\phi^n)$, n=1 | $\alpha$<br>$\tau_1$<br>$\tau_2$<br>k | 0.11<br>5.46<br>199.8<br>72.56 | (0.05, 0.22)<br>(NaN, NaN)<br>(1e-14, NaN)<br>(NaN, NaN) |
| 58 | $q=\phi^n/(k^n+\phi^n)$, n=3<br>$\tau=\tau_1+\tau_2\phi^n/(k^n+\phi^n)$, n=2 | $\alpha$<br>$\tau_1$<br>$\tau_2$<br>k | 0.11<br>6.07<br>4.4<br>199.9 | (0.05, 0.22)<br>(4.44, 8.27)<br>(1e-14, NaN)<br>(NaN, NaN) |
| 59 | $q=\phi^n/(k^n+\phi^n)$, n=3<br>$\tau=\tau_1+\tau_2\phi^n/(k^n+\phi^n)$, n=3 | $\alpha$<br>$\tau_1$<br>$\tau_2$<br>k | 0.11<br>6.07<br>33.05<br>3.76 | (0.05, 0.22)<br>(4.44, 8.27)<br>(NaN, NaN)<br>(NaN, NaN) |
| 60 | LR size | $\alpha$ | 0.038 | (0.019, 0.075) |
| | $q=\phi^n/(k^n+\phi^n)$, $n=1$<br>$\tau=\tau_1+\tau_2\phi^n/(k^n+\phi^n)$, $n=1$ | $\tau_1$<br>$\tau_2$<br>$k_q$<br>$k_d$ | 5<br>2.01<br>38.13<br>0.15 | (NaN, NaN)<br>(NaN, NaN)<br>(1.27, 96.71)<br>(NaN, NaN) |
| 61 | LR size<br>$q=\phi^n/(k^n+\phi^n)$, $n=1$<br>$\tau=\tau_1+\tau_2\phi^n/(k^n+\phi^n)$, $n=2$ | $\alpha$<br>$\tau_1$<br>$\tau_2$<br>$k_q$<br>$k_d$ | 0.38<br>4.55<br>1.64<br>45.21<br>0.023 | (NaN, NaN)<br>(1.46, 13.85)<br>(0.09, 35.46)<br>(NaN, NaN)<br>(NaN, NaN) |
| 62 | LR size<br>$q=\phi^n/(k^n+\phi^n)$, $n=1$<br>$\tau=\tau_1+\tau_2\phi^n/(k^n+\phi^n)$, $n=3$ | $\alpha$<br>$\tau_1$<br>$\tau_2$<br>$k_q$<br>$k_d$ | 0.039<br>4.34<br>8.71<br>7.66<br>0.41 | (0.033, 0.045)<br>(3.59, 5.23)<br>(7.53, 10.07)<br>(6.29, 9.3)<br>(0.34, 0.5) |
| 63 | LR size<br>$q=\phi^n/(k^n+\phi^n)$, $n=2$<br>$\tau=\tau_1+\tau_2\phi^n/(k^n+\phi^n)$, $n=1$ | $\alpha$<br>$\tau_1$<br>$\tau_2$<br>$k_q$<br>$k_d$ | 0.037<br>5<br>2.01<br>68.61<br>0.15 | (0.019, 0.073)<br>(1.1, 21.7)<br>(0.01, 146.24)<br>(NaN, NaN)<br>(8.63e-5, 73.3) |
| 64 | LR size<br>$q=\phi^n/(k^n+\phi^n)$, $n=2$<br>$\tau=\tau_1+\tau_2\phi^n/(k^n+\phi^n)$, $n=2$ | $\alpha$<br>$\tau_1$<br>$\tau_2$<br>$k_q$<br>$k_d$ | 0.037<br>3.41<br>2.85<br>7.95<br>0.02 | (0.0021, 0.065)<br>(NaN, NaN)<br>(NaN, NaN)<br>(0.88, 45.67)<br>(4.94e-14, 99.99) |
| 65 | LR size<br>$q=\phi^n/(k^n+\phi^n)$, $n=2$<br>$\tau=\tau_1+\tau_2\phi^n/(k^n+\phi^n)$, $n=3$ | $\alpha$<br>$\tau_1$<br>$\tau_2$<br>$k_q$<br>$k_d$ | 0.038<br>5.45<br>0.98<br>2.75<br>0.14 | (0.02, 0.08)<br>(2.91, 10.14)<br>(0.011, 65.21)<br>(0.03, 96.1)<br>(0.08, 0.25) |
| 66 | LR size<br>$q=\phi^n/(k^n+\phi^n)$, $n=3$<br>$\tau=\tau_1+\tau_2\phi^n/(k^n+\phi^n)$, $n=1$ | $\alpha$<br>$\tau_1$<br>$\tau_2$<br>$k_q$<br>$k_d$ | 0.12<br>3.93<br>3.91<br>0.06<br>0.01 | (NaN, NaN)<br>(NaN, NaN)<br>(NaN, NaN)<br>(NaN, NaN)<br>(NaN, NaN) |
| 67 | LR size<br>$q=\phi^n/(k^n+\phi^n)$, $n=3$<br>$\tau=\tau_1+\tau_2\phi^n/(k^n+\phi^n)$, $n=2$ | $\alpha$<br>$\tau_1$<br>$\tau_2$ | 0.03<br>4.41<br>5.44 | (0.034, 0.04)<br>(2.9, 6.710)<br>(51.54, 57.31) |
| | | $k_q$ | 1.27 | (NaN, NaN) |
| | | $k_d$ | 1.46 | (1.31, 1.63) |
| 68 | LR size<br>$q=\phi^n/(k^n+\phi^n)$, $n=3$<br>$\tau=\tau_1+\tau_2\phi^n/(k^n+\phi^n)$, $n=3$ | $\alpha$ | 0.03 | (0.01, 0.07) |
| | | $\tau_1$ | 2.08 | (NaN, NaN) |
| | | $\tau_2$ | 4.02E+00 | (NaN, NaN) |
| | | $k_q$ | 5.53 | (5.53, 5.53) |
| | | $k_d$ | 0.002 | (0.001, 0.006) |
| 69 | LR size<br>$q=\phi^n/(k^n+\phi^n)$, $n=1$<br>$\tau=\tau_1-\tau_2LR$ | $\alpha$ | 0.04 | (0.02, 0.79) |
| | | $\tau_1$ | 12.29 | (NaN, NaN) |
| | | $\tau_2$ | 1.83 | (NaN, NaN) |
| | | $k$ | 42.98 | (1.65, 97.12) |
| 70 | LR size<br>$q=\phi^n/(k^n+\phi^n)$, $n=2$<br>$\tau=\tau_1-\tau_2LR$ | $\alpha$ | 0.046 | (0.03, 0.07) |
| | | $\tau_1$ | 30.41 | (30.03, 30.79) |
| | | $\tau_2$ | 7.27 | (7.12, 7.43) |
| | | $k$ | 7.2 | (0.71, 45.3) |
| 71 | LR size<br>$q=\phi^n/(k^n+\phi^n)$, $n=3$<br>$\tau=\tau_1-\tau_2LR$ | $\alpha$ | 4.11 | (4.11, 4.11) |
| | | $\tau_1$ | 9.75 | (9.75, 9.75) |
| | | $\tau_2$ | 1.02 | (NaN, NaN) |
| | | $k$ | 0.017 | (NaN, NaN) |
| 72 | LR size<br>$q=\phi^n/(k^n+\phi^n)$, $n=1$<br>$\tau=\tau_1+\tau_2/LR$ | $\alpha$ | 0.08 | (0.01, 0.39) |
| | | $\tau_1$ | 4.65 | (NaN, NaN) |
| | | $\tau_2$ | 4.85 | (NaN, NaN) |
| | | $k$ | 0.14 | (0.009, 1.13) |
| 73 | LR size<br>$q=\phi^n/(k^n+\phi^n)$, $n=2$<br>$\tau=\tau_1+\tau_2\phi/LR$ | $\alpha$ | 0.04 | (0.022, 0.08) |
| | | $\tau_1$ | 3.37E-05 | (1.14e-14, 299.8) |
| | | $\tau_2$ | 21.22 | (16.73, 26.82) |
| | | $k$ | 1.99 | (NaN, NaN) |
| 74 | LR size<br>$q=\phi^n/(k^n+\phi^n)$, $n=3$<br>$\tau=\tau_1+\tau_2/LR$ | $\alpha$ | 0.04 | (0.02, 0.0*) |
| | | $\tau_1$ | 3.37E-05 | (1e-14, NaN) |
| | | $\tau_2$ | 21.23 | (16.73, 26.81) |
| | | $k$ | 10.2 | (NaN, NaN) |
| 75 | LR size<br>$q=\phi^n/(k^n+\phi^n)$, $n=1$<br>$\tau=\tau_1/LR$ | $\alpha$ | 0.049 | (0.02, 0.09) |
| | | $\tau_1$ | 21.89 | (18.64, 25.66) |
| | | $k$ | 1.99 | (1e-14, 2) |
| 76 | LR size<br>$q=\phi^n/(k^n+\phi^n)$, $n=2$<br>$\tau=\tau_1/LR$ | $\alpha$ | 0.044 | (0.02, 0.09) |
| | | $\tau_1$ | 21.23 | (16.75, 26.77) |
| | | $k$ | 1.94 | (1.49, 1.99) |
| 77 | LR size | $\alpha$ | NaN | (NaN, NaN) |
| | $q = \phi^n / (k^n + \phi^n), n=1$ | $\tau_1$ | NaN | (NaN, NaN) |
| | $\tau = \tau_1 + \tau_2(1 - LR^n / (k^n + LR^n)), n=1$ | $\tau_2$ | NaN | (NaN, NaN) |
| | | $k_q$ | NaN | (NaN, NaN) |
| | | $k_d$ | NaN | (NaN, NaN) |
| 78 | LR size | $\alpha$ | 0.04 | (0.04, 0.048) |
| | $q = \phi^n / (k^n + \phi^n), n=1$ | $\tau_1$ | 1.75 | (1.57, 1.88) |
| | $\tau = \tau_1 + \tau_2(1 - LR^n / (k^n + LR^n)), n=2$ | $\tau_2$ | 299.99 | (1e-14, 300) |
| | | $k_q$ | 3.83 | (2.17, 6.79) |
| | | $k_d$ | 0.4 | (0.4, 0.4) |
| 79 | LR size | $\alpha$ | | |
| | $q = \phi^n / (k^n + \phi^n), n=1$ | $\tau_1$ | | |
| | $\tau = \tau_1 + \tau_2(1 - LR^n / (k^n + LR^n)), n=3$ | $\tau_2$ | | |
| | | $k_q$ | | |
| | | $k_d$ | | |
| 80 | LR size | $\alpha$ | | |
| | $q = \phi^n / (k^n + \phi^n), n=2$ | $\tau_1$ | | |
| | $\tau = \tau_1 + \tau_2(1 - LR^n / (k^n + LR^n)), n=1$ | $\tau_2$ | | |
| | | $k_q$ | | |
| | | $k_d$ | | |
| 81 | LR size | $\alpha$ | 0.04 | (0.04, 0.04) |
| | $q = \phi^n / (k^n + \phi^n), n=2$ | $\tau_1$ | 1.79 | (1.67, 1.91) |
| | $\tau = \tau_1 + \tau_2(1 - LR^n / (k^n + LR^n)), n=2$ | $\tau_2$ | 299.9 | (NaN, NaN) |
| | | $k_q$ | 9.04 | (NaN, NaN) |
| | | $k_d$ | 0.4 | (0.39, 0.41) |
| 82 | LR size | $\alpha$ | NaN | (NaN, NaN) |
| | $q = \phi^n / (k^n + \phi^n), n=2$ | $\tau_1$ | NaN | (NaN, NaN) |
| | $\tau = \tau_1 + \tau_2(1 - LR^n / (k^n + LR^n)), n=3$ | $\tau_2$ | NaN | (NaN, NaN) |
| | | $k_q$ | NaN | (NaN, NaN) |
| | | $k_d$ | NaN | (NaN, NaN) |
| 83 | LR size | $\alpha$ | NaN | (NaN, NaN) |
| | $q = \phi^n / (k^n + \phi^n), n=3$ | $\tau_1$ | NaN | (NaN, NaN) |
| | $\tau = \tau_1 + \tau_2(1 - LR^n / (k^n + LR^n)), n=1$ | $\tau_2$ | NaN | (NaN, NaN) |
| | | $k_q$ | NaN | (NaN, NaN) |
| | | $k_d$ | NaN | (NaN, NaN) |
| 84 | LR size | $\alpha$ | NaN | (NaN, NaN) |
| | $q = \phi^n / (k^n + \phi^n), n=3$ | $\tau_1$ | NaN | (NaN, NaN) |
| | $\tau = \tau_1 + \tau_2(1 - LR^n / (k^n + LR^n)), n=2$ | $\tau_2$ | NaN | (NaN, NaN) |
| | | $k_q$ | NaN | (NaN, NaN) |
| | | $k_d$ | NaN | (NaN, NaN) |
| 85 | LR size<br>$q=\phi^n/(k^n+\phi^n)$, $n=3$<br>$\tau=\tau_1+\tau_2(1-LR^n/(k^n+LR^n))$, $n=3$ | $\alpha$ | NaN | (NaN, NaN) |
| | | $\tau_1$ | NaN | (NaN, NaN) |
| | | $\tau_2$ | NaN | (NaN, NaN) |
| | | $k_q$ | NaN | (NaN, NaN) |
| | | $k_d$ | NaN | (NaN, NaN) |
| 86 | LR size<br>$\tau=\tau_1+\tau_2\phi/LR$ | $\alpha(1-q)$ | 0.04 | (0.02, 0.08) |
| | | $\tau_1$ | 3.37E-05 | (3.37e-5, 3.37) |
| | | $\tau_2$ | 85.16 | (69.17, 103.1) |
| 87 | LR size<br>$\tau=\tau_1+\tau_2\phi-\tau_3LR$ | $\alpha(1-q)$ | 0.04 | (0.03, 0.07) |
| | | $\tau_1$ | 23.9 | (23.04, 24.83) |
| | | $\tau_2$ | 7.97 | (4.94, 12.67) |
| | | $\tau_3$ | 5.83 | (5.57, 6.09) |
| 88 | LR size<br>$q=\min(q_0\phi, 1)$<br>$\tau=\tau_1+\tau_2\phi/LR$ | $\alpha$ | 0.04 | (0.02, 0.08) |
| | | $q_0$ | 2.25E-07 | (NaN, NaN) |
| | | $\tau_1$ | 3.37E-05 | (1e-14, 300) |
| | | $\tau_2$ | 85.16 | (69.55, 102.71) |
| 89 | LR size<br>$q=\min(q_0\phi, 1)$<br>$\tau=\tau_1+\tau_2\phi-\tau_3LR$ | $\alpha$ | 0.15 | (0.13, 0.17) |
| | | $q_0$ | 1.93 | (1.11, 1.99) |
| | | $\tau_1$ | 20.6 | (20.21, 21.01) |
| | | $\tau_2$ | 49.3 | (47.25, 51.45) |
| | | $\tau_3$ | 8.02 | (7.75, 8.3) |
| 90 | LR size<br>$q=\phi^n/(k^n+\phi^n)$, $n=1$<br>$\tau=\tau_1+\tau_2\phi/LR$ | $\alpha$ | 0.049 | (0.024, 0.09) |
| | | $k$ | 1.99 | (NaN, NaN) |
| | | $\tau_1$ | 3.37E-05 | (1e-14, 300) |
| | | $\tau_2$ | 87.77 | (71.43, 106.1) |
| 91 | LR size<br>$q=\phi^n/(k^n+\phi^n)$, $n=2$<br>$\tau=\tau_1+\tau_2\phi/LR$ | $\alpha$ | 0.044 | (0.02, 0.08) |
| | | $k$ | 1.99 | (NaN, NaN) |
| | | $\tau_1$ | 3.37E-05 | (1e-14, NaN) |
| | | $\tau_2$ | 85.68 | (69.56, 103.85) |
| 92 | LR size<br>$q=\phi^n/(k^n+\phi^n)$, $n=3$<br>$\tau=\tau_1+\tau_2\phi/LR$ | $\alpha$ | 0.043 | (0.02, 0.08) |
| | | $k$ | 1.99 | (NaN, NaN) |
| | | $\tau_1$ | 3.37E-05 | (1e-14, 300) |
| | | $\tau_2$ | 85.27 | (69.25, 103.34) |
| 93 | LR size<br>$q=\phi^n/(k^n+\phi^n)$, $n=1$ | $\alpha$ | 0.19 | (0.16, 0.21) |
| | | $k$ | 0.1 | (0.09, 0.12) |
| | $\tau=\tau_1+\tau_2\phi-\tau_3LR$ | $\tau_1$ | 17.39 | (16.77, 18.04) |
| | | $\tau_2$ | 66.15 | (64.11, 68.24) |
| | | $\tau_3$ | 8.84 | (8.59, 9.1) |
| 94 | LR size<br>$q=\phi^n/(k^n+\phi^n)$, $n=2$<br>$\tau=\tau_1+\tau_2\phi-\tau_3LR$ | $\alpha$ | 0.05 | (0.02, 0.11) |
| | | $k$ | 0.68 | (0.02, 1.9) |
| | | $\tau_1$ | 28.77 | (28.29, 29.2) |
| | | $\tau_2$ | 7.94 | (5.46, 11.51) |
| | | $\tau_3$ | 7.3 | (7.02, 7.59) |
| 95 | LR size<br>$q=\phi^n/(k^n+\phi^n)$, $n=3$<br>$\tau=\tau_1+\tau_2\phi-\tau_3LR$ | $\alpha$ | 0.56 | (0.5, 0.64) |
| | | $k$ | 0.16 | (0.15, 0.17) |
| | | $\tau_1$ | 14.56 | (13.63, 15.55) |
| | | $\tau_2$ | 77.12 | (74.67, 79.62) |
| | | $\tau_3$ | 9.17 | (8.89, 9.46) |
| 96 | LR size<br>$\tau=\tau_1+\tau_2\phi-\tau_3LR+\tau_4\phi/LR$ | $\alpha(1-q)$ | 0.047 | (0.039, 0.05) |
| | | $\tau_1$ | 7.2 | (6.62, 7.82) |
| | | $\tau_2$ | 2.77 | (0.88, 8.51) |
| | | $\tau_3$ | 2.98 | (2.72, 3.26) |
| | | $\tau_4$ | 103.68 | (99.48, 107.96) |
| 97 | LR size<br>$\tau=\tau_1+\tau_2\phi+\tau_3LR+\tau_4\phi/LR$ | $\alpha(1-q)$ | 0.04 | (0.02, 0.08) |
| | | $\tau_1$ | 3.37E-05 | (3.37e-5, 3.37e-5) |
| | | $\tau_2$ | 3.37E-05 | (3.37e-5, 3.37e-5) |
| | | $\tau_3$ | 2.02E+01 | (NaN, NaN) |
| | | $\tau_4$ | 4.59 | (NaN, NaN) |
| 98 | LR size<br>$q=\min(q_0\phi, 1)$<br>$\tau=\tau_1+\tau_2\phi-\tau_3LR+\tau_4\phi/LR$ | $\alpha$ | 0.048 | (0.04, 0.05) |
| | | $q_0$ | 0.15 | (0.003, 1.62) |
| | | $\tau_1$ | 5.51 | (4.98, 6.09) |
| | | $\tau_2$ | 10.17 | (8.63, 11.97) |
| | | $\tau_3$ | 3.11 | (2.85, 3.37) |
| | | $\tau_4$ | 102.9 | (99.05, 106.83) |
| 99 | LR size<br>$q=\phi^n/(k^n+\phi^n)$, $n=1$<br>$\tau=\tau_1+\tau_2\phi-\tau_3LR+\tau_4\phi/LR$ | $\alpha$ | 0.05 | (0.046, 0.06) |
| | | $k$ | 1.38 | (0.09, 1.97) |
| | | $\tau_1$ | 7.29 | (6.7, 7.93) |
| | | $\tau_2$ | 0.58 | (0.002, 92.33) |
| | | $\tau_3$ | 2.89 | (2.62, 3.18) |
| | | $\tau_4$ | 108.9 | (104.52, 113.43) |
| 100 | LR size<br>$q=\phi^n/(k^n+\phi^n)$, $n=2$ | $\alpha$ | 0.05 | (0.04, 0.06) |
| | | $k$ | 0.98 | (0.24, 1.73) |
| | $\tau=\tau_1+\tau_2\phi-\tau_3LR+\tau_4\phi/LR$ | $\tau_1$ | 8.75 | (8.2, 9.33) |
| | | $\tau_2$ | 6.63 | (5.27, 8.35) |
| | | $\tau_3$ | 3.5 | (3.2, 3.76) |
| | | $\tau_4$ | 95.24 | (91.29, 99.28) |
| 101 | LR size<br>$q=\phi^n/(k^n+\phi^n)$, $n=3$<br>$\tau=\tau_1+\tau_2\phi-\tau_3LR+\tau_4\phi/LR$ | $\alpha$ | 0.05 | (0.04, 0.06) |
| | | $k$ | 0.77 | (0.56, 0.99) |
| | | $\tau_1$ | 9.17 | (8.63, 9.74) |
| | | $\tau_2$ | 13.12 | (11.6, 14.8) |
| | | $\tau_3$ | 3.95 | (3.71, 4.21) |
| | | $\tau_4$ | 85.74 | (81.91, 89.67) |
| 102 | LR size<br>$q=\min(q_0\phi, 1)$<br>$\tau=\tau_1+\tau_2\phi+\tau_3/LR+\tau_4\phi/LR$ | $\alpha$ | 0.05 | (0.04, 0.09) |
| | | $q_0$ | 0.01 | (8.38e-10, 1.99) |
| | | $\tau_1$ | 3.37E-05 | (3.37e-5, 3.37e-5) |
| | | $\tau_2$ | 3.37E-05 | (3.37e-5, 3.37e-5) |
| | | $\tau_3$ | 19.6 | (NaN, NaN) |
| | | $\tau_4$ | 7.32E+00 | (NaN, NaN) |
| 103 | LR size<br>$q=\phi^n/(k^n+\phi^n)$, $n=1$<br>$\tau=\tau_1+\tau_2\phi+\tau_3/LR+\tau_4\phi/LR$ | $\alpha$ | 4.00E-02 | (0.02, 0.09) |
| | | $k$ | 1.99 | (NaN, NaN) |
| | | $\tau_1$ | 4.28E-05 | (1e-14, NaN) |
| | | $\tau_2$ | 3.37E-05 | (NaN, NaN) |
| | | $\tau_3$ | 1.95E+01 | (NaN, NaN) |
| | | $\tau_4$ | 7.87 | (NaN, NaN) |
| 104 | LR size<br>$q=\phi^n/(k^n+\phi^n)$, $n=2$<br>$\tau=\tau_1+\tau_2\phi+\tau_3/LR+\tau_4\phi/LR$ | $\alpha$ | NaN | (NaN, NaN) |
| | | $k$ | NaN | (NaN, NaN) |
| | | $\tau_1$ | NaN | (NaN, NaN) |
| | | $\tau_2$ | NaN | (NaN, NaN) |
| | | $\tau_3$ | NaN | (NaN, NaN) |
| | | $\tau_4$ | NaN | (NaN, NaN) |
| 105 | LR size<br>$q=\phi^n/(k^n+\phi^n)$, $n=3$<br>$\tau=\tau_1+\tau_2\phi+\tau_3/LR+\tau_4\phi/LR$ | $\alpha$ | NaN | (NaN, NaN) |
| | | $k$ | NaN | (NaN, NaN) |
| | | $\tau_1$ | NaN | (NaN, NaN) |
| | | $\tau_2$ | NaN | (NaN, NaN) |
| | | $\tau_3$ | NaN | (NaN, NaN) |
| | | $\tau_4$ | NaN | (NaN, NaN) |
| 106 | LR size<br>$\tau=\tau_1+\tau_2\phi-\tau_3LR+\tau_4\phi*LR$ | $\alpha(1-q)$ | 0.04 | (0.03, 0.06) |
| | | $\tau_1$ | 27.48 | (26.99, 27.98) |
| | | $\tau_2$ | 45.62 | (42.41, 49.04) |
| | | $\tau_3$ | 3.4 | (3.22, 3.73) |
| | | $\tau_4$ | -34.14 | (-35.68, -32.58) |
| 107 | LR size<br>$q=\min(q_0\phi, 1)$<br>$\tau=\tau_1+\tau_2\phi-\tau_3LR+\tau_4\phi*LR$ | $\alpha$ | 0.14 | (0.12, 0.17) |
| | | $q_0$ | 1.9 | (1.4, 1.98) |
| | | $\tau_1$ | 21.21 | (20.8, 21.63) |
| | | $\tau_2$ | 46.66 | (44.68, 48.72) |
| | | $\tau_3$ | 8.2 | (8.02, 8.3) |
| | | $\tau_4$ | 0.82 | (-0.06, 1.71) |
| 108 | LR size<br>$q=\phi^n/(k^n+\phi^n)$, $n=1$<br>$\tau=\tau_1+\tau_2\phi-\tau_3LR+\tau_4\phi*LR$ | $\alpha$ | 1.19 | (NaN, NaN) |
| | | $k$ | 0.014 | (NaN, NaN) |
| | | $\tau_1$ | 28.32 | (23.17, 34.47) |
| | | $\tau_2$ | 15.18 | (NaN, NaN) |
| | | $\tau_3$ | 11.69 | (9.47, 14.4) |
| | | $\tau_4$ | 13.94 | (NaN, NaN) |
| 109 | LR size<br>$q=\phi^n/(k^n+\phi^n)$, $n=2$<br>$\tau=\tau_1+\tau_2\phi-\tau_3LR+\tau_4\phi*LR$ | $\alpha$ | 0.1 | (0.09, 0.12) |
| | | $k$ | 0.32 | (0.28, 0.37) |
| | | $\tau_1$ | 7.19 | (6.2, 8.33) |
| | | $\tau_2$ | 103.01 | (100.14, 105.92) |
| | | $\tau_3$ | 10.39 | (9.95, 10.84) |
| | | $\tau_4$ | 1.15 | (-0.05, 2.35) |
| 110 | LR size<br>$q=\phi^n/(k^n+\phi^n)$, $n=3$<br>$\tau=\tau_1+\tau_2\phi-\tau_3LR+\tau_4\phi*LR$ | $\alpha$ | 0.04 | (NaN, NaN) |
| | | $k$ | 1.19 | (NaN, NaN) |
| | | $\tau_1$ | 22.28 | (21.56, 23.03) |
| | | $\tau_2$ | 34.9 | (32.9, 36.9) |
| | | $\tau_3$ | 5.49 | (5.23, 5.76) |
| | | $\tau_4$ | -7.62 | (-8.54, -6.71) |

| AIC | $\Delta AIC$ |
| --- | --- |
| 45.05 | 0 |
| 47.05 | 2 |
| 47.05 | 2 |
| 47.05 | 2 |
| 49.05 | 4 |
| 49.05 | 4 |
| 45.51 | 0.46 |
| 47.51 | 2.46 |
| 5.43 | -45.05<br>-39.62 |
| 47.52 | 2.47 |
| 49.51 | 4.46 |
| 49.51 | 4.46 |
| 47.05 | 2 |
| 49.05 | 4 |
| 47.51 | 2.46 |
| 49.51 | 4.46 |
| 47.05 | 2 |
| 49.05 | 4 |
| 47.53 | 2.48 |
| 49.52 | 4.47 |
| 47.05 | 2 |
| 47.05 | 2 |
| 47.05 | 2 |
| 49.05 | 4 |
| 47.53 | 2.48 |
| 47.52 | 2.47 |
| 49.51 | 4.46 |
| 49.05 | 4 |
| 49.05 | 4 |
| 49.05 | 4 |
| 50.98 | 5.93 |
| 51.04 | 5.99 |
| NaN |  |
| 45.64 | 0.59 |
| NaN |  |
| 51.51 | 6.46 |
| 51.51 | 6.46 |
| 51.51 | 6.46 |
| 44.18 | -0.87 |
| 45.69 | 0.64 |
| 43.69 | -1.36 |
| 46.1 | 1.05 |
| 45.69 | 0.64 |
| 49.51 | 4.46 |
| 49.51 | 4.46 |
| 49.51 | 4.46 |
| NaN |  |
| 47.76 | 2.71 |
| NaN |  |

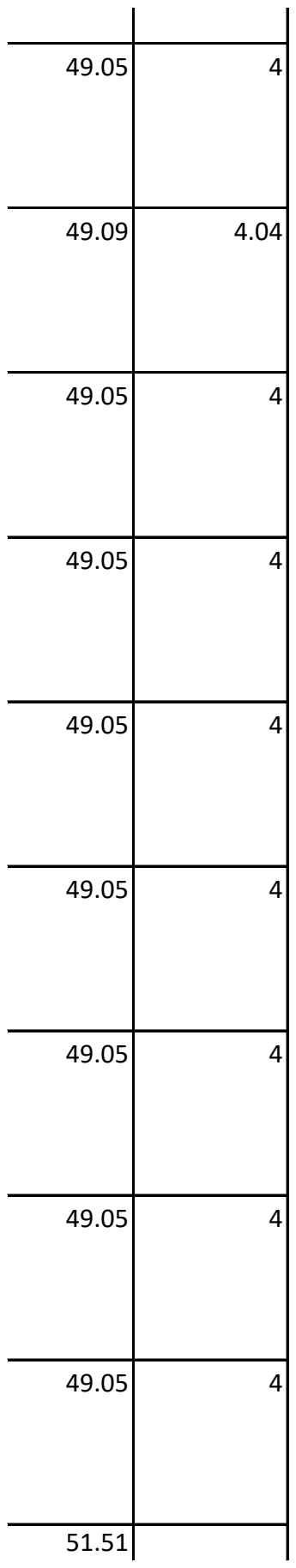

|  |  |
| --- | --- |
| 51.51 | 6.46 |
| 51.53 | 6.48 |
| 51.51 | 6.46 |
| 51.51 | 6.46 |
| 51.51 | 6.46 |
| 73.57 | 28.52 |
| 51.53 | 6.48 |
| 51.51 | 6.46 |
| 48.58 | 3.53 |
| 46.17 | 1.12 |
| 73.96 | 28.91 |
| 50.11 | 5.06 |
| 47.69 | 2.64 |
| 47.69 | 2.64 |
| 45.79 | 0.74 |
| 45.69 | 0.64 |
| NaN |  |
| 47.44 | 2.39 |
| 47.64 | 2.59 |
| NaN |  |
| NaN |  |
| NaN |  |
| NaN |  |
| 45.69 | 0.64 |
| 46.34 | 1.29 |
| 47.69 | 2.64 |
| 47.53 | 2.48 |
| 47.62 | 2.57 |
| 47.69 | 2.64 |
| 47.69 | 2.64 |
| 47.78 | 2.73 |
| 48.14 | 3.09 |
| 47.11 | 2.06 |
| 47.67 | 2.62 |
| 49.69 | 4.64 |
| 49.67 | 4.62 |
| 49.67 | 4.62 |
| 49.72 | 4.67 |
| 49.75 | 4.7 |
| 51.69 | 6.64 |
| 51.7 | 6.65 |
| NaN |  |
| NaN |  |
| 47.42 | 2.37 |
| 49.49 | 4.44 |
| 48.41 | 3.36 |
| 49.92 | 4.87 |
| 50.17 | 5.12 |

| Model No | Model Description | Parameters | Parameter | 95% CI | AIC |
| --- | --- | --- | --- | --- | --- |
| 1 | Null | $\alpha(1-q)$<br>$\tau$ | 0.012<br>17 | (0.004, 0.03)<br>(16.3, 17.71) | 35.42 |
| 2 | $q=\min(q_0\phi, 1)$ | $\alpha$<br>$q_0$<br>$\tau$ | 0.02<br>1.99<br>17 | (0.001, 0.33)<br>(1e-14, NaN)<br>(16.36, 17.63) | 37.13 |
| 3 | $\tau=\tau_1+(\tau_2-\tau_1)\phi$ | $\alpha(1-q)$<br>$\tau_1$<br>$\tau_2$ | 0.01<br>18.54<br>11.48 | (0.004, 0.03)<br>(17.92, 18.99)<br>(9.46, 13.73) | 37.24 |
| 4 | $\tau=\tau_1+(\tau_2-\tau_1)\phi^2$ | $\alpha(1-q)$<br>$\tau_1$<br>$\tau_2$ | 0.02<br>17.58<br>8.47 | (0.004, 0.03)<br>(17.05, 18.03)<br>(4.58, 14.32) | 37.24 |
| 5 | $q=\min(q_0\phi, 1)$<br>$\tau=\tau_1+(\tau_2-\tau_1)\phi$ | $\alpha$<br>$q_0$<br>$\tau_1$<br>$\tau_2$ | 0.012<br>1.99<br>10.45<br>18.87 | (0.005, 0.03)<br>(NaN, NaN)<br>(10.19, 10.71)<br>(18.01, 19.74) | 39.24 |
| 6 | $q=\min(q_0\phi, 1)$<br>$\tau=\tau_1+(\tau_2-\tau_1)\phi^2$ | $\alpha$<br>$q_0$<br>$\tau_1$<br>$\tau_2$ | 0.012<br>0.05<br>16.98<br>17.21 | (0.003, 0.04)<br>(1e-14, NaN)<br>(16.42, 17.46)<br>(12.15, 22.67) | 39.24 |
| 7 | LR size | $\alpha(1-q)$<br>$\tau$ | 0.005<br>17.21 | (0.002, 0.015)<br>(10, 25.25) | 35.19 |
| 8 | LR size<br>$q=\min(q_0\phi, 1)$ | $\alpha$<br>$q_0$<br>$\tau$ | 0.016<br>1.99<br>17 | (0.0006, 0.17)<br>(1e-14, NaN)<br>(16.39, 17.61) | 37.02 |
| 9 | LR size<br>$\tau=\tau_1+(\tau_2-\tau_1)\phi$ | $\alpha(1-q)$<br>$\tau_1$<br>$\tau_2$ | 0.005<br>10.15<br>39.99 | (0.002, 0.013)<br>(NaN, NaN)<br>(1e-14, NaN) | 37.31 |
| 10 | LR size<br>$\tau=\tau_1+(\tau_2-\tau_1)\phi^2$ | $\alpha(1-q)$<br>$\tau_1$<br>$\tau_2$ | 0.005<br>18.14<br>0.71 | (0.002, 0.013)<br>(17.73, 19.1)<br>(0.05, 18.66) | 37.19 |
| 11 | LR size<br>$q=\min(q_0\phi, 1)$<br>$\tau=\tau_1+(\tau_2-\tau_1)\phi$ | $\alpha$<br>$q_0$<br>$\tau_1$<br>$\tau_2$ | 0.01<br>1.85E+00<br>9.78<br>52.86 | (0.003, 0.029)<br>(0.18, 1.99)<br>(8.06, 11.82)<br>(44.17, 61.38) | 39.1 |
| 12 | LR size<br>$q=\min(q_0\phi, 1)$ | $\alpha$<br>$q_0$ | 0.01<br>1.99 | (0.004, 0.02)<br>(1e-14, 2) | 39.08 |
| | $\tau=\tau_1+(\tau_2-\tau_1)\phi^2$ | $\tau_1$ | 17.93 | (17.21, 18.68) | |
| | | $\tau_2$ | 0.51 | (0.0004, 86.9) | |
| 13 | $\tau=\tau_1+\tau_2\phi$ | $\alpha(1-q)$ | 0.12 | (0.004, 0.034) | 37.24 |
| | | $\tau_1$ | 16.78 | (16.26, 17.23) | |
| | | $\tau_2$ | 0.99 | (0.3, 3.12) | |
| 14 | $q=\min(q_0\phi, 1)$<br>$\tau=\tau_1+\tau_2\phi$ | $\alpha$ | 0.01 | (0.003, 0.03) | 39.47 |
| | | $q_0$ | 0.0001 | (1e-14, NaN) | |
| | | $\tau_1$ | 14.22 | (12.04, 16) | |
| | | $\tau_2$ | 12.65 | (5.67, 22.67) | |
| 15 | LR size<br>$\tau=\tau_1+\tau_2\phi$ | $\alpha(1-q)$ | 0.005 | (0.0029, 0.01) | 37.19 |
| | | $\tau_1$ | 11.61 | (10.96, 12.28) | |
| | | $\tau_2$ | 25.4 | (23.23, 27.44) | |
| 16 | LR size<br>$q=\min(q_0\phi, 1)$<br>$\tau=\tau_1+\tau_2\phi$ | $\alpha$ | 0.001 | (0.004, 0.027) | 39.08 |
| | | $q_0$ | 1.99 | (1e-14, 2) | |
| | | $\tau_1$ | 15.33 | (14.63, 16.06) | |
| | | $\tau_2$ | 11.74 | (9.29, 14.73) | |
| 17 | $\tau=\tau_1+(\tau_2+\tau_1)\phi$ | $\alpha(1-q)$ | 0.012 | (0.004, 0.03) | 37.26 |
| | | $\tau_1$ | 11.52 | (NaN, NaN) | |
| | | $\tau_2$ | 13.33 | (NaN, NaN) | |
| 18 | $q=\min(q_0\phi, 1)$<br>$\tau=\tau_1+(\tau_2+\tau_1)\phi$ | $\alpha$ | 0.01 | (0.004, 0.03) | 39.3 |
| | | $q_0$ | 1.99 | (NaN, NaN) | |
| | | $\tau_1$ | 14.14 | (6.43, 18.49) | |
| | | $\tau_2$ | 4.50E-06 | (1e-14, NaN) | |
| 19 | LR size<br>$\tau=\tau_1+(\tau_2+\tau_1)\phi$<br>$\tau=\tau_1+(\tau_2+\tau_1)\phi$ | $\alpha(1-q)$ | 0.005 | (0.002, 0.01) | 37.19 |
| | | $\tau_1$ | 11.39 | (10.84, 11.96) | |
| | | $\tau_2$ | 14.97 | (12.93, 17.12) | |
| 20 | LR size<br>$q=\min(q_0\phi, 1)$<br>$\tau=\tau_1+(\tau_2+\tau_1)\phi$ | $\alpha$ | 0.006 | (0.003, 0.01) | 39.19 |
| | | $q_0$ | 0.6 | (0.31, 0.98) | |
| | | $\tau_1$ | 8.93 | (8.37, 9.54) | |
| | | $\tau_2$ | 30.16 | (27.49, 32.97) | |
| 21 | $q=\phi^n/(k^n+\phi^n)$, $n=1$ | $\alpha(1-q)$ | 3.2 | (0.0005, 4.99) | 36.52 |
| | | $k$ | 0.0005 | (1.67e-5, 0.02) | |
| | | $\tau$ | 17 | (16.55, 17.44) | |
| 22 | $q=\phi^n/(k^n+\phi^n)$, $n=2$ | $\alpha(1-q)$ | 1.31 | (3.15e-9, 5) | 36.14 |
| | | $k$ | 0.016 | (9.16e-6, 26.8) | |
| | | $\tau$ | 18.97 | (9.85, 28.54) | |
| 23 | $q=\phi^n/(k^n+\phi^n)$, $n=3$ | $\alpha(1-q)$ | 2.06 | (9.29e-9, 5) | 36.14 |
|  |  | k | 0.03 | (0.0003, 2.41) |  |
| | | $\tau$ | 20.04 | (13.9, 26.1) | |
| 24 | $q=q_0+q_1\phi$ | $\alpha$ | 0.16 | (0.033, 0.69) | 38.69 |
| | | $q_0$ | 0.66 | (0.35, 0.87) | |
| | | $q_1$ | 0.96 | (8.91e-10, 1) | |
| | | $\tau$ | 17.03 | (15.49, 18.61) | |
| 25 | LR size<br>$q=\phi^n/(k^n+\phi^n)$, $n=1$ | $\alpha$ | 0.02 | (NaN, NaN) | 36.96 |
|  |  | k | 0.06 | (NaN, NaN) |  |
| | | $\tau$ | 17.07 | (16.39, 17.76) | |
| 26 | LR size<br>$q=\phi^n/(k^n+\phi^n)$, $n=2$ | $\alpha$ | 0.6 | (0.25, 1.35) | 36.11 |
|  |  | k | 0.01 | (0.005, 0.04) |  |
| | | $\tau$ | 19.11 | (15.3, 22.99) | |
| 27 | LR size<br>$q=\phi^n/(k^n+\phi^n)$, $n=3$ | $\alpha$ | 0.27 | (NaN, NaN) | 36.11 |
|  |  | k | 0.045 | (0.02, 0.09) |  |
| | | $\tau$ | 18.96 | (15.41, 22.58) | |
| 28 | LR size<br>$q=q_0+q_1\phi$ | $\alpha$ | 0.044 | (NaN, NaN) | 38.64 |
| | | $q_0$ | 0.49 | | |
| | | $q_1$ | 1.43 | | |
| | | $\tau$ | 17 | (16.43, 21.76) | |
| 29 | $\tau=\tau_1+\tau_2\phi^n/(k^n+\phi^n)$, $n=1$ | $\alpha(1-q)$ | 0.01 | (0.004, 0.03) | 39.24 |
| | | $\tau_1$ | 12.1 | (11.46, 12.72) | |
| | | $\tau_2$ | 7.1 | (6.27, 8.02) | |
|  |  | k | 0.06 | (0.04, 0.09) |  |
| 30 | $\tau=\tau_1+\tau_2\phi^n/(k^n+\phi^n)$, $n=2$ | $\alpha(1-q)$ | 0.01 | (0.004, 0.03) | 39.24 |
| | | $\tau_1$ | 12.1 | (11.47, 12.72) | |
| | | $\tau_2$ | 6.16 | (5.43, 6.97) | |
|  |  | k | 0.06 | (0.04, 0.09) |  |
| 31 | $q=\phi^n/(k^n+\phi^n)$, $n=3$ | $\alpha(1-q)$ | 0.01 | (0.004, 0.03) | 39.24 |
| | | $\tau_1$ | 12.1 | (11.49, 12.72) | |
| | | $\tau_2$ | 6.08 | (5.36, 6.88) | |
|  |  | k | 0.07 | (0.04, 0.1) |  |
| 32 | $q=\min(q_0\phi, 1)$<br>$q=\phi^n/(k^n+\phi^n)$, $n=1$ | $\alpha$ | 0.01 | (0.003, 0.05) | 41.24 |
| | | $q_0$ | 0.12 | (1e-14, 2) | |
| | | $\tau_1$ | 10.91 | (10.2, 11.66) | |
| | | $\tau_2$ | 6.23 | (5.61, 6.92) | |
|  |  | k | 0.002 | (0.0002, 0.011) |  |
| 33 | $q=\min(q_0\phi, 1)$<br>$\tau=\tau_1+\tau_2\phi^n/(k^n+\phi^n)$, $n=2$ | $\alpha$ | 0.01 | (0.01, 0.01) | 41.4 |
| | | $q_0$ | 0.9 | (0.89, 0.9) | |
| | | $\tau_1$ | 11.27 | (11.21, 11.32) | |
| | | $\tau_2$ | 5.21 | (5.18, 5.23) | |
|  |  | k | 0.01 | (0.1, 0.01) |  |
| 33 | $q=\min(q_0\phi, 1)$<br>$\tau=\tau_1+\tau_2\phi^n/(k^n+\phi^n), n=3$ | $\alpha$ | 0.013 | (0.008, 0.02) | 41.21 |
| | | $q_0$ | 0.78 | (1e-7, 1.99) | |
| | | $\tau_1$ | 12.1 | (11.45, 11.79) | |
| | | $\tau_2$ | 4.9 | (4.67, 5.13) | |
|  |  | k | 0.0005 | (0.0002, 0.001) |  |
| 34 | LR size<br>$\tau=\tau_1+\tau_2\phi^n/(k^n+\phi^n), n=1$ | $\alpha(1-q)$ | 0.005 | (0.002, 0.01) | 39.19 |
| | | $\tau_1$ | 10.43 | (9.73, 11.18) | |
| | | $\tau_2$ | 9.02 | (8.2, 9.92) | |
|  |  | k | 0.045 | (0.03, 0.06) |  |
| 35 | LR size<br>$\tau=\tau_1+\tau_2\phi^n/(k^n+\phi^n), n=2$ | $\alpha(1-q)$ | 0.005 | (0.003, 0.009) | 39.19 |
| | | $\tau_1$ | 10.23 | (9.69, 10.8) | |
| | | $\tau_2$ | 10.02 | (9.36, 10.7) | |
|  |  | k | 0.099 | (0.083, 0.11) |  |
| 36 | LR size<br>$\tau=\tau_1+\tau_2\phi^n/(k^n+\phi^n), n=3$ | $\alpha(1-q)$ | 0.005 | (0.003, 0.01) | 39.19 |
| | | $\tau_1$ | 11.12 | (10.4, 11.89) | |
| | | $\tau_2$ | 8.12 | (7.3, 9.02) | |
|  |  | k | 0.092 | (0.07, 0.11) |  |
| 37 | LR size<br>$q=\min(q_0\phi, 1)$<br>$\tau=\tau_1+\tau_2\phi^n/(k^n+\phi^n), n=1$ | $\alpha$ | 1.77 | (0.07, 1.99) | 41.19 |
| | | $q_0$ | 10.35 | (NaN, NaN) | |
| | | $\tau_1$ | 7.05 | (1.12, 40.23) | |
| | | $\tau_2$ | 0.01 | (6.29e-7, 61.38) | |
|  |  | k | 0.01 | (NaN, NaN) |  |
| 38 | LR size<br>$q=\min(q_0\phi, 1)$<br>$\tau=\tau_1+\tau_2\phi^n/(k^n+\phi^n), n=2$ | $\alpha$ | 1.99 | (1e-14, 2) | 41.04 |
| | | $q_0$ | 15.74 | (15.06, 16.46) | |
| | | $\tau_1$ | 1.32 | (1.03, 1.07) | |
| | | $\tau_2$ | 0.009 | (0.003, 0.02) | |
|  |  | k | 0.01 | (0.001, 0.1) |  |
| 39 | LR size<br>$q=\min(q_0\phi, 1)$<br>$\tau=\tau_1+\tau_2\phi^n/(k^n+\phi^n), n=3$ | $\alpha$ | 8.64E-05 | (NaN, NaN) | 51.51 |
| | | $q_0$ | 4.62 | (3.85, 5.55) | |
| | | $\tau_1$ | 1.49 | (0.19, 11.27) | |
| | | $\tau_2$ | 0.01 | (0.0006, 0.16) | |
|  |  | k | 0.03 | (0.019, 0.07) |  |
| 40 | LR size<br>$\tau=\tau_1-\tau_2\text{LR}$ | $\alpha(1-q)$ | 0.005 | (0.002, 0.01) | 37.15 |
| | | $\tau_1$ | 20.73 | (19.94, 21.55) | |
| | | $\tau_2$ | 1.37 | (1.18, 1.6) | |
| 41 | LR size<br>$\tau=\tau_1+\tau_2/\text{LR}$ | $\alpha(1-q)$<br>$\tau_1$<br>$\tau_2$ | 0.005<br>4.96<br>29.63 | (0.002, 0.01)<br>(4.39, 5.61)<br>(27.93, 31.43) | 37.02 |
| 42 | LR size<br>$\tau=\tau_1/\text{LR}$ | $\alpha(1-q)$<br>$\tau_1$ | 0.005<br>40.71 | (0.002, 0.01)<br>(NaN, NaN) | 35.05 |
| 43 | LR size<br>$q=\min(q_0\phi, 1)$<br>$\tau=\tau_1-\tau_2\text{LR}$ | $\alpha$<br>$q_0$<br>$\tau_1$<br>$\tau_2$ | 0.01<br>1.99<br>30.44<br>5.27 | (0.004, 0.02)<br>(1e-14, NaN)<br>(29.51, 31.39)<br>(4.96, 5.6) | 38.9 |
| 44 | LR size<br>$q=\min(q_0\phi, 1)$<br>$\tau=\tau_1/\text{LR}$ | $\alpha$<br>$q_0$<br>$\tau_1$ | 0.01<br>1.99<br>41.4 | (0.004, 0.02)<br>(1e-14, 2)<br>(39.7, 43.1) | 36.8 |
| 45 | LR size<br>$\tau=\tau_1+\tau_2(1-\text{LR}^n/(k^n+\text{LR}^n)), n=1$ | $\alpha(1-q)$<br>$\tau_1$<br>$\tau_2$<br>$k$ | NaN<br>NaN<br>NaN<br>NaN | (NaN, NaN)<br>(NaN, NaN)<br>(NaN, NaN)<br>(NaN, NaN) | NaN |
| 46 | LR size<br>$\tau=\tau_1+\tau_2(1-\text{LR}^n/(k^n+\text{LR}^n)), n=2$ | $\alpha(1-q)$<br>$\tau_1$<br>$\tau_2$<br>$k$ | 0.005<br>16.48<br>299.9<br>0.12 | (0.002, 0.01)<br>(15.8, 17.1)<br>(NaN, NaN)<br>(0.1, 0.13) | 39.1 |
| 47 | LR size<br>$\tau=\tau_1+\tau_2(1-\text{LR}^n/(k^n+\text{LR}^n)), n=3$ | $\alpha(1-q)$<br>$\tau_1$<br>$\tau_2$<br>$k$ | 0.005<br>12.7<br>4.56<br>27.66 | (0.002, 0.01)<br>(12, 13.47)<br>(4.01, 5.17)<br>(3.03e-5, 99.9) | 39.19 |
| 48 | LR size<br>$q=\min(q_0\phi, 1)$<br>$\tau=\tau_1+\tau_2(1-\text{LR}^n/(k^n+\text{LR}^n)), n=1$ | $\alpha$<br>$q_0$<br>$\tau_1$<br>$\tau_2$<br>$k$ | NaN<br>NaN<br>NaN<br>NaN<br>NaN | (NaN, NaN)<br>(NaN, NaN)<br>(NaN, NaN)<br>(NaN, NaN)<br>(NaN, NaN) | NaN |
| 49 | LR size<br>$q=\min(q_0\phi, 1)$<br>$\tau=\tau_1+\tau_2(1-\text{LR}^n/(k^n+\text{LR}^n)), n=2$ | $\alpha$<br>$q_0$<br>$\tau_1$<br>$\tau_2$<br>$k$ | 0.01<br>2.87<br>14.02<br>2999.9<br>0.12 | (0.002, 0.03)<br>(NaN, NaN)<br>(NaN, NaN)<br>(1e-14, NaN)<br>(NaN, NaN) | 42.1 |
| 50 | LR size<br>$q=\min(q_0\phi, 1)$<br>$\tau=\tau_1+\tau_2(1-\text{LR}^n/(k^n+\text{LR}^n)), n=3$ | $\alpha$<br>$q_0$<br>$\tau_1$<br>$\tau_2$ | NaN<br>NaN<br>NaN<br>NaN | (NaN, NaN)<br>(NaN, NaN)<br>(NaN, NaN)<br>(NaN, NaN) | NaN<br>NaN<br>NaN<br>NaN |

|  |  | k | NaN | (NaN, NaN) | NaN |
| --- | --- | --- | --- | --- | --- |
| 51 | $q=\phi n/(kn+\phi n)$, $n=1$<br>$\tau=\tau_1+\tau_2\phi^n/(k^n+\phi^n)$, $n=1$ | $\alpha$<br>$\tau_1$<br>$\tau_2$<br>k | 0.03<br>8.21<br>18.05<br>0.12 | (0.013, 0.07)<br>(7.57, 8.97)<br>(16.6, 19.57)<br>(0.1, 0.15) | 39.15 |
| 52 | $q=\phi^n/(k^n+\phi^n)$, $n=1$<br>$\tau=\tau_1+\tau_2\phi^n/(k^n+\phi^n)$, $n=2$ | $\alpha$<br>$\tau_1$<br>$\tau_2$<br>k | 3.26<br>4.83E+00<br>12.74<br>0.0006 | (NaN, NaN)<br>(0.15, 87.1)<br>(3.14, 44.9)<br>(NaN, NaN) | 38.59 |
| 53 | $q=\phi^n/(k^n+\phi^n)$, $n=1$<br>$\tau=\tau_1+\tau_2\phi^n/(k^n+\phi^n)$, $n=3$ | $\alpha$<br>$\tau_1$<br>$\tau_2$<br>k | 0.83<br>5.47<br>12.08<br>0.002 | (NaN, NaN)<br>(4, 7.45)<br>(10.5, 13.92)<br>(NaN, NaN) | 38.62 |
| 54 | $q=\phi^n/(k^n+\phi^n)$, $n=2$<br>$\tau=\tau_1+\tau_2\phi^n/(k^n+\phi^n)$, $n=1$ | $\alpha$<br>$\tau_1$<br>$\tau_2$<br>k | 19.9<br>4.36<br>16.26<br>0.004 | (1e-14, 20)<br>(NaN, NaN)<br>(NaN, NaN)<br>(0.016, 0.01) | 38.2 |
| 55 | $q=\phi^n/(k^n+\phi^n)$, $n=2$<br>$\tau=\tau_1+\tau_2\phi^n/(k^n+\phi^n)$, $n=2$ | $\alpha$<br>$\tau_1$<br>$\tau_2$<br>k | 0.18<br>4.63<br>21.81<br>0.05 | (0.07, 0.45)<br>(2.97, 7.19)<br>(18.9, 25.06)<br>(0.021, 0.11) | 38.54 |
| 56 | $q=\phi^n/(k^n+\phi^n)$, $n=2$<br>$\tau=\tau_1+\tau_2\phi^n/(k^n+\phi^n)$, $n=3$ | $\alpha$<br>$\tau_1$<br>$\tau_2$<br>k | 0.01<br>14.6<br>199.9<br>1.24 | (0.004, 0.02)<br>(NaN, NaN)<br>(1e-14, NaN)<br>(NaN, NaN) | 39.58 |
| 57 | $q=\phi^n/(k^n+\phi^n)$, $n=2$<br>$\tau=\tau_1+\tau_2\phi^n/(k^n+\phi^n)$, $n=1$ | $\alpha$<br>$\tau_1$<br>$\tau_2$<br>k | 1.51<br>11.69<br>13.59<br>0.035 | (1e-14, 20)<br>(4.48, 28.81)<br>(NaN, NaN)<br>(2.42e-9, 199.2) | 38.1 |
| 58 | $q=\phi^n/(k^n+\phi^n)$, $n=3$<br>$\tau=\tau_1+\tau_2\phi^n/(k^n+\phi^n)$, $n=2$ | $\alpha$<br>$\tau_1$<br>$\tau_2$<br>k | 0.34<br>11.97<br>13.96<br>0.06 | (0.0001, 19.65)<br>(1.64, 65.77)<br>(0.43, 143.79)<br>(0.003, 1.26) | 38.31 |
| 59 | $q=\phi^n/(k^n+\phi^n)$, $n=3$<br>$\tau=\tau_1+\tau_2\phi^n/(k^n+\phi^n)$, $n=3$ | $\alpha$<br>$\tau_1$<br>$\tau_2$<br>k | 0.24<br>12.11<br>13.93<br>0.06 | (0.0003, 17.77)<br>(2.29, 52.8)<br>(0.65, 126.5)<br>(0.005, 0.93) | 38.36 |
| 60 | LR size | $\alpha$ | 0.01 | (0.004, 0.03) | 41.1 |
| | $q=\phi^n/(k^n+\phi^n)$, $n=1$<br>$\tau=\tau_1+\tau_2\phi^n/(k^n+\phi^n)$, $n=1$ | $\tau_1$<br>$\tau_2$<br>$k_q$<br>$k_d$ | 12.99<br>4.51<br>0.2<br>0.009 | (12.41, 13.6)<br>(4.07, 5)<br>(0.11, 0.36)<br>(0.004, 0.02) | |
| 61 | LR size<br>$q=\phi^n/(k^n+\phi^n)$, $n=2$<br>$\tau=\tau_1+\tau_2\phi^n/(k^n+\phi^n)$, $n=2$ | $\alpha$<br>$\tau_1$<br>$\tau_2$<br>$k_q$<br>$k_d$ | 0.018<br>14.22<br>2.87<br>0.12<br>0.001 | (0.013, 0.03)<br>(8.81, 22.69)<br>(0.27, 28.06)<br>(0.0001, 52.89)<br>(3.18e-5, 0.09) | 41.33 |
| 62 | LR size<br>$q=\phi^n/(k^n+\phi^n)$, $n=1$<br>$\tau=\tau_1+\tau_2\phi^n/(k^n+\phi^n)$, $n=3$ | $\alpha$<br>$\tau_1$<br>$\tau_2$<br>$k_q$<br>$k_d$ | 0.01<br>14.02<br>3.04<br>0.16<br>0.007 | (NaN, NaN)<br>(13.71, 14.33)<br>(NaN, NaN)<br>(NaN, NaN)<br>(NaN, NaN) | 44.46 |
| 63 | LR size<br>$q=\phi^n/(k^n+\phi^n)$, $n=2$<br>$\tau=\tau_1+\tau_2\phi^n/(k^n+\phi^n)$, $n=1$ | $\alpha$<br>$\tau_1$<br>$\tau_2$<br>$k_q$<br>$k_d$ | 0.33<br>14.82<br>3.87<br>0.02<br>2.27E-05 | (0.02, 3.16)<br>(NaN, NaN)<br>(NaN, NaN)<br>(0.008, 0.05)<br>(1e-14, 200) | 40.12 |
| 64 | LR size<br>$q=\phi^n/(k^n+\phi^n)$, $n=2$<br>$\tau=\tau_1+\tau_2\phi^n/(k^n+\phi^n)$, $n=2$ | $\alpha$<br>$\tau_1$<br>$\tau_2$<br>$k_q$<br>$k_d$ | 0.36<br>18.82<br>3.33<br>0.02<br>3.2 | (0.29, 0.450)<br>(15.81, 22.3)<br>(NaN, NaN)<br>(0.01, 0.02)<br>(NaN, NaN) | 40.12 |
| 65 | LR size<br>$q=\phi^n/(k^n+\phi^n)$, $n=2$<br>$\tau=\tau_1+\tau_2\phi^n/(k^n+\phi^n)$, $n=3$ | $\alpha$<br>$\tau_1$<br>$\tau_2$<br>$k_q$<br>$k_d$ | 0.05<br>14.9<br>5.07<br>0.06<br>0.06 | (NaN, NaN)<br>(11.63, 19.03)<br>(NaN, NaN)<br>(NaN, NaN)<br>(NaN, NaN) | 35.2 |
| 66 | LR size<br>$q=\phi^n/(k^n+\phi^n)$, $n=3$<br>$\tau=\tau_1+\tau_2\phi^n/(k^n+\phi^n)$, $n=1$ | $\alpha$<br>$\tau_1$<br>$\tau_2$<br>$k_q$<br>$k_d$ | 0.03<br>3.89<br>26.24<br>0.05<br>0.03 | (0.003, 0.32)<br>(3.63, 4.18)<br>(NaN, NaN)<br>(0.004, 0.69)<br>(0.003, 0.036) | 40.17 |
| 67 | LR size<br>$q=\phi^n/(k^n+\phi^n)$, $n=3$<br>$\tau=\tau_1+\tau_2\phi^n/(k^n+\phi^n)$, $n=2$ | $\alpha$<br>$\tau_1$<br>$\tau_2$ | 0.94<br>20.29<br>3.75 | (0.91, 0.98)<br>(19.27, 21.37)<br>(NaN, NaN) | 34.63 |
| | | $k_q$ | 0.03 | (0.03, 0.03) | |
| | | $k_d$ | 0.11 | (NaN, NaN) | |
| 68 | LR size<br>$q=\phi^n/(k^n+\phi^n)$, $n=3$<br>$\tau=\tau_1+\tau_2\phi^n/(k^n+\phi^n)$, $n=3$ | $\alpha$ | 0.39 | (0.28, 0.53) | 40.05 |
| | | $\tau_1$ | 17.4 | (16.71, 18.13) | |
| | | $\tau_2$ | 2.3 | (1.02, 5.15) | |
| | | $k_q$ | 0.04 | (0.039, 0.04) | |
| | | $k_d$ | 2.25E-05 | (1e-14, NaN) | |
| 69 | LR size<br>$q=\phi^n/(k^n+\phi^n)$, $n=1$<br>$\tau=\tau_1-\tau_2LR$ | $\alpha$ | 0.013 | (0.01, 0.016) | 39.05 |
| | | $\tau_1$ | 21.18 | (20.54, 21.83) | |
| | | $\tau_2$ | 1.56 | (1.42, 1.71) | |
| | | $k$ | 0.14 | (0.11, 0.16) | |
| 70 | LR size<br>$q=\phi^n/(k^n+\phi^n)$, $n=2$<br>$\tau=\tau_1-\tau_2LR$ | $\alpha$ | 0.44 | (NaN, NaN) | 37.05 |
| | | $\tau_1$ | 49.33 | (NaN, NaN) | |
| | | $\tau_2$ | 16.3 | (NaN, NaN) | |
| | | $k$ | 0.016 | (0.009, 0.034) | |
| 71 | LR size<br>$q=\phi^n/(k^n+\phi^n)$, $n=3$<br>$\tau=\tau_1-\tau_2LR$ | $\alpha$ | 0.9 | (0.4, 1.92) | 37.32 |
| | | $\tau_1$ | 33.5 | (NaN, NaN) | |
| | | $\tau_2$ | 7.63 | (NaN, NaN) | |
| | | $k$ | 0.029 | (0.018, 0.048) | |
| 72 | LR size<br>$q=\phi^n/(k^n+\phi^n)$, $n=1$<br>$\tau=\tau_1+\tau_2/LR$ | $\alpha$ | 0.012 | (0.005, 0.025) | 39.38 |
| | | $\tau_1$ | 11.88 | (NaN, NaN) | |
| | | $\tau_2$ | 11.88 | (NaN, NaN) | |
| | | $k$ | 0.09 | (0.01, 0.46) | |
| 73 | LR size<br>$q=\phi^n/(k^n+\phi^n)$, $n=2$<br>$\tau=\tau_1+\tau_2\phi/LR$ | $\alpha$ | 0.23 | (0.17, 0.29) | 37.39 |
| | | $\tau_1$ | 0.02 | (0.02, 0.02) | |
| | | $\tau_2$ | 34.98 | (30.68, 39.79) | |
| | | $k$ | 0.027 | (0.013, 0.052) | |
| 74 | LR size<br>$q=\phi^n/(k^n+\phi^n)$, $n=3$<br>$\tau=\tau_1+\tau_2/LR$ | $\alpha$ | 0.2 | (0.17, 0.23) | 37.22 |
| | | $\tau_1$ | 0.03 | (NaN, NaN) | |
| | | $\tau_2$ | 35.13 | (NaN, NaN) | |
| | | $k$ | 0.054 | (0.046, 0.063) | |
| 75 | LR size<br>$q=\phi^n/(k^n+\phi^n)$, $n=1$<br>$\tau=\tau_1/LR$ | $\alpha$ | 0.3 | (NaN, NaN) | 35.83 |
| | | $\tau_1$ | 35.97 | (NaN, NaN) | |
| | | $k$ | 0.003 | (0.022, 0.005) | |
| 76 | LR size<br>$q=\phi^n/(k^n+\phi^n)$, $n=2$<br>$\tau=\tau_1/LR$ | $\alpha$ | 0.23 | (0.05, 0.87) | 35.41 |
| | | $\tau_1$ | 34.94 | (16.81, 67.98) | |
| | | $k$ | 0.03 | (0.009, 0.07) | |
| 77 | LR size | $\alpha$ | 0.011 | (0.004, 0.03) | 40.91 |
| | $q = \phi^n / (k^n + \phi^n), n=1$<br>$\tau = \tau_1 + \tau_2(1 - LR^n / (k^n + LR^n)), n=1$ | $\tau_1$<br>$\tau_2$<br>$k_q$<br>$k_d$ | 12.96<br>4.51<br>0.2<br>0.009 | (12.41, 13.6)<br>(4.07, 5)<br>(0.11, 0.36)<br>(0.004, 0.02) | |
| 78 | LR size<br>$q = \phi^n / (k^n + \phi^n), n=1$<br>$\tau = \tau_1 + \tau_2(1 - LR^n / (k^n + LR^n)), n=2$ | $\alpha$<br>$\tau_1$<br>$\tau_2$<br>$k_q$<br>$k_d$ | 0.023<br>14.69<br>206.15<br>0.06<br>0.26 | (0.022, 0.024)<br>(14.61, 14.76)<br>(202.25, 209.96)<br>(0.058, 0.062)<br>(0.25, 0.262) | 41.33 |
| 79 | LR size<br>$q = \phi^n / (k^n + \phi^n), n=1$<br>$\tau = \tau_1 + \tau_2(1 - LR^n / (k^n + LR^n)), n=3$ | $\alpha$<br>$\tau_1$<br>$\tau_2$<br>$k_q$<br>$k_d$ | 0.01<br>14.02<br>3.04<br>0.16<br>0.007 | (NaN, NaN)<br>(13.71, 14.33)<br>(NaN, NaN)<br>(NaN, NaN)<br>(NaN, NaN) | 44.46 |
| 80 | LR size<br>$q = \phi^n / (k^n + \phi^n), n=2$<br>$\tau = \tau_1 + \tau_2(1 - LR^n / (k^n + LR^n)), n=1$ | $\alpha$<br>$\tau_1$<br>$\tau_2$<br>$k_q$<br>$k_d$ | 0.33<br>14.82<br>3.87<br>0.021<br>2.27E-05 | (0.026, 3.16)<br>(NaN, NaN)<br>(NaN, NaN)<br>(0.008, 0.54)<br>(1e-14, 200) | 40.12 |
| 81 | LR size<br>$q = \phi^n / (k^n + \phi^n), n=2$<br>$\tau = \tau_1 + \tau_2(1 - LR^n / (k^n + LR^n)), n=2$ | $\alpha$<br>$\tau_1$<br>$\tau_2$<br>$k_q$<br>$k_d$ | 0.42<br>6.04<br>299.96<br>0.017<br>0.18 | (0.38, 0.45)<br>(2.03, 17.49)<br>(1e-14, 300)<br>(0.013, 0.022)<br>(0.17, 0.19) | 39.21 |
| 82 | LR size<br>$q = \phi^n / (k^n + \phi^n), n=2$<br>$\tau = \tau_1 + \tau_2(1 - LR^n / (k^n + LR^n)), n=3$ | $\alpha$<br>$\tau_1$<br>$\tau_2$<br>$k_q$<br>$k_d$ | 0.05<br>14.9<br>5.07<br>0.066<br>0.06 | (NaN, NaN)<br>(11.63, 19.03)<br>(NaN, NaN)<br>(NaN, NaN)<br>(NaN, NaN) | 35.26 |
| 83 | LR size<br>$q = \phi^n / (k^n + \phi^n), n=3$<br>$\tau = \tau_1 + \tau_2(1 - LR^n / (k^n + LR^n)), n=1$ | $\alpha$<br>$\tau_1$<br>$\tau_2$<br>$k_q$<br>$k_d$ | 0.03<br>3.89<br>26.24<br>0.05<br>0.03 | (0.003, 0.004)<br>(3.63, 4.19)<br>(NaN, NaN)<br>(0.004, 0.69)<br>(0.002, 0.36) | 41.69 |
| 84 | LR size<br>$q = \phi^n / (k^n + \phi^n), n=3$<br>$\tau = \tau_1 + \tau_2(1 - LR^n / (k^n + LR^n)), n=2$ | $\alpha$<br>$\tau_1$<br>$\tau_2$ | 0.416<br>14.81<br>216.29 | (0.414, 0.418)<br>(13.82, 15.86)<br>(195.46, 234.37) | 39.42 |
| | | $k_q$ | 0.038 | (0.031, 0.046) | |
| | | $k_d$ | 0.19 | (0.18, 0.2) | |
| 85 | LR size<br>$q=\phi^n/(k^n+\phi^n)$, $n=3$<br>$\tau=\tau_1+\tau_2(1-LR^n/(k^n+LR^n))$, $n=3$ | $\alpha$ | 0.39 | (0.28, 0.54) | 41.56 |
| | | $\tau_1$ | 17.42 | (15.78, 18.16) | |
| | | $\tau_2$ | 2.3 | (1.02, 5.18) | |
| | | $k_q$ | 0.04 | (0.039, 0.043) | |
| | | $k_d$ | 2.23E-05 | (NaN, NaN) | |
| 86 | LR size<br>$\tau=\tau_1+\tau_2\phi/LR$ | $\alpha(1-q)$ | 0.005 | (0.003, 0.009) | 37.05 |
| | | $\tau_1$ | 6.65 | (6.01, 73.5) | |
| | | $\tau_2$ | 113.6 | (106.66, 120.72) | |
| 87 | LR size<br>$\tau=\tau_1+\tau_2\phi-\tau_3LR$ | $\alpha(1-q)$ | 0.005 | (0.003, 0.009) | 39.17 |
| | | $\tau_1$ | 7.88 | (7.2, 8.62) | |
| | | $\tau_2$ | 52.79 | (49.64, 56.1) | |
| | | $\tau_3$ | 1.01 | (0.82, 1.24) | |
| 88 | LR size<br>$q=\min(q_0\phi, 1)$<br>$\tau=\tau_1+\tau_2\phi/LR$ | $\alpha$ | 1.99 | (9.27e-10, 2) | 39.05 |
| | | $q_0$ | 11.94 | (11.24, 12.68) | |
| | | $\tau_1$ | 77.37 | (69.71, 85.68) | |
| | | $\tau_2$ | 0.011 | (0.005, 0.02) | |
| 89 | LR size<br>$q=\min(q_0\phi, 1)$<br>$\tau=\tau_1+\tau_2\phi-\tau_3LR$ | $\alpha$ | 0.01 | (0.005, 0.02) | 41.06 |
| | | $q_0$ | 1.92 | (1.5, 1.99) | |
| | | $\tau_1$ | 12.25 | (11.51, 13.05) | |
| | | $\tau_2$ | 47.7 | (43.92, 51.86) | |
| | | $\tau_3$ | 1.26 | (1.05, 1.51) | |
| 90 | LR size<br>$q=\phi^n/(k^n+\phi^n)$, $n=1$<br>$\tau=\tau_1+\tau_2\phi/LR$ | $\alpha$ | 0.08 | (0.054, 0.11) | 38.7 |
| | | $k$ | 0.014 | (0.01, 0.018) | |
| | | $\tau_1$ | 9.6 | (8.95, 10.3) | |
| | | $\tau_2$ | 142.4 | (131.89, 152.98) | |
| 91 | LR size<br>$q=\phi^n/(k^n+\phi^n)$, $n=2$<br>$\tau=\tau_1+\tau_2\phi/LR$ | $\alpha$ | 0.13 | (0.06, 0.27) | 38.26 |
| | | $k$ | 0.04 | (0.026, 0.06) | |
| | | $\tau_1$ | 9.78E+00 | (1.93, 44.6) | |
| | | $\tau_2$ | 207.76 | (NaN, NaN) | |
| 92 | LR size<br>$q=\phi^n/(k^n+\phi^n)$, $n=3$<br>$\tau=\tau_1+\tau_2\phi/LR$ | $\alpha$ | 0.19 | (0.16, 0.22) | 38.01 |
| | | $k$ | 0.06 | (0.05, 0.07) | |
| | | $\tau_1$ | 6.81 | (5.61, 8.26) | |
| | | $\tau_2$ | 290.7 | (NaN, NaN) | |
| 93 | LR size<br>$q=\phi^n/(k^n+\phi^n)$, $n=1$ | $\alpha$ | 0.1 | (0.08, 0.13) | 40.74 |
| | | $k$ | 0.009 | (0.007, 0.011) | |
| | $\tau=\tau_1+\tau_2\phi-\tau_3LR$ | $\tau_1$ | 14.91 | (14.23, 15.61) | |
| | | $\tau_2$ | 49.91 | (45.5, 54.65) | |
| | | $\tau_3$ | 1.46 | (1.27, 1.68) | |
| 94 | LR size<br>$q=\phi^n/(k^n+\phi^n)$, $n=2$<br>$\tau=\tau_1+\tau_2\phi-\tau_3LR$ | $\alpha$ | 1.26 | (1.15, 1.38) | 39 |
| | | $k$ | 0.009 | (0.008, 0.01) | |
| | | $\tau_1$ | 35.58 | (33.49, 37.78) | |
| | | $\tau_2$ | 4.53 | (4.79e-8, 299.9) | |
| | | $\tau_3$ | 10.01 | (9.13, 10.97) | |
| 95 | LR size<br>$q=\phi^n/(k^n+\phi^n)$, $n=3$<br>$\tau=\tau_1+\tau_2\phi-\tau_3LR$ | $\alpha$ | 0.71 | (0.51, 0.99) | 39.33 |
| | | $k$ | 0.032 | (0.02, 0.04) | |
| | | $\tau_1$ | 34.39 | (NaN, NaN) | |
| | | $\tau_2$ | 4.88 | (NaN, NaN) | |
| | | $\tau_3$ | 7.88 | (NaN, NaN) | |
| 96 | LR size<br>$\tau=\tau_1+\tau_2\phi-\tau_3LR+\tau_4\phi/LR$ | $\alpha(1-q)$ | 0.005 | (0.005, 0.006) | 40.99 |
| | | $\tau_1$ | 6.22 | (5.05, 7.64) | |
| | | $\tau_2$ | 5.8 | (2.24, 14.72) | |
| | | $\tau_3$ | 1.1 | (0.69, 1.74) | |
| | | $\tau_4$ | 132.64 | (119.19, 146.38) | |
| 97 | LR size<br>$\tau=\tau_1+\tau_2\phi+\tau_3LR+\tau_4\phi/LR$ | $\alpha(1-q)$ | 0.005 | (0.002, 0.011) | 41 |
| | | $\tau_1$ | 7.57 | (6.91, 8.31) | |
| | | $\tau_2$ | 5.33 | (3.72, 7.6) | |
| | | $\tau_3$ | 20.45 | (18.8, 22.23) | |
| | | $\tau_4$ | 0.41 | (0.0019, 67.7) | |
| 98 | LR size<br>$q=\min(q_0\phi, 1)$<br>$\tau=\tau_1+\tau_2\phi-\tau_3LR+\tau_4\phi/LR$ | $\alpha$ | 0.011 | (0.006, 0.019) | 43.03 |
| | | $q_0$ | 1.97 | (0.021, 1.99) | |
| | | $\tau_1$ | 9.33 | (8.58, 10.13) | |
| | | $\tau_2$ | 58.63 | (54.47, 63.03) | |
| | | $\tau_3$ | 1.33 | (1.11, 1.6) | |
| | | $\tau_4$ | 17.72 | (12.11, 25.68) | |
| 99 | LR size<br>$q=\phi^n/(k^n+\phi^n)$, $n=1$<br>$\tau=\tau_1+\tau_2\phi-\tau_3LR+\tau_4\phi/LR$ | $\alpha$ | 0.016 | (0.01, 0.02) | 42.89 |
| | | $k$ | 0.1 | (0.086, 0.13) | |
| | | $\tau_1$ | 7.19 | (6.52, 7.92) | |
| | | $\tau_2$ | 12.33 | (9.7, 15.63) | |
| | | $\tau_3$ | 1.27 | (1.05, 1.54) | |
| | | $\tau_4$ | 153.46 | (144.65, 16.25) | |
| 100 | LR size<br>$q=\phi^n/(k^n+\phi^n)$, $n=2$ | $\alpha$ | 0.79 | (0.19, 2.72) | 40.01 |
| | | $k$ | 0.01 | (0.0019, 0.08) | |
| | $\tau=\tau_1+\tau_2\phi-\tau_3LR+\tau_4\phi/LR$ | $\tau_1$ | 40.02 | (NaN, NaN) | |
| | | $\tau_2$ | 0.0001 | (0.001, 0.0001) | |
| | | $\tau_3$ | 12.26 | (NaN, NaN) | |
| | | $\tau_4$ | 40.2 | (7.39e-7, 299.9) | |
| 101 | LR size<br>$q=\phi^n/(k^n+\phi^n)$, $n=3$<br>$\tau=\tau_1+\tau_2\phi-\tau_3LR+\tau_4\phi/LR$ | $\alpha$ | 0.02 | (0.005, 0.07) | 42.79 |
| | | $k$ | 0.15 | (NaN, NaN) | |
| | | $\tau_1$ | 15.2 | (12.21, 18.8) | |
| | | $\tau_2$ | 33.1 | (14.13, 71.16) | |
| | | $\tau_3$ | 1.11 | (0.32, 3.82) | |
| | | $\tau_4$ | 17.65 | (0.46, 214.69) | |
| 102 | LR size<br>$q=\min(q_0\phi, 1)$<br>$\tau=\tau_1+\tau_2\phi+\tau_3LR+\tau_4\phi/LR$ | $\alpha$ | 0.011 | (0.009, 0.01) | 43.07 |
| | | $q_0$ | 1.99 | (1e-14, NaN) | |
| | | $\tau_1$ | 7.02 | (6.44, 7.66) | |
| | | $\tau_2$ | 55.11 | (51.42, 59.01) | |
| | | $\tau_3$ | 2.24 | (0.5, 9.5) | |
| | | $\tau_4$ | 0.76 | (8.46e-13, 300) | |
| 103 | LR size<br>$q=\phi^n/(k^n+\phi^n)$, $n=1$<br>$\tau=\tau_1+\tau_2\phi+\tau_3LR+\tau_4\phi/LR$ | $\alpha$ | NaN | (NaN, NaN) | NaN |
| | | $k$ | NaN | (NaN, NaN) | |
| | | $\tau_1$ | NaN | (NaN, NaN) | |
| | | $\tau_2$ | NaN | (NaN, NaN) | |
| | | $\tau_3$ | NaN | (NaN, NaN) | |
| | | $\tau_4$ | NaN | (NaN, NaN) | |
| 104 | LR size<br>$q=\phi^n/(k^n+\phi^n)$, $n=2$<br>$\tau=\tau_1+\tau_2\phi+\tau_3LR+\tau_4\phi/LR$ | $\alpha$ | NaN | (NaN, NaN) | NaN |
| | | $k$ | NaN | (NaN, NaN) | |
| | | $\tau_1$ | NaN | (NaN, NaN) | |
| | | $\tau_2$ | NaN | (NaN, NaN) | |
| | | $\tau_3$ | NaN | (NaN, NaN) | |
| | | $\tau_4$ | NaN | (NaN, NaN) | |
| 105 | LR size<br>$q=\phi^n/(k^n+\phi^n)$, $n=3$<br>$\tau=\tau_1+\tau_2\phi+\tau_3LR+\tau_4\phi/LR$ | $\alpha$ | NaN | (NaN, NaN) | NaN |
| | | $k$ | NaN | (NaN, NaN) | |
| | | $\tau_1$ | NaN | (NaN, NaN) | |
| | | $\tau_2$ | NaN | (NaN, NaN) | |
| | | $\tau_3$ | NaN | (NaN, NaN) | |
| | | $\tau_4$ | NaN | (NaN, NaN) | |
| 106 | LR size<br>$\tau=\tau_1+\tau_2\phi-\tau_3LR+\tau_4\phi*LR$ | $\alpha(1-q)$ | 0.005 | (0.003, 0.009) | 41.2 |
| | | $\tau_1$ | 7.05 | (6.37, 7.8) | |
| | | $\tau_2$ | 14.26 | (11.95, 16.99) | |
| | | $\tau_3$ | 0.63 | (0.46, 0.85) | |
| | | $\tau_4$ | 15.11 | (13.84, 16.39) | |
| 107 | LR size<br>$q = \min(q_0\phi, 1)$<br>$\tau = \tau_1 + \tau_2\phi - \tau_3LR + \tau_4\phi * LR$ | $\alpha$ | 0.011 | (0.007, 0.017) | 43.09 |
| | | $q_0$ | 1.95 | (1.8, 1.99) | |
| | | $\tau_1$ | 7.96 | (7.25, 8.73) | |
| | | $\tau_2$ | 57.95 | (53.86, 62.26) | |
| | | $\tau_3$ | 1 | (0.81, 1.24) | |
| | | $\tau_4$ | 4.32 | (2.72, 5.91) | |
| 108 | LR size<br>$q = \phi^n / (k^n + \phi^n), n=1$<br>$\tau = \tau_1 + \tau_2\phi - \tau_3LR + \tau_4\phi * LR$ | $\alpha$ | 0.012 | (0.012, 0.013) | 43.14 |
| | | $k$ | 0.14 | (0.13, 0.15) | |
| | | $\tau_1$ | 11.8 | (11.22, 12.4) | |
| | | $\tau_2$ | 19 | (16.73, 21.55) | |
| | | $\tau_3$ | 0.7 | (0.58, 0.85) | |
| | | $\tau_4$ | 8.83 | (7.74, 9.91) | |
| 109 | LR size<br>$q = \phi^n / (k^n + \phi^n), n=2$<br>$\tau = \tau_1 + \tau_2\phi - \tau_3LR + \tau_4\phi * LR$ | $\alpha$ | 0.02 | (0.003, 0.3) | 42.61 |
| | | $k$ | 0.07 | (0.02, 0.25) | |
| | | $\tau_1$ | 16.71 | (14.66, 19.03) | |
| | | $\tau_2$ | 21.03 | (7.7, 53.25) | |
| | | $\tau_3$ | 0.85 | (0.3, 2.4) | |
| | | $\tau_4$ | 2.06 | (-6.34, 10.46) | |
| 110 | LR size<br>$q = \phi^n / (k^n + \phi^n), n=3$<br>$\tau = \tau_1 + \tau_2\phi - \tau_3LR + \tau_4\phi * LR$ | $\alpha$ | 0.11 | (0.069, 0.2) | 42 |
| | | $k$ | 0.07 | (0.06, 0.08) | |
| | | $\tau_1$ | 24.09 | (21.32, 27.19) | |
| | | $\tau_2$ | 108.06 | (98.11, 118.41) | |
| | | $\tau_3$ | 6.54 | (6.17, 6.92) | |
| | | $\tau_4$ | -0.26 | (-9.17, 8.64) | |

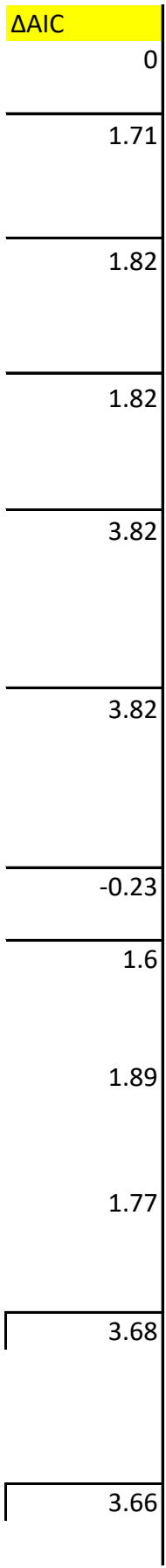

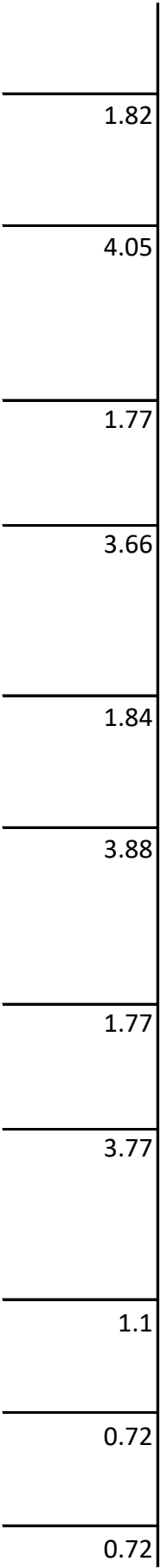

3.27

1.54

0.69

0.69

3.22

3.82

3.82

3.82

5.82

5.98

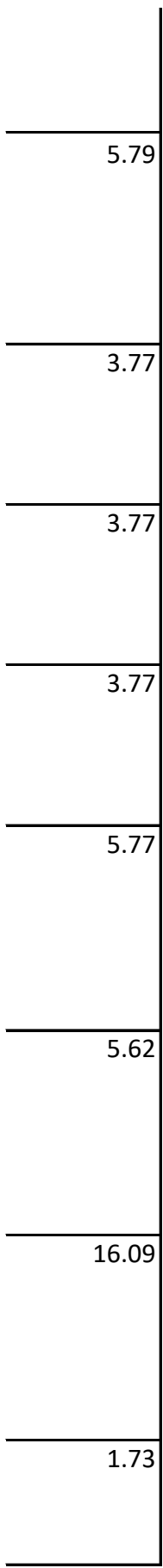

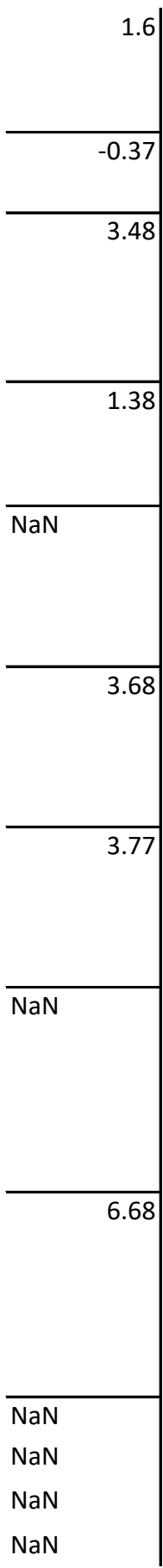

NaN

3.73

3.17

3.2

2.78

3.12

4.16

2.68

2.89

2.94

5.68

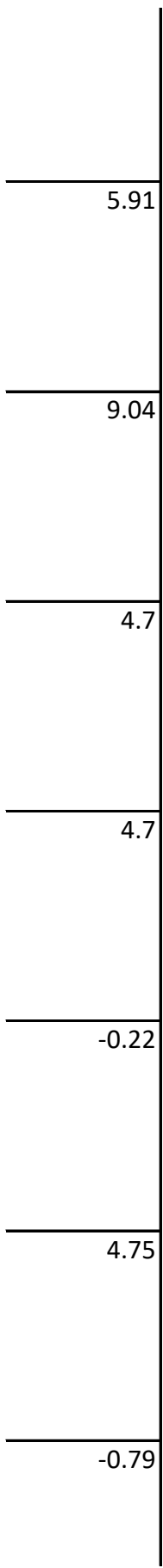

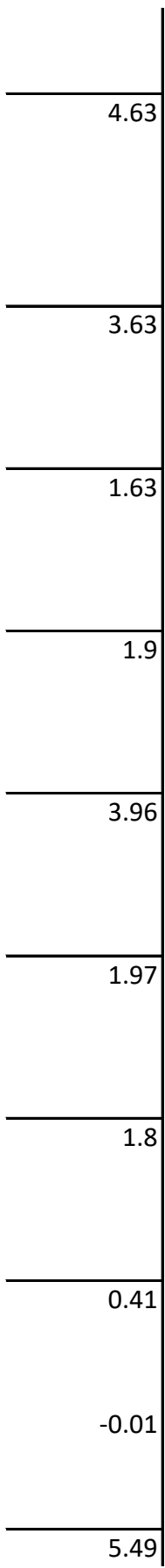

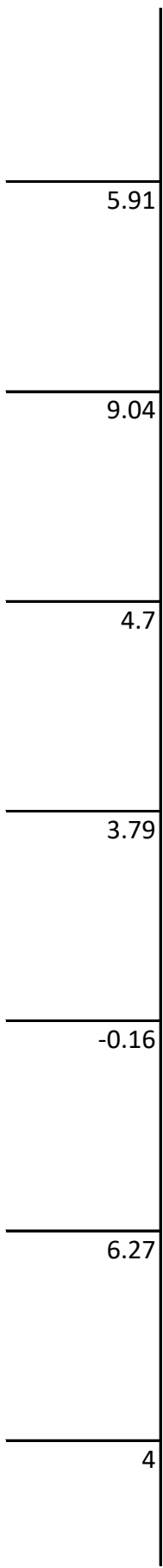

6.14

1.63

3.75

3.63

5.64

3.28

2.84

2.59

5.32

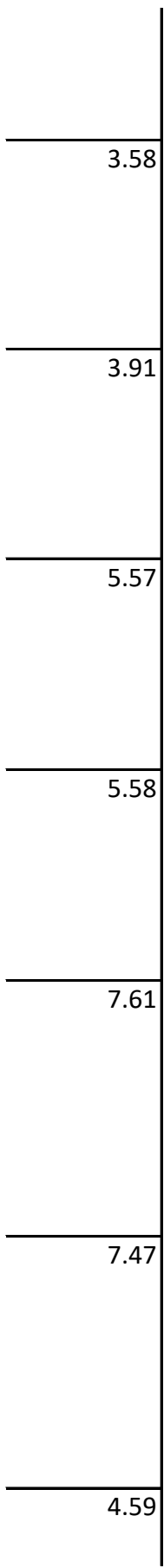

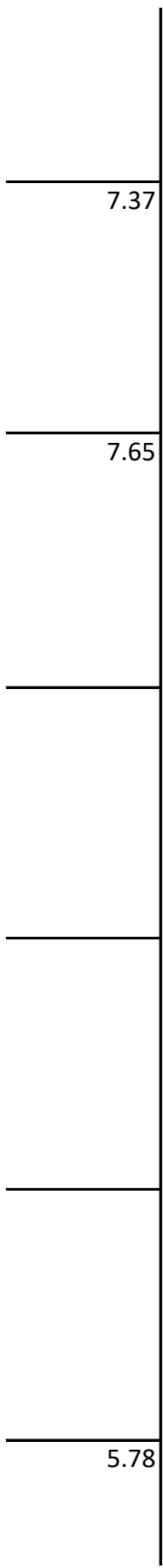

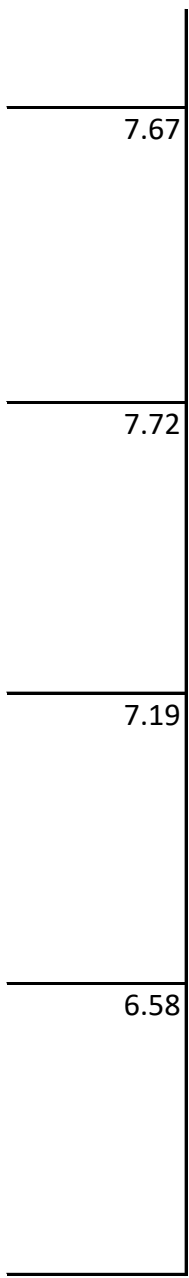

| Model | Model Description | Parameters | Parameter | 95% CI | AIC |
| --- | --- | --- | --- | --- | --- |
| 1 | Null | $\alpha(1-q)$<br>$\tau$ | 0.032<br>4.88 | (0.02, 0.05)<br>(2.52, 8.73) | 157.57 |
| 2 | $q=\min(q_0\phi, 1)$ | $\alpha$<br>$q_0$<br>$\tau$ | 0.03<br>2.25E-07<br>4.88 | (0.02, 0.05)<br>(2.25e-7, 2.25e-7)<br>(2.54, 8.84) | 159.58 |
| 3 | $\tau=\tau_1+(\tau_2-\tau_1)\phi$ | $\alpha(1-q)$<br>$\tau_1$<br>$\tau_2$ | 0.03<br>5.76<br>3.72 | (0.02, 0.05)<br>(2.12, 11.6)<br>(0.41, 20.05) | 159.43 |
| 4 | $\tau=\tau_1+(\tau_2-\tau_1)\phi^2$ | $\alpha(1-q)$<br>$\tau_1$<br>$\tau_2$ | 0.03<br>5.78<br>3.5 | (0.02, 0.05)<br>(3.28, 9.14)<br>(0.25, 23.52) | 159.58 |
| 5 | $q=\min(q_0\phi, 1)$<br>$\tau=\tau_1+(\tau_2-\tau_1)\phi$ | $\alpha$<br>$q_0$<br>$\tau_1$<br>$\tau_2$ | 0.03<br>2.25E-07<br>5.76<br>3.72 | (0.02, 0.05)<br>(2.25e-7, 2.25e-7)<br>(2.12, 11.6)<br>(0.41, 20.05) | 161.43 |
| 6 | $q=\min(q_0\phi, 1)$<br>$\tau=\tau_1+(\tau_2-\tau_1)\phi^2$ | $\alpha$<br>$q_0$<br>$\tau_1$<br>$\tau_2$ | 0.03<br>2.25E-07<br>5.28<br>3.93 | (0.02, 0.05)<br>(2.25e-7, 2.25e-7)<br>(2.26, 10.04)<br>(0.39, 21.71) | 161.51 |
| 7 | LR size | $\alpha(1-q)$<br>$\tau$ | 0.012<br>5.08 | (0.008, 0.02)<br>(2.94, 8.45) | 152.43 |
| 8 | LR size<br>$q=\min(q_0\phi, 1)$ | $\alpha$<br>$q_0$<br>$\tau$ | 0.012<br>2.34E-07<br>5.11 | (0.008, 0.02)<br>(2.34e-7, 2.34e-7)<br>(2.94, 8.52) | 154.46 |
| 9 | LR size<br>$\tau=\tau_1+(\tau_2-\tau_1)\phi$ | $\alpha(1-q)$<br>$\tau_1$<br>$\tau_2$ | 0.012<br>5.34<br>4.82 | (0.008, 0.02)<br>(1.95, 12.66)<br>(1.37, 13.8) | 154.42 |
| 10 | LR size<br>$\tau=\tau_1+(\tau_2-\tau_1)\phi^2$ | $\alpha(1-q)$<br>$\tau_1$<br>$\tau_2$ | 0.012<br>5.02<br>5.21 | (0.008, 0.02)<br>(2.01, 11.2)<br>(1.41, 15.1) | 154.43 |
| 11 | LR size<br>$q=\min(q_0\phi, 1)$<br>$\tau=\tau_1+(\tau_2-\tau_1)\phi$ | $\alpha$<br>$q_0$<br>$\tau_1$<br>$\tau_2$ | 0.012<br>2.84E-06<br>5.34<br>4.82 | (0.008, 0.02)<br>(2.84E-06, 2.84E-06)<br>(2.03, 13.59)<br>(1.45, 14.8) | 156.42 |
| 12 | LR size<br>$q=\min(q_0\phi, 1)$ | $\alpha$<br>$q_0$ | 0.012<br>2.25E-07 | (0.008, 0.02)<br>(2.25e-7, 2.25e-7) | 156.44 |
| | $\tau=\tau_1+(\tau_2-\tau_1)\phi^2$ | $\tau_1$ | 5.02 | (2.06, 11.7) | |
| | | $\tau_2$ | 5.21 | (1.49, 16.46) | |
| 13 | $\tau=\tau_1+\tau_2\phi$ | $\alpha(1-q)$ | 0.03 | (0.02, 0.05) | 159.73 |
| | | $\tau_1$ | 4.15 | (0.18, 17.62) | |
| | | $\tau_2$ | 0.44 | (4.37e-13, 40) | |
| 14 | $q=\min(q_0\phi, 1)$ | $\alpha$ | 0.03 | (0.02, 0.05) | 161.58 |
| | $\tau=\tau_1+\tau_2\phi$ | $q_0$ | 2.25E-07 | (2.25e-7, 2.25e-7) | |
| | | $\tau_1$ | 4.88 | (NaN, NaN) | |
| | | $\tau_2$ | 4.50E-06 | (NaN, NaN) | |
| 15 | LR size | $\alpha(1-q)$ | 0.02 | (0.008, 0.02) | 154.44 |
| | $\tau=\tau_1+\tau_2\phi$ | $\tau_1$ | 5.09 | (2.92, 8.5) | |
| | | $\tau_2$ | 4.50E-06 | (4.5e-6, 4.5e-6) | |
| 16 | LR size | $\alpha$ | 0.01 | (0.008, 0.02) | 156.43 |
| | $q=\min(q_0\phi, 1)$ | $q_0$ | 2.25E-07 | (2.25e-7, 2.25e-7) | |
| | $\tau=\tau_1+\tau_2\phi$ | $\tau_1$ | 5.09 | (2.95, 8.6) | |
| | | $\tau_2$ | 1.12E-05 | (1.12e-5, 1.12e-5) | |
| 17 | $\tau=\tau_1+(\tau_2+\tau_1)\phi$ | $\alpha(1-q)$ | 0.03 | (0.02, 0.05) | 160.28 |
| | | $\tau_1$ | 2.8 | (1.38, 5.42) | |
| | | $\tau_2$ | 4.50E-06 | (4.50E-06, 4.50E-06) | |
| 18 | $q=\min(q_0\phi, 1)$ | $\alpha$ | 0.03 | (0.02, 0.05) | 161.43 |
| | $\tau=\tau_1+(\tau_2+\tau_1)\phi$ | $q_0$ | 2.25E-07 | (2.25e-7, 2.25e-7) | |
| | | $\tau_1$ | 5.76 | (2.12, 11.6) | |
| | | $\tau_2$ | 3.72 | (0.41, 20.05) | |
| 19 | LR size | $\alpha(1-q)$ | 0.01 | (0.008, 0.02) | 155.43 |
| | $\tau=\tau_1+(\tau_2+\tau_1)\phi$ | $\tau_1$ | 1.96 | (NaN, NaN) | |
| | | $\tau_2$ | 2.3 | (NaN, NaN) | |
| 20 | LR size | $\alpha$ | 0.01 | (0.008, 0.02) | 156.97 |
| | $q=\min(q_0\phi, 1)$ | $q_0$ | 2.25E-07 | (2.25e-7, 2.25e-7) | |
| | $\tau=\tau_1+(\tau_2+\tau_1)\phi$ | $\tau_1$ | 3.01 | (1.72, 5.22) | |
| | | $\tau_2$ | 1.12E-05 | (1.12e-5, 1.12e-5) | |
| 21 | $q=\phi^n/(k^n+\phi^n), n=1$ | $\alpha(1-q)$ | 0.03 | (0.02, 0.05) | 159.58 |
| | | $k$ | 499.99 | (1.11e-15, 500) | |
| | | $\tau$ | 4.87 | (2.54, 8.84) | |
| 22 | $q=\phi^n/(k^n+\phi^n), n=2$ | $\alpha(1-q)$ | 0.03 | (0.02, 0.05) | 159.57 |
| | | $k$ | 490.77 | (490.77, 490.77) | |
| | | $\tau$ | 4.87 | (2.54, 8.84) | |
| 23 | $q=\phi^n/(k^n+\phi^n), n=3$ | $\alpha(1-q)$ | 0.03 | (0.02, 0.05) | 159.57 |
|  |  | k | 231.77 | (231.77, 231.77) |  |
| | | $\tau$ | 4.88 | (2.54, 8.84) | |
| 24 | $q=q_0+q_1\phi$ | $\alpha$ | 0.03 | (0.02, 0.05) | 161.58 |
| | | $q_0$ | 3.41 | (NaN, NaN) | |
| | | $q_1$ | 1.12E-07 | (1e-14, NaN) | |
| | | $\tau$ | 4.88 | (2.54, 8.84) | |
| 25 | LR size<br>$q=\phi^n/(k^n+\phi^n)$, n=1 | $\alpha$ | 0.014 | (0.09, 0.02) | 154.99 |
|  |  | k | 1.99 | (1.16e-9, 2) |  |
| | | $\tau$ | 5.07 | (2.88, 8.55) | |
| 26 | LR size<br>$q=\phi^n/(k^n+\phi^n)$, n=2 | $\alpha$ | 0.014 | (0.009, 0.02) | 154.56 |
|  |  | k | 1.99 | (1.99, 1.99) |  |
| | | $\tau$ | 5.08 | (2.92, 8.47) | |
| 27 | LR size<br>$q=\phi^n/(k^n+\phi^n)$, n=3 | $\alpha$ | 0.0142 | (0.009, 0.02) | 154.38 |
|  |  | k | 1.99 | (1.99, 1.99) |  |
| | | $\tau$ | 5.09 | (2.89, 8.56) | |
| 28 | LR size<br>$q=q_0+q_1\phi$ | $\alpha$ | 0.02 | (0.0003, 0.98) | 156.44 |
| | | $q_0$ | 0.35 | (5.75e-5, 1.99) | |
| | | $q_1$ | 1.25E-05 | (1.25e-5, 1.25e-5) | |
| | | $\tau$ | 5.09 | (2.92, 8.5) | |
| 29 | $\tau=\tau_1+\tau_2\phi^n/(k^n+\phi^n)$, n=1 | $\alpha(1-q)$ | 0.03 | (0.02, 0.04) | 161.58 |
| | | $\tau_1$ | 4.86 | (2.45, 8.5) | |
| | | $\tau_2$ | 2.63 | (NaN, NaN) | |
|  |  | k | 68.68 | (NaN, NaN) |  |
| 30 | $q=\phi^n/(k^n+\phi^n)$, n=2 | $\alpha(1-q)$ | 0.03 | (0.02, 0.04) | 161.58 |
| | | $\tau_1$ | 4.46 | (NaN, NaN) | |
| | | $\tau_2$ | 0.41 | (NaN, NaN) | |
|  |  | k | 1.12E-05 | (1e-14, NaN) |  |
| 31 | $\tau=\tau_1+\tau_2\phi^n/(k^n+\phi^n)$, n=3 | $\alpha(1-q)$ | 0.03 | (0.02, 0.04) | 161.58 |
| | | $\tau_1$ | 4.15 | (NaN, NaN) | |
| | | $\tau_2$ | 0.72 | (NaN, NaN) | |
|  |  | k | 2.01E-05 | (1e-14, 100) |  |
| 32 | $q=\min(q_0\phi, 1)$<br>$\tau=\tau_1+\tau_2\phi^n/(k^n+\phi^n)$, n=1 | $\alpha$ | 0.03 | (0.02, 0.04) | 163.58 |
| | | $q_0$ | 2.25E-07 | (2.25e-7, 2.25e-7) | |
| | | $\tau_1$ | 0.93 | (NaN, NaN) | |
| | | $\tau_2$ | 3.94 | (NaN, NaN) | |
|  |  | k | 1.12E-05 | (3.59e-9, 0.03) |  |
| 33 | $q=\min(q_0\phi, 1)$<br>$\tau=\tau_1+\tau_2\phi^n/(k^n+\phi^n)$, n=2 | $\alpha$ | 0.03 | (0.02, 0.04) | 163.58 |
| | | $q_0$ | 2.25E-07 | (2.25e-7, 2.25e-7) | |
| | | $\tau_1$ | 1.49 | (NaN, NaN) | |
| | | $\tau_2$ | 3.38 | (NaN, NaN) | |
|  |  | k | 1.12E-05 | (1e-14, 100) |  |
| 33 | $q=\min(q_0\phi, 1)$<br>$\tau=\tau_1+\tau_2\phi^n/(k^n+\phi^n), n=3$ | $\alpha$ | 0.1 | (0.08, 0.13) | 201.26 |
| | | $q_0$ | 0.5 | (0.16, 1.09) | |
| | | $\tau_1$ | 3.41 | (2.13, 5.44) | |
| | | $\tau_2$ | 2 | (NaN, NaN) | |
|  |  | k | 0.1 | (0.012, 0.82) |  |
| 34 | LR size<br>$\tau=\tau_1+\tau_2\phi^n/(k^n+\phi^n), n=1$ | $\alpha(1-q)$ | 0.01 | (0.009, 0.02) | 156.44 |
| | | $\tau_1$ | 2.64 | (0.05, 98.42) | |
| | | $\tau_2$ | 2.48 | (0.03, 109.78) | |
|  |  | k | 0.0001 | (4.38e-11, 86.73) |  |
| 35 | LR size<br>$\tau=\tau_1+\tau_2\phi^n/(k^n+\phi^n), n=2$ | $\alpha(1-q)$ | 0.01 | (0.009, 0.02) | 156.44 |
| | | $\tau_1$ | 2.41 | (NaN, NaN) | |
| | | $\tau_2$ | 2.67 | (NaN, NaN) | |
|  |  | k | 2.73E-05 | (1e-14, 100) |  |
| 36 | LR size<br>$\tau=\tau_1+\tau_2\phi^n/(k^n+\phi^n), n=3$ | $\alpha(1-q)$ | 0.01 | (0.008, 0.02) | 156.44 |
| | | $\tau_1$ | 2.54 | (NaN, NaN) | |
| | | $\tau_2$ | 2.54 | (NaN, NaN) | |
|  |  | k | 0.0008 | (1e-14, 100) |  |
| 37 | LR size<br>$q=\min(q_0\phi, 1)$<br>$\tau=\tau_1+\tau_2\phi^n/(k^n+\phi^n), n=1$ | $\alpha$ | NaN | (NaN, NaN) | NaN |
| | | $q_0$ | NaN | (NaN, NaN) | |
| | | $\tau_1$ | NaN | (NaN, NaN) | |
| | | $\tau_2$ | NaN | (NaN, NaN) | |
|  |  | k | NaN | (NaN, NaN) |  |
| 38 | LR size<br>$q=\min(q_0\phi, 1)$<br>$\tau=\tau_1+\tau_2\phi^n/(k^n+\phi^n), n=2$ | $\alpha$ | 0.32 | (0.15, 0.68) | 152.59 |
| | | $q_0$ | 1.86 | (1.32, 2.6) | |
| | | $\tau_1$ | 299.99 | (1e-14, 300) | |
| | | $\tau_2$ | 0.4 | (0.32, 0.49) | |
|  |  | k | 0.01 | (0.014, 0.02) |  |
| 39 | LR size<br>$q=\min(q_0\phi, 1)$<br>$\tau=\tau_1+\tau_2\phi^n/(k^n+\phi^n), n=3$ | $\alpha$ | NaN | (NaN, NaN) | NaN |
| | | $q_0$ | NaN | (NaN, NaN) | |
| | | $\tau_1$ | NaN | (NaN, NaN) | |
| | | $\tau_2$ | NaN | (NaN, NaN) | |
|  |  | k | NaN | (NaN, NaN) |  |
| 40 | LR size<br>$\tau=\tau_1-\tau_2LR$ | $\alpha(1-q)$ | 0.013 | (0.009, 0.02) | 148.56 |
| | | $\tau_1$ | 28.59 | (28, 24, 28.95) | |
| | | $\tau_2$ | 6.73 | (6.62, 6.85) | |
| 41 | LR size<br>$\tau=\tau_1+\tau_2/LR$ | $\alpha(1-q)$<br>$\tau_1$<br>$\tau_2$ | 0.013<br>3.37E-05<br>19.78 | (0.009, 0.02)<br>(3.37E-05, 3.37E-05)<br>(13.79, 28.1) | 151.8 |
| 42 | LR size<br>$\tau=\tau_1/LR$ | $\alpha(1-q)$<br>$\tau_1$ | 0.012<br>16.92 | (NaN, NaN)<br>(9.6, 29.26) | 150.22 |
| 43 | LR size<br>$q=\min(q_0\phi, 1)$<br>$\tau=\tau_1-\tau_2LR$ | $\alpha$<br>$q_0$<br>$\tau_1$<br>$\tau_2$ | 0.015<br>0.22<br>21.52<br>4.67 | (0.007, 0.03)<br>(0.002, 1.86)<br>(NaN, NaN)<br>(NaN, NaN) | 151.81 |
| 44 | LR size<br>$q=\min(q_0\phi, 1)$<br>$\tau=\tau_1/LR$ | $\alpha$<br>$q_0$<br>$\tau_1$ | 0.01<br>0.1<br>19.81 | (0.006, 0.02)<br>(3.97e-6, 1.99)<br>(19.85, 28.09) | 151.77 |
| 45 | LR size<br>$\tau=\tau_1+\tau_2(1-LR^n/(k^n+LR^n)), n=1$ | $\alpha(1-q)$<br>$\tau_1$<br>$\tau_2$<br>$k$ | NaN<br>NaN<br>NaN<br>NaN | (NaN, NaN)<br>(NaN, NaN)<br>(NaN, NaN)<br>(NaN, NaN) | NaN |
| 46 | LR size<br>$\tau=\tau_1+\tau_2(1-LR^n/(k^n+LR^n)), n=2$ | $\alpha(1-q)$<br>$\tau_1$<br>$\tau_2$<br>$k$ | NaN<br>NaN<br>NaN<br>NaN | (NaN, NaN)<br>(NaN, NaN)<br>(NaN, NaN)<br>(NaN, NaN) | NaN |
| 47 | LR size<br>$\tau=\tau_1+\tau_2(1-LR^n/(k^n+LR^n)), n=3$ | $\alpha(1-q)$<br>$\tau_1$<br>$\tau_2$<br>$k$ | NaN<br>NaN<br>NaN<br>NaN | (NaN, NaN)<br>(NaN, NaN)<br>(NaN, NaN)<br>(NaN, NaN) | NaN |
| 48 | LR size<br>$q=\min(q_0\phi, 1)$<br>$\tau=\tau_1+\tau_2(1-LR^n/(k^n+LR^n)), n=1$ | $\alpha$<br>$q_0$<br>$\tau_1$<br>$\tau_2$<br>$k$ | 0.009<br>1.77<br>10.35<br>7.05<br>0.01 | (NaN, NaN)<br>(0.06, 1.99)<br>(NaN, NaN)<br>(NaN, NaN)<br>(0.001, 0.02) | 141.19 |
| 49 | LR size<br>$q=\min(q_0\phi, 1)$<br>$\tau=\tau_1+\tau_2(1-LR^n/(k^n+LR^n)), n=2$ | $\alpha$<br>$q_0$<br>$\tau_1$<br>$\tau_2$<br>$k$ | 0.012<br>0.1<br>2.68<br>299.99<br>0.34 | (0.006, 0.022)<br>(0.0012, 7.47)<br>(0.59, 11.87)<br>(1e-14, 200)<br>(0.19, 0.6) | 155.18 |
| 50 | LR size<br>$q=\min(q_0\phi, 1)$<br>$\tau=\tau_1+\tau_2(1-LR^n/(k^n+LR^n)), n=3$ | $\alpha$<br>$q_0$<br>$\tau_1$<br>$\tau_2$ | NaN<br>NaN<br>NaN<br>NaN | (NaN, NaN)<br>(NaN, NaN)<br>(NaN, NaN)<br>(NaN, NaN) | NaN |

|  |  | k | NaN | (NaN, NaN) |  |
| --- | --- | --- | --- | --- | --- |
| 51 | $q=\phi^n/(k^n+\phi^n)$, n=1<br>$\tau=\tau_1+\tau_2\phi^n/(k^n+\phi^n)$, n=1 | $\alpha$<br>$\tau_1$<br>$\tau_2$<br>k | 0.03<br>4.86<br>4.24<br>199.99 | (0.02, 0.05)<br>(2.49, 9.4)<br>(1e-14, 200)<br>(NaN, NaN) | 161.59 |
| 52 | $q=\phi^n/(k^n+\phi^n)$, n=1<br>$\tau=\tau_1+\tau_2\phi^n/(k^n+\phi^n)$, n=2 | $\alpha$<br>$\tau_1$<br>$\tau_2$<br>k | 0.03<br>3.86<br>199.99<br>199.99 | (0.019, 0.05)<br>(0.77, 18.1)<br>(NaN, NaN)<br>(NaN, NaN) | 161.9 |
| 53 | $q=\phi^n/(k^n+\phi^n)$, n=1<br>$\tau=\tau_1+\tau_2\phi^n/(k^n+\phi^n)$, n=3 | $\alpha$<br>$\tau_1$<br>$\tau_2$<br>k | 0.032<br>4.87<br>2.36<br>199.99 | (0.02, 0.05)<br>(2.59, 9.08)<br>(1.84, 3.02)<br>(165.32, 200) | 161.59 |
| 54 | $q=\phi^n/(k^n+\phi^n)$, n=2<br>$\tau=\tau_1+\tau_2\phi^n/(k^n+\phi^n)$, n=1 | $\alpha$<br>$\tau_1$<br>$\tau_2$<br>k | 0.03<br>4.88<br>2.25E-05<br>199.99 | (0.02, 0.05)<br>(2.59, 9.08)<br>(NaN, NaN)<br>(NaN, NaN) | 161.57 |
| 55 | $q=\phi^n/(k^n+\phi^n)$, n=2<br>$\tau=\tau_1+\tau_2\phi^n/(k^n+\phi^n)$, n=2 | $\alpha$<br>$\tau_1$<br>$\tau_2$<br>k | 0.03<br>4.88<br>5.41<br>199.99 | (0.02, 0.05)<br>(2.59, 9.08)<br>(1e-14, 200)<br>(NaN, NaN) | 161.57 |
| 56 | $q=\phi^n/(k^n+\phi^n)$, n=2<br>$\tau=\tau_1+\tau_2\phi^n/(k^n+\phi^n)$, n=3 | $\alpha$<br>$\tau_1$<br>$\tau_2$<br>k | 0.03<br>4.88<br>5.41<br>199.99 | (0.02, 0.05)<br>(2.59, 9.08)<br>(NaN, NaN)<br>(NaN, NaN) | 161.57 |
| 57 | $q=\phi^n/(k^n+\phi^n)$, n=3<br>$\tau=\tau_1+\tau_2\phi^n/(k^n+\phi^n)$, n=1 | $\alpha$<br>$\tau_1$<br>$\tau_2$<br>k | 0.03<br>4.88<br>0.0001<br>199.99 | (0.02, 0.05)<br>(2.59, 9.08)<br>(7.15e-5, 0.0003)<br>(NaN, NaN) | 161.57 |
| 58 | $q=\phi^n/(k^n+\phi^n)$, n=3<br>$\tau=\tau_1+\tau_2\phi^n/(k^n+\phi^n)$, n=3 | $\alpha$<br>$\tau_1$<br>$\tau_2$<br>k | 0.03<br>4.88<br>2.65E-05<br>195.4 | (0.02, 0.05)<br>(2.59, 9.08)<br>(NaN, NaN)<br>(195.21, 195.59) | 161.57 |
| 59 | $q=\phi^n/(k^n+\phi^n)$, n=3<br>$\tau=\tau_1+\tau_2\phi^n/(k^n+\phi^n)$, n=3 | $\alpha$<br>$\tau_1$<br>$\tau_2$<br>k | 0.03<br>4.88<br>2.9<br>193.45 | (0.02, 0.05)<br>(2.59, 9.08)<br>(NaN, NaN)<br>(1e-14, NaN) | 161.57 |
| 60 | LR size | $\alpha$ | 64.17 | (NaN, NaN) | 159.59 |

|  |  |  |  |  |
| --- | --- | --- | --- | --- |
| | $q = \phi^n / (k^n + \phi^n), n=1$<br>$\tau = \tau_1 + \tau_2 \phi^n / (k^n + \phi^n), n=1$ | $\tau_1$<br>$\tau_2$<br>$k_q$<br>$k_d$ | 3.01 (NaN, NaN)<br>182.21 (NaN, NaN)<br>93.9 (NaN, NaN)<br>0.01 (NaN, NaN) | |
| 61 | LR size<br>$q = \phi^n / (k^n + \phi^n), n=1$<br>$\tau = \tau_1 + \tau_2 \phi^n / (k^n + \phi^n), n=2$ | $\alpha$<br>$\tau_1$<br>$\tau_2$<br>$k_q$<br>$k_d$ | 0.02 (0.01, 0.04)<br>4 (0.19, 65.52)<br>0.28 (NaN, NaN)<br>3.49 (NaN, NaN)<br>0.0002 (1e-14, 100) | 161.7 |
| 62 | LR size<br>$q = \phi^n / (k^n + \phi^n), n=1$<br>$\tau = \tau_1 + \tau_2 \phi^n / (k^n + \phi^n), n=3$ | $\alpha$<br>$\tau_1$<br>$\tau_2$<br>$k_q$<br>$k_d$ | 0.013 (0.009, 0.019)<br>2.03 (NaN, NaN)<br>299.98 (1e-14, NaN)<br>99.99 (NaN, NaN)<br>7.34 (NaN, NaN) | 160.72 |
| 63 | LR size<br>$q = \phi^n / (k^n + \phi^n), n=2$<br>$\tau = \tau_1 + \tau_2 \phi^n / (k^n + \phi^n), n=1$ | $\alpha$<br>$\tau_1$<br>$\tau_2$<br>$k_q$<br>$k_d$ | 0.19 (0.15, 0.25)<br>1.85 (0.15, 20.06)<br>4.46 (1.77, 11.06)<br>0.098 (NaN, NaN)<br>0.0002 (NaN, NaN) | 187.35 |
| 64 | LR size<br>$q = \phi^n / (k^n + \phi^n), n=2$<br>$\tau = \tau_1 + \tau_2 \phi^n / (k^n + \phi^n), n=2$ | $\alpha$<br>$\tau_1$<br>$\tau_2$<br>$k_q$<br>$k_d$ | 0.012 (0.07, 0.02)<br>2.6 (NaN, NaN)<br>2.49 (NaN, NaN)<br>4.15 (0.0005, 99.7)<br>0.0004 (2.98e-13, 99.98) | 158.15 |
| 65 | LR size<br>$q = \phi^n / (k^n + \phi^n), n=2$<br>$\tau = \tau_1 + \tau_2 \phi^n / (k^n + \phi^n), n=3$ | $\alpha$<br>$\tau_1$<br>$\tau_2$<br>$k_q$<br>$k_d$ | 0.012 (0.007, 0.021)<br>2.89 (NaN, NaN)<br>2.26 (NaN, NaN)<br>4.35 (0.004, 97.98)<br>0.016 (NaN, NaN) | 158.17 |
| 66 | LR size<br>$q = \phi^n / (k^n + \phi^n), n=3$<br>$\tau = \tau_1 + \tau_2 \phi^n / (k^n + \phi^n), n=1$ | $\alpha$<br>$\tau_1$<br>$\tau_2$<br>$k_q$<br>$k_d$ | 0.022 (0.02, 0.023)<br>3.11 (1.82, 5.27)<br>2.48 (NaN, NaN)<br>0.57 (0.55, 0.6)<br>0.0001 (NaN, NaN) | 168.17 |
| 67 | LR size<br>$q = \phi^n / (k^n + \phi^n), n=3$<br>$\tau = \tau_1 + \tau_2 \phi^n / (k^n + \phi^n), n=2$ | $\alpha$<br>$\tau_1$<br>$\tau_2$ | 0.012 (0.009, 0.019)<br>5.07 (NaN, NaN)<br>1.3 (NaN, NaN) | 158.16 |

|  |  |  |  |  |  |
| --- | --- | --- | --- | --- | --- |
| | | $k_q$ | 94.86 | (NaN, NaN) | |
| | | $k_d$ | 3.9 | | |
| 68 | LR size<br>$q=\phi^n/(k^n+\phi^n)$, $n=3$<br>$\tau=\tau_1+\tau_2\phi^n/(k^n+\phi^n)$, $n=3$ | $\alpha$ | 0.012 | (0.008, 0.02) | 158.38 |
| | | $\tau_1$ | 3.38 | (NaN, NaN) | |
| | | $\tau_2$ | 1.7 | (NaN, NaN) | |
| | | $k_q$ | 2.03 | (0.11, 26.8) | |
| | | $k_d$ | 0.0008 | (5.7e-14, 99.99) | |
| 69 | LR size<br>$q=\phi^n/(k^n+\phi^n)$, $n=1$<br>$\tau=\tau_1-\tau_2LR$ | $\alpha$ | 0.01 | (0.007, 0.04) | 150.3 |
| | | $\tau_1$ | 28.15 | (0.02, 81.5) | |
| | | $\tau_2$ | 6.57 | (27.84, 28.46) | |
| | | $k$ | 2.72 | (0.017, 81.53) | |
| 70 | LR size<br>$q=\phi^n/(k^n+\phi^n)$, $n=2$<br>$\tau=\tau_1-\tau_2LR$ | $\alpha$ | 0.01 | (0.009, 0.029) | 150.06 |
| | | $\tau_1$ | 28.47 | (0.28, 7.77) | |
| | | $\tau_2$ | 28.47 | (28.18, 28.77) | |
| | | $k$ | 1.53 | (0.28, 7.77) | |
| 71 | LR size<br>$q=\phi^n/(k^n+\phi^n)$, $n=3$<br>$\tau=\tau_1-\tau_2LR$ | $\alpha$ | 0.01 | (0.009, 0.027) | 149.78 |
| | | $\tau_1$ | 29.83 | (29.52, 30.13) | |
| | | $\tau_2$ | 7.18 | (7.08, 7.28) | |
| | | $k$ | 1.22 | (0.48, 3.07) | |
| 72 | LR size<br>$q=\phi^n/(k^n+\phi^n)$, $n=1$<br>$\tau=\tau_1+\tau_2/LR$ | $\alpha$ | 0.016 | (0.01, 0.03) | 154.03 |
| | | $\tau_1$ | 3.37E-05 | (NaN, NaN) | |
| | | $\tau_2$ | 19.9 | (14.04, 27.96) | |
| | | $k$ | 1.99 | (NaN, NaN) | |
| 73 | LR size<br>$q=\phi^n/(k^n+\phi^n)$, $n=2$<br>$\tau=\tau_1+\tau_2\phi/LR$ | $\alpha$ | 0.014 | (0.013, 0.016) | 153.7 |
| | | $\tau_1$ | 3.37E-05 | (4.87e-8, 0.023) | |
| | | $\tau_2$ | 19.88 | (14.18, 27.68) | |
| | | $k$ | 1.99 | (NaN, NaN) | |
| 74 | LR size<br>$q=\phi^n/(k^n+\phi^n)$, $n=3$<br>$\tau=\tau_1+\tau_2/LR$ | $\alpha$ | 0.014 | (0.008, 0.025) | 153.57 |
| | | $\tau_1$ | 3.37E-05 | (3.37E-05, 3.37E-05) | |
| | | $\tau_2$ | 19.87 | (13.98, 28.01) | |
| | | $k$ | 1.56 | (0.3, 7.73) | |
| 75 | LR size<br>$q=\phi^n/(k^n+\phi^n)$, $n=1$<br>$\tau=\tau_1/LR$ | $\alpha$ | 0.015 | (0.01, 0.025) | 152.03 |
| | | $\tau_1$ | 19.9 | (14.04, 27.96) | |
| | | $k$ | 1.99 | (2.28e-11, 2) | |
| 76 | LR size<br>$q=\phi^n/(k^n+\phi^n)$, $n=2$<br>$\tau=\tau_1/LR$ | $\alpha$ | 0.014 | (0.009, 0.022) | 151.7 |
| | | $\tau_1$ | 19.86 | (13.97, 28) | |
| | | $k$ | 1.99 | (1.99, 1.99) | |
| 77 | LR size | $\alpha$ | NaN | (NaN, NaN) | NaN |
| | $q = \phi^n / (k^n + \phi^n), n=1$<br>$\tau = \tau_1 + \tau_2(1 - LR^n / (k^n + LR^n)), n=1$ | $\tau_1$<br>$\tau_2$<br>$k_q$<br>$k_d$ | NaN<br>NaN<br>NaN<br>NaN | (NaN, NaN)<br>(NaN, NaN)<br>(NaN, NaN)<br>(NaN, NaN) | |
| 78 | LR size<br>$q = \phi^n / (k^n + \phi^n), n=1$<br>$\tau = \tau_1 + \tau_2(1 - LR^n / (k^n + LR^n)), n=2$ | $\alpha$<br>$\tau_1$<br>$\tau_2$<br>$k_q$<br>$k_d$ | NaN<br>NaN<br>NaN<br>NaN<br>NaN | (NaN, NaN)<br>(NaN, NaN)<br>(NaN, NaN)<br>(NaN, NaN)<br>(NaN, NaN) | NaN |
| 79 | LR size<br>$q = \phi^n / (k^n + \phi^n), n=1$<br>$\tau = \tau_1 + \tau_2(1 - LR^n / (k^n + LR^n)), n=3$ | $\alpha$<br>$\tau_1$<br>$\tau_2$<br>$k_q$<br>$k_d$ | NaN<br>NaN<br>NaN<br>NaN<br>NaN | (NaN, NaN)<br>(NaN, NaN)<br>(NaN, NaN)<br>(NaN, NaN)<br>(NaN, NaN) | NaN |
| 80 | LR size<br>$q = \phi^n / (k^n + \phi^n), n=2$<br>$\tau = \tau_1 + \tau_2(1 - LR^n / (k^n + LR^n)), n=1$ | $\alpha$<br>$\tau_1$<br>$\tau_2$<br>$k_q$<br>$k_d$ | NaN<br>NaN<br>NaN<br>NaN<br>NaN | (NaN, NaN)<br>(NaN, NaN)<br>(NaN, NaN)<br>(NaN, NaN)<br>(NaN, NaN) | NaN |
| 81 | LR size<br>$q = \phi^n / (k^n + \phi^n), n=2$<br>$\tau = \tau_1 + \tau_2(1 - LR^n / (k^n + LR^n)), n=2$ | $\alpha$<br>$\tau_1$<br>$\tau_2$<br>$k_q$<br>$k_d$ | 0.019<br>0.28<br>299.95<br>1.36<br>0.46 | (0.019, 0.019)<br>(0.28, 0.29)<br>(NaN, NaN)<br>(1.25, 1.27)<br>(0.46, 0.46) | 150.45 |
| 82 | LR size<br>$q = \phi^n / (k^n + \phi^n), n=2$<br>$\tau = \tau_1 + \tau_2(1 - LR^n / (k^n + LR^n)), n=3$ | $\alpha$<br>$\tau_1$<br>$\tau_2$<br>$k_q$<br>$k_d$ | NaN<br>NaN<br>NaN<br>NaN<br>NaN | (NaN, NaN)<br>(NaN, NaN)<br>(NaN, NaN)<br>(NaN, NaN)<br>(NaN, NaN) | |
| 83 | LR size<br>$q = \phi^n / (k^n + \phi^n), n=3$<br>$\tau = \tau_1 + \tau_2(1 - LR^n / (k^n + LR^n)), n=1$ | $\alpha$<br>$\tau_1$<br>$\tau_2$<br>$k_q$<br>$k_d$ | NaN<br>NaN<br>NaN<br>NaN<br>NaN | (NaN, NaN)<br>(NaN, NaN)<br>(NaN, NaN)<br>(NaN, NaN)<br>(NaN, NaN) | |
| 84 | LR size<br>$q = \phi^n / (k^n + \phi^n), n=3$<br>$\tau = \tau_1 + \tau_2(1 - LR^n / (k^n + LR^n)), n=2$ | $\alpha$<br>$\tau_1$<br>$\tau_2$ | 0.12<br>4.62<br>299.99 | (NaN, NaN)<br>(NaN, NaN)<br>(8.43, 300) | 189.37 |
| | | $k_q$ | 0.2 | (NaN, NaN) | |
| | | $k_d$ | 0.27 | (NaN, NaN) | |
| 85 | LR size<br>$q=\phi^n/(k^n+\phi^n)$, $n=3$<br>$\tau=\tau_1+\tau_2(1-LR^n/(k^n+LR^n))$, $n=3$ | $\alpha$ | 0.012 | (0.01, 0.014) | 158.43 |
| | | $\tau_1$ | 5.09 | (4.21, 6.13) | |
| | | $\tau_2$ | 298.79 | (1e-14, NaN) | |
| | | $k_q$ | 99.99 | (1e-14, NaN) | |
| | | $k_d$ | 0.0012 | (1e-14, NaN) | |
| 86 | LR size<br>$\tau=\tau_1+\tau_2\phi/LR$ | $\alpha(1-q)$ | 0.012 | (0.007, 0.019) | 154.37 |
| | | $\tau_1$ | 4.33 | (0.53, 32.4) | |
| | | $\tau_2$ | 5.08 | (0.0002, 299.24) | |
| 87 | LR size<br>$\tau=\tau_1+\tau_2\phi-\tau_3LR$ | $\alpha(1-q)$ | 0.012 | (0.008, 0.02) | 155.53 |
| | | $\tau_1$ | 3.35E-05 | (3.35E-05, 3.35E-05) | |
| | | $\tau_2$ | 26.26 | (13.52m 48.98) | |
| | | $\tau_3$ | 0.25 | (0.0005, 89.24) | |
| 88 | LR size<br>$q=\min(q_0\phi, 1)$<br>$\tau=\tau_1+\tau_2\phi/LR$ | $\alpha$ | 0.012 | (0.008, 0.02) | 156.37 |
| | | $q_0$ | 2.27E-07 | (2.27E-07, 2.27E-07) | |
| | | $\tau_1$ | 4.33 | (0.53, 32.39) | |
| | | $\tau_2$ | 5.08 | (0.0002, 299.24) | |
| 89 | LR size<br>$q=\min(q_0\phi, 1)$<br>$\tau=\tau_1+\tau_2\phi-\tau_3LR$ | $\alpha$ | 0.01 | (0.008, 0.029) | 151.6 |
| | | $q_0$ | 0.21 | (0.002, 1.83) | |
| | | $\tau_1$ | 27.58 | (27.11, 28.05) | |
| | | $\tau_2$ | 3.27 | (2.61, 4.10) | |
| | | $\tau_3$ | 7.01 | (6.86, 7.15) | |
| 90 | LR size<br>$q=\phi^n/(k^n+\phi^n)$, $n=1$<br>$\tau=\tau_1+\tau_2\phi/LR$ | $\alpha$ | 0.014 | (0.009, 0.023) | 156.99 |
| | | $k$ | 1.99 | (NaN, NaN) | |
| | | $\tau_1$ | 5.05 | (2.9, 8.73) | |
| | | $\tau_2$ | 0.19 | (NaN, NaN) | |
| 91 | LR size<br>$q=\phi^n/(k^n+\phi^n)$, $n=2$<br>$\tau=\tau_1+\tau_2\phi/LR$ | $\alpha$ | 0.013 | (0.011, 0.01) | 156.54 |
| | | $k$ | 1.99 | (1.99, 1.99) | |
| | | $\tau_1$ | 4.63 | (3.94, 5.44) | |
| | | $\tau_2$ | 3.01 | (0.53, 16.35) | |
| 92 | LR size<br>$q=\phi^n/(k^n+\phi^n)$, $n=3$<br>$\tau=\tau_1+\tau_2\phi/LR$ | $\alpha$ | 0.012 | (0.008, 0.02) | 156.35 |
| | | $k$ | 1.99 | (1.99, 1.99) | |
| | | $\tau_1$ | 4.64 | (0.87, 23.47) | |
| | | $\tau_2$ | 3.2 | (3.98e-6, 299.96) | |
| 93 | LR size<br>$q=\phi^n/(k^n+\phi^n)$, $n=1$<br>$\tau=\tau_1+\tau_2\phi-\tau_3LR$ | $\alpha$ | 0.017 | (0.011, 0.02) | 151.72 |
| | | $k$ | 1.999 | (6.59e-11, 2) | |
| | | $\tau_1$ | 28.34 | (27.44, 29.26) | |
| | | $\tau_2$ | 2.44 | (1.96, 3.05) | |
| | | $\tau_3$ | 7.04 | (6.8, 7.27) | |
| 94 | LR size<br>$q=\phi^n/(k^n+\phi^n)$, $n=2$<br>$\tau=\tau_1+\tau_2\phi-\tau_3LR$ | $\alpha$ | 0.015 | (0.012, 0.02) | 151.41 |
| | | $k$ | 1.91 | (0.005, 1.99) | |
| | | $\tau_1$ | 29.37 | (29.06, 29.6) | |
| | | $\tau_2$ | 4.14 | (3.7, 4.63) | |
| | | $\tau_3$ | 7.64 | (7.55, 7.74) | |
| 95 | LR size<br>$q=\phi^n/(k^n+\phi^n)$, $n=3$<br>$\tau=\tau_1+\tau_2\phi-\tau_3LR$ | $\alpha$ | 0.015 | (0.012, 0.02) | 151.24 |
| | | $k$ | 1.34 | (0.16, 1.95) | |
| | | $\tau_1$ | 28.5 | (28.06, 28.94) | |
| | | $\tau_2$ | 2.92 | (2.42, 3.52) | |
| | | $\tau_3$ | 7.17 | (6.98, 7.36) | |
| 96 | LR size<br>$\tau=\tau_1+\tau_2\phi-\tau_3/LR+\tau_4\phi/LR$ | $\alpha(1-q)$ | 0.013 | (0.009, 0.02) | 151.75 |
| | | $\tau_1$ | 27.87 | (27.31, 28.45) | |
| | | $\tau_2$ | 2.93 | (2.33, 3.68) | |
| | | $\tau_3$ | 7.03 | (6.88, 7.17) | |
| | | $\tau_4$ | 0.3 | (1.38e-8, 299.98) | |
| 97 | LR size<br>$\tau=\tau_1+\tau_2\phi+\tau_3/LR+\tau_4\phi/LR$ | $\alpha(1-q)$ | 0.013 | (0.009, 0.02) | 155.79 |
| | | $\tau_1$ | 3.37E-05 | (3.37E-05, 3.37E-05) | |
| | | $\tau_2$ | 3.37E-05 | (3.37E-05, 3.37E-05) | |
| | | $\tau_3$ | 19.77 | (13.86, 27.96) | |
| | | $\tau_4$ | 0.2 | (0.2, 0.2) | |
| 98 | LR size<br>$q=\min(q_0\phi, 1)$<br>$\tau=\tau_1+\tau_2\phi-\tau_3/LR+\tau_4\phi/LR$ | $\alpha$ | 0.015 | (0.014, 0.016) | 153.88 |
| | | $q_0$ | 0.18 | (0.009, 1.4) | |
| | | $\tau_1$ | 26.52 | (26.18, 26.86) | |
| | | $\tau_2$ | 0.004 | (NaN, NaN) | |
| | | $\tau_3$ | 6.82 | (6.68, 6.96) | |
| | | $\tau_4$ | 14.29 | (13.35, 15.3) | |
| 99 | LR size<br>$q=\phi^n/(k^n+\phi^n)$, $n=1$<br>$\tau=\tau_1+\tau_2\phi-\tau_3/LR+\tau_4\phi/LR$ | $\alpha$ | 0.017 | (0.011, 0.026) | 154 |
| | | $k$ | 1.99 | (1.05e-14, 2) | |
| | | $\tau_1$ | 27.39 | (26.83, 27.96) | |
| | | $\tau_2$ | 0.003 | (NaN, NaN) | |
| | | $\tau_3$ | 6.92 | (6.74, 7.1) | |
| | | $\tau_4$ | 11.96 | (11.25, 12.71) | |
| 100 | LR size<br>$q=\phi^n/(k^n+\phi^n)$, $n=2$<br>$\tau=\tau_1+\tau_2\phi-\tau_3/LR+\tau_4\phi/LR$ | $\alpha$ | 0.015 | (0.009, 0.02) | 153.48 |
| | | $k$ | 1.82 | (1.67e-5, 1.99) | |
| | | $\tau_1$ | 28.45 | (27.59, 29.33) | |
| | | $\tau_2$ | 2.63 | (2.11, 3.27) | |
| | | $\tau_3$ | 7.25 | (7.03, 7.49) | |
| | | $\tau_4$ | 2.91 | (1.68, 5.04) | |
| 101 | LR size<br>$q=\phi^n/(k^n+\phi^n)$, $n=3$<br>$\tau=\tau_1+\tau_2\phi-\tau_3/LR+\tau_4\phi/LR$ | $\alpha$ | 0.015 | (0.014, 0.02) | 153.48 |
| | | $k$ | 1.36 | (0.31, 1.91) | |
| | | $\tau_1$ | 26.68 | (26.29, 27.07) | |
| | | $\tau_2$ | 0.02 | (2.45e-6, 112.89) | |
| | | $\tau_3$ | 6.58 | (6.44, 6.27) | |
| | | $\tau_4$ | 9.05 | (7.08, 11.53) | |
| 102 | LR size<br>$q=\min(q_0\phi, 1)$<br>$\tau=\tau_1+\tau_2\phi+\tau_3/LR+\tau_4\phi/LR$ | $\alpha$ | 0.013 | (0.008, 0.02) | 157.79 |
| | | $q_0$ | 2.25E-07 | (NaN, NaN) | |
| | | $\tau_1$ | 3.35E-05 | (1e-14, 300) | |
| | | $\tau_2$ | 3.37E-05 | (1e-14, 300) | |
| | | $\tau_3$ | 19.41 | (10.2, 35.96) | |
| | | $\tau_4$ | 1.04 | (4.25e-12, 300) | |
| 103 | LR size<br>$q=\phi^n/(k^n+\phi^n)$, $n=1$<br>$\tau=\tau_1+\tau_2\phi+\tau_3/LR+\tau_4\phi/LR$ | $\alpha$ | 0.014 | (0.01, 0.02) | 158.04 |
| | | $k$ | 1.99 | (NaN, NaN) | |
| | | $\tau_1$ | 3.35E-05 | (3.35E-05, 3.35E-05) | |
| | | $\tau_2$ | 3.37E-05 | (3.35E-05, 3.35E-05) | |
| | | $\tau_3$ | 19.9 | (14.04, 27.96) | |
| | | $\tau_4$ | 3.35E-05 | (3.35E-05, 3.35E-05) | |
| 104 | LR size<br>$q=\phi^n/(k^n+\phi^n)$, $n=2$<br>$\tau=\tau_1+\tau_2\phi+\tau_3/LR+\tau_4\phi/LR$ | $\alpha$ | 0.014 | (0.009, 0.02) | 157.7 |
| | | $k$ | 1.99 | (1.99, 1.99) | |
| | | $\tau_1$ | 3.35E-05 | (3.35E-05, 3.35E-05) | |
| | | $\tau_2$ | 3.37E-05 | (3.35E-05, 3.35E-05) | |
| | | $\tau_3$ | 19.86 | (13.97, 28.01) | |
| | | $\tau_4$ | 3.37E-05 | (3.35E-05, 3.35E-05) | |
| 105 | LR size<br>$q=\phi^n/(k^n+\phi^n)$, $n=3$<br>$\tau=\tau_1+\tau_2\phi+\tau_3/LR+\tau_4\phi/LR$ | $\alpha$ | 0.014 | (0.009, 0.02) | 157.57 |
| | | $k$ | 1.57 | (0.01, 1.99) | |
| | | $\tau_1$ | 3.35E-05 | (3.35E-05, 3.35E-05) | |
| | | $\tau_2$ | 3.37E-05 | (3.35E-05, 3.35E-05) | |
| | | $\tau_3$ | 19.98 | (13.98, 28.01) | |
| | | $\tau_4$ | 3.37E-05 | (3.35E-05, 3.35E-05) | |
| 106 | LR size<br>$\tau=\tau_1+\tau_2\phi-\tau_3LR+\tau_4\phi^*LR$ | $\alpha(1-q)$ | 0.013 | (0.01, 0.017) | 151.18 |
| | | $\tau_1$ | 31.68 | (25.06, 39.78) | |
| | | $\tau_2$ | 0.03 | (1.07e-14, 300) | |
| | | $\tau_3$ | 8.76 | (6.89, 11.11) | |
| | | $\tau_4$ | 1.86 | (-0.87, 4.61) | |
| 107 | LR size | $\alpha$ | 0.015 | (0.011, 0.02) | 153.5 |
| | $q = \min(q_0\phi, 1)$<br>$\tau = \tau_1 + \tau_2\phi - \tau_3LR + \tau_4\phi * LR$ | $q_0$<br>$\tau_1$<br>$\tau_2$<br>$\tau_3$<br>$\tau_4$ | 0.21<br>29.68<br>4.25<br>7.88<br>0.16 | (0.13, 0.3)<br>(29.22, 0.3)<br>(3.6, 5.02)<br>(7.76, 8.01)<br>(0.003, 0.33) | |
| 108 | LR size<br>$q = \phi^n / (k^n + \phi^n)$, $n=1$<br>$\tau = \tau_1 + \tau_2\phi - \tau_3LR + \tau_4\phi * LR$ | $\alpha$<br>$k$<br>$\tau_1$<br>$\tau_2$<br>$\tau_3$<br>$\tau_4$ | 0.017<br>1.99<br>29.73<br>2.39<br>7.59<br>0.26 | (0.016, 0.018)<br>(7.023-13, 2)<br>(29.43, 30.03)<br>(1.84, 3.1)<br>7.5, 7.68)<br>(0.076, 0.46) | 153.58 |
| 109 | LR size<br>$q = \phi^n / (k^n + \phi^n)$, $n=2$<br>$\tau = \tau_1 + \tau_2\phi - \tau_3LR + \tau_4\phi * LR$ | $\alpha$<br>$k$<br>$\tau_1$<br>$\tau_2$<br>$\tau_3$<br>$\tau_4$ | 0.015<br>1.98<br>27.61<br>3.2<br>6.91<br>-0.11 | (0.01, 0.02)<br>(NaN, NaN)<br>(26.96, 28.27)<br>(NaN, NaN)<br>(6.74, 7.08)<br>(-0.8, 0.56) | 153.52 |
| 110 | LR size<br>$q = \phi^n / (k^n + \phi^n)$, $n=3$<br>$\tau = \tau_1 + \tau_2\phi - \tau_3LR + \tau_4\phi * LR$ | $\alpha$<br>$k$<br>$\tau_1$<br>$\tau_2$<br>$\tau_3$<br>$\tau_4$ | 0.015<br>1.47<br>23.97<br>9.85<br>6.06<br>-1.71 | (0.01, 0.02)<br>(0.08, 1.98)<br>(23.44, 24.52)<br>(9.13, 10.64)<br>(5.92, 6.2)<br>(-19.2, -1.5) | 153.91 |

| $\Delta AIC$ |
| --- |
| 0 |
| 2.01 |
| 1.86 |
| 2.01 |
| 3.86 |
| 3.94 |
| -5.14 |
| -3.11 |
| -3.15 |
| -3.14 |
| -1.15 |
| -1.13 |

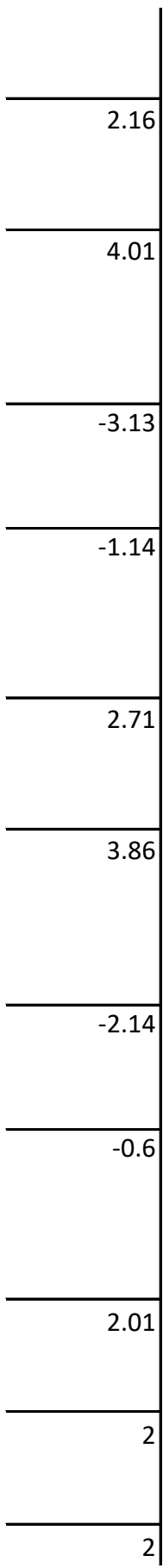

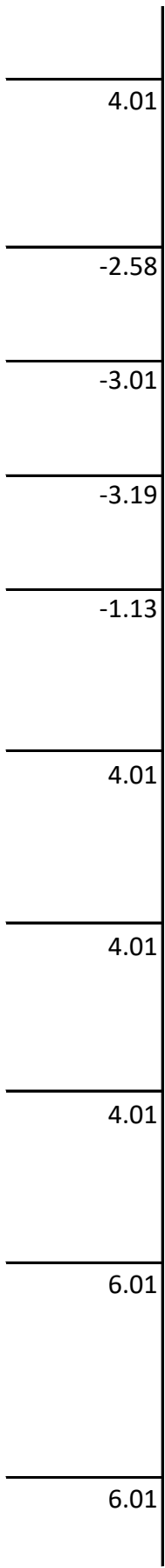

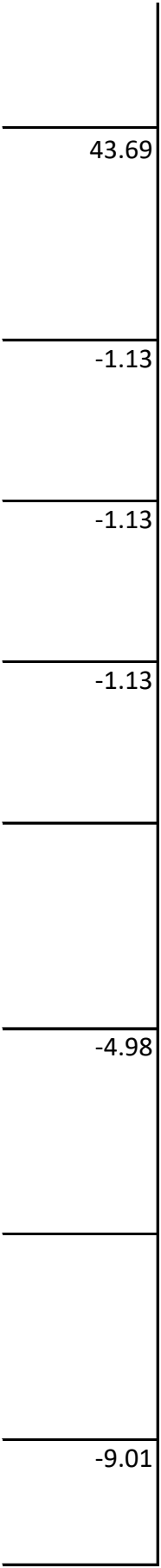

-5.77

-7.35

-5.76

-5.8

-16.38

-2.39

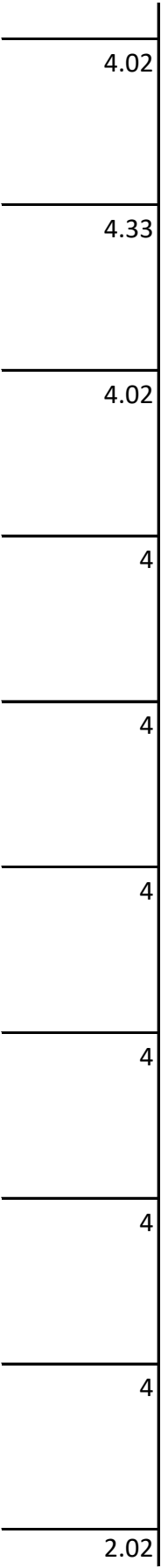

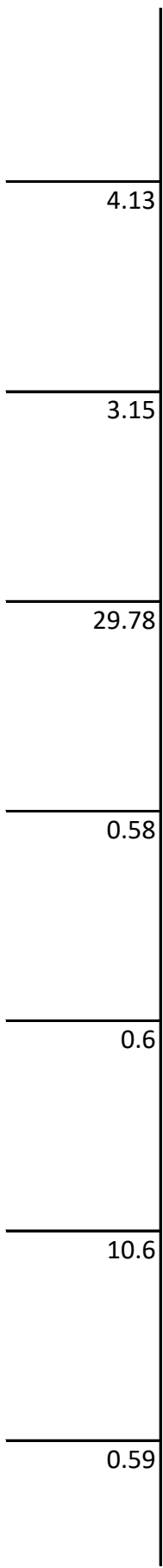

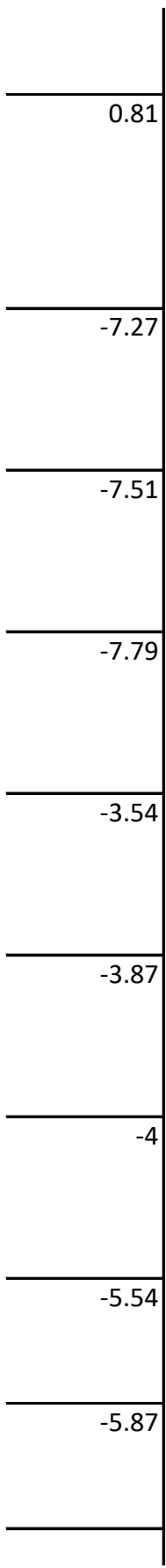

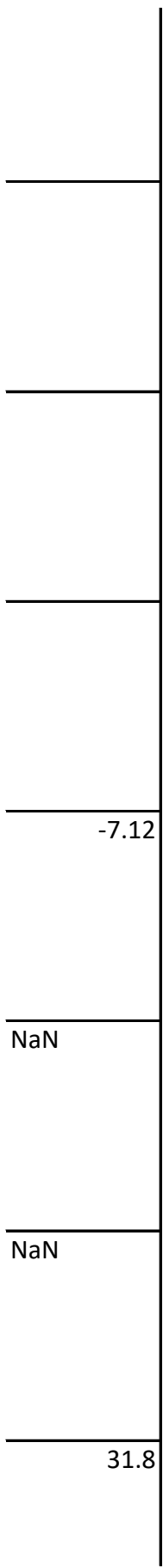

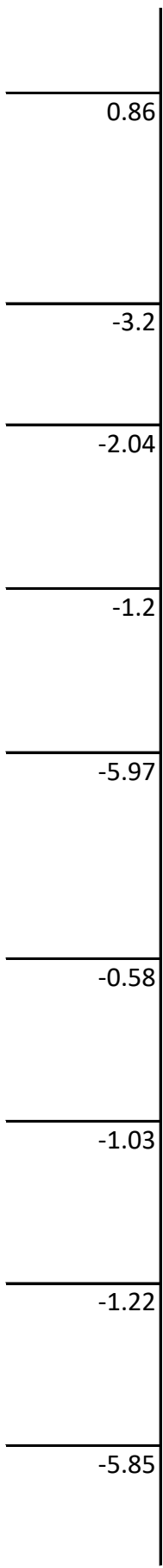

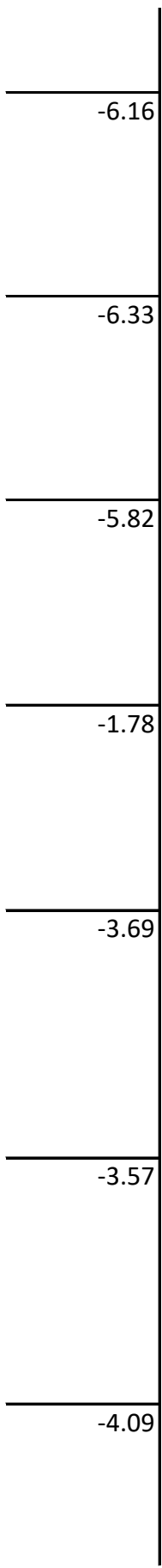

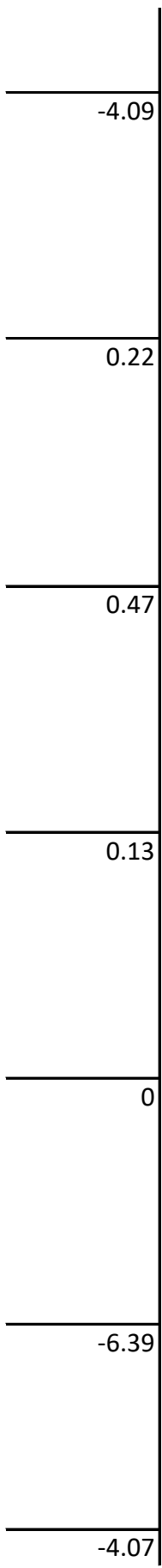

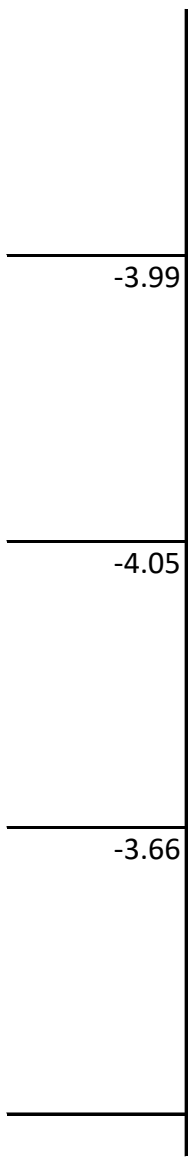
